# Water oxidation-driven histidine dioxidation enables probe-free proximity labeling

**DOI:** 10.64898/2026.06.10.731337

**Authors:** Chaiheon Lee, Jeong Kyeong Lee, Chang-Mo Yoo, Byeong Gyu Kim, Gwangsu Yoon, Jiwoong Kang, Seungjin Na, Hyun-Woo Rhee, Tae-Hyuk Kwon

## Abstract

Photocatalytic proximity labeling (photo-PL) has emerged as a powerful tool for spatial proteomics in subcellular compartments. However, many photo-PL toolboxes rely on singlet oxygen (^1^O_2_) to generate highly unstable endoperoxide intermediates that must be trapped immediately by high concentrations of exogenous probes. This constraint can bias spatial proteome coverage, particularly in dynamic and heterogeneous compartments such as endosomes and exosomes, where probe accessibility is intrinsically nonuniform. Here, we develop IDM, an organic photocatalyst that generates hydroxyl (•OH) and superoxide (O_2_ ^•−^) radicals via water oxidation instead of ^1^O , enabling a synergistic dual-radical mechanism for probe-free proximal protein mapping in live cells (PF-Map). The resulting radicals dioxidize proximal histidine residues into a persistent dioxidized state (His-2O) that is thermodynamically stabilized as lactam tautomers, which remain electrophilic and chemically addressable after cell lysis. By decoupling histidine oxidation from live□cell probe capture, PF□Map can minimize spatial bias arising from heterogeneous probe distribution. Applying PF□Map to intracellular vesicle trafficking, we find that both PF□Map and a probe□dependent workflow (PD□Map) robustly identify exosome markers, whereas PF□Map additionally reveals a hidden vesicle trafficking–related subproteome that PD-Map underestimated. Together, we establish a minimally biased photocatalytic strategy for spatial protein mapping in complex biological systems.

**Graphical abstract:** 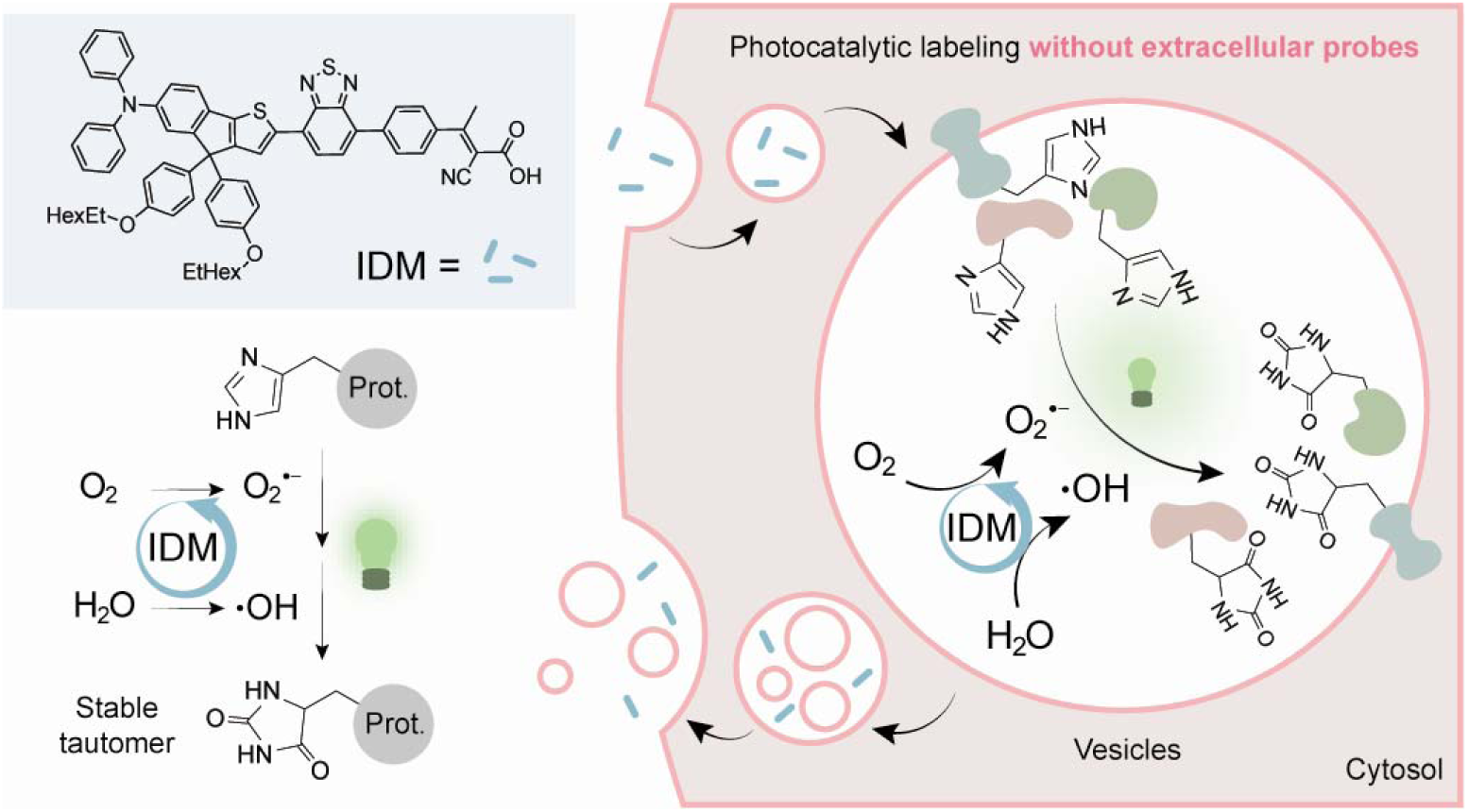

## Introduction

Proximity labeling (PL) has emerged as a powerful approach for spatially resolved proteomics^1–3^. Recently, photocatalyst-driven PL (photo-PL) has been widely employed due to its high spatiotemporal resolution^4–7^. Many photo-PL toolboxes rely exclusively on singlet oxygen (^1^O_2_) generation to oxidize target amino acids, such as histidines^8–11^. While effective in certain environments, this ^1^O_2_-driven mechanism suffers from a chemical limitation: the resulting endoperoxide intermediates are extremely transient and highly unstable^11–13^. To prevent the loss of these labeled targets, conventional photo□PL relies on immediate trapping by high concentrations of exogenous probes^6, 14^, coupling oxidative labeling and probe capture in a single step. This probe dependence can introduce spatial bias and incomplete proteome coverage due to intracellular probe accessibility, particularly when investigating dynamic cellular processes such as vesicle trafficking (e.g., endosomes and exosomes).

Dynamic vesicular organelles, including endosomes and exosomes, play a central role in the uptake, processing, and secretion of extracellular cargos^15, 16^. Because of the complex interactions among vesicles during trafficking, individual vesicles have different biomolecular compositions across cellular states and microenvironments^17–19^, which can create substantial heterogeneity in probe access and local capture efficiency during photo□PL. Therefore, the proteomic landscape across diverse vesicle trafficking processes has remained difficult to interrogate comprehensively using photo-PL. In particular, the limited accessibility of chemical probes raises the possibility that spatial proteomics preferentially reflects probe-accessible vesicle subsets rather than the full vesicle proteome. These challenges highlight the need for protein labeling approaches that do not rely on exogenous small-molecule probes to map the proteome across vesicle trafficking pathways.

Herein, we introduce a water oxidation-driven photocatalytic reaction mode to enable “probe-free” proximal protein mapping in live cells (PF-Map), in which proximal histidines are oxidized without an exogenous capture probe and subsequently tagged after lysis. Instead of the ^1^O_2_-mediated protein labeling that generates unstable intermediates (**Fig. 1a**), we employ water and oxygen as ubiquitous labeling reagents to produce stable dioxidized histidines (His-2O) through a dual-radical mechanism. Although a recent study has used oxidative radical species to label proteins at lipid droplets, it still relies on exogenous probes to simultaneously capture labeled proteins^20^. For PF-Map, we develop an indenothiophene-based organic photocatalyst, IDM, specifically engineered to bypass ^1^O_2_ generation and instead redirect the formation of reactive radicals via water oxidation. Unlike conventional photocatalysts, IDM exhibits a sufficiently positive excited-state redox potential to oxidize water molecules and subsequently generate hydroxyl radicals (•OH) under green light irradiation. Combined GC-MS analysis and density functional theory (DFT) calculations support a cooperative mechanism in which superoxide (O_2_^•−^) and hydroxyl radicals (•OH) act sequentially: nucleophilic attack by O_2_^•−^ initiates the reaction, followed by the addition of water-derived •OH (**Fig. 1b**). This unique synergistic mechanism transforms proximal histidines into a thermodynamically stable double-lactam tautomer (His-2O), thereby converting a transient trapping event into a persistent and chemically addressable oxidation state.

**Fig. 1.**
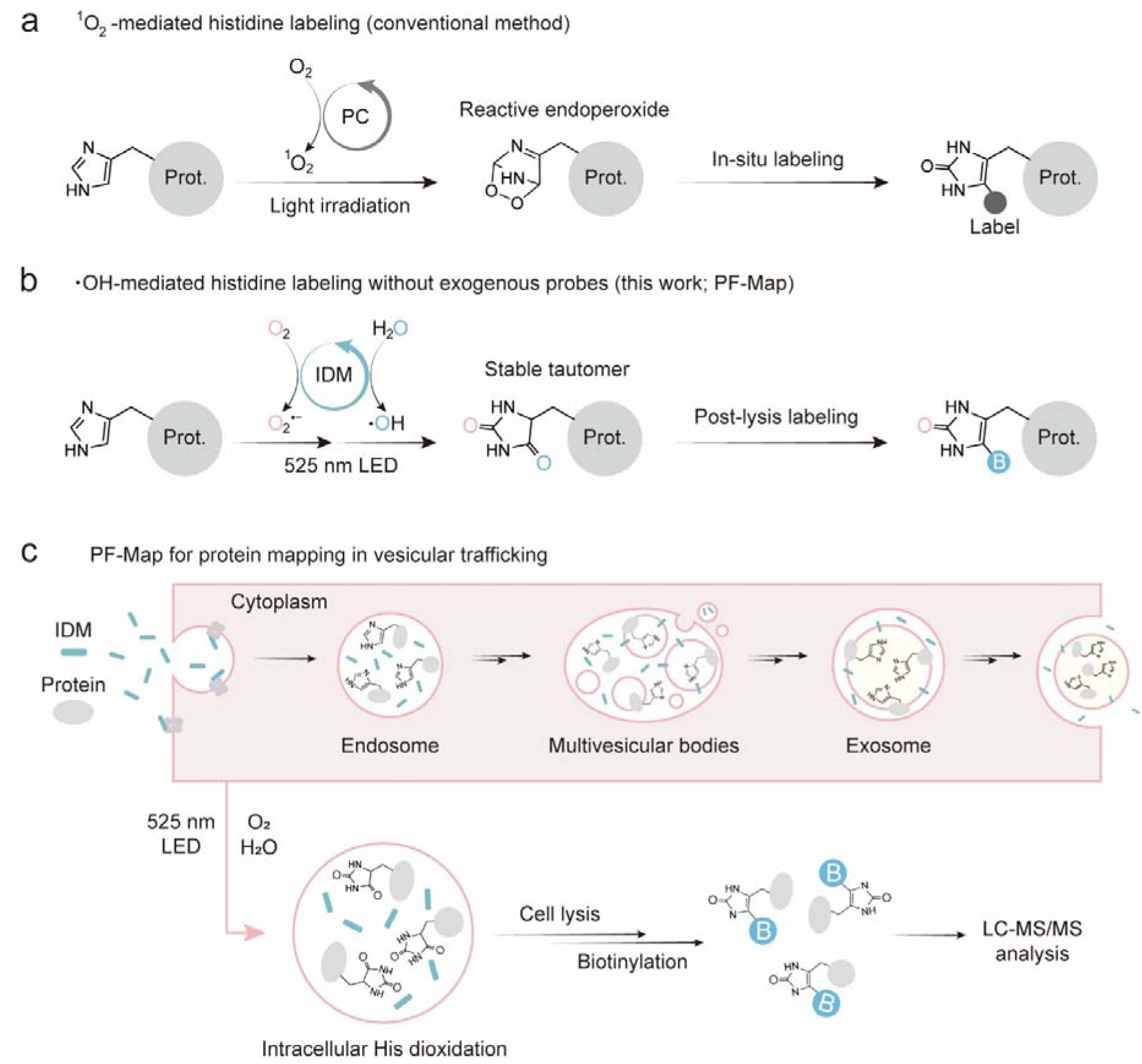
Schematic illustrations of protein mapping for vesicle trafficking processes. **a** mechanistic scheme of conventional histidine labeling with photocatalysts generating ^1^O_2_. **b.** Scheme of hydroxyl radical-mediated histidine dioxidation by IDM photocatalysis. Dioxidized histidine can be stabilized via tautomerization in cell environments, enabling further biotin tagging after cell lysis (PF-Map). **c.** Scheme of PF-Map for protein mapping in vesicle trafficking. IDM reversibly conjugates to random membrane proteins, leading to involvement in vesicle trafficking. Under green light irradiation, IDM in vesicles induces histidine dioxidation, and the labeled proteins are subsequently biotinylated after cell lysis.

Given the stability of His-2O generated by IDM, we employ this persistent oxidation state to establish PF-Map for post-lysis capture of proximity-labeled proteomes in intracellular vesicle trafficking (**Fig. 1c**). Using a uniform photocatalyst, we leverage complementary, orthogonal proteomics workflows—a probe-dependent and a probe-free method—to identify intracellular vesicular proteomes. Notably, PF□Map expands vesicle trafficking coverage by capturing established vesicle markers and a vesicle trafficking–related subproteome of 125 proteins (e.g., Rab5, Rab11, SEC31A) that are underrepresented in the probe□dependent workflow. Collectively, this work provides a minimally biased, probe-free spatial proteomics toolbox that is potentially applicable to dynamic subcellular compartments and in vivo systems where the accessibility of exogenous chemical probes is limited.

## Results

### IDM photocatalysis generates hydroxyl radicals and oxidizes histidine

To guide the rational design of photocatalysts capable of generating •OH from water, we synthesized a model series of reported photocatalysts with different redox potentials (TP-1, TP-2, and TP-3) following a literature procedure (**Supplementary Fig. S1a**)^21^. We then examined •OH production using hydroxyphenyl fluorescein (HPF), a •OH-selective fluorescent probe^22^. Notably, TP-3 exhibited efficient •OH generation, whereas TP-1 and TP-2 produced little •OH, consistent with the oxidizing ability of the photocatalysts’ excited states (**Supplementary Fig. S1b-d**). Because TP-3 has more positive excited-state reduction potentials (E*^(0/−)^=1.57 V vs. NHE at pH 7.4) than TP-1 (E*^(0/−)^=0.86 V) and TP-2 (E*^(0/−)^=0.59 V), we hypothesized that a sufficiently positive redox potential, and the resulting ability to drive photocatalytic water oxidation, is a key determinant for designing •OH-generating photocatalysts. However, carbazole-conjugated triphenylamine (TPA) is hydrophobic and large, leading to aggregation in physiological environments.

Therefore, we developed a more biocompatible photocatalyst, IDM—an indenothiophene-based photocatalyst featuring a donor-acceptor architecture with diphenylamine as the electron donor and benzothiadiazole as the electron acceptor (**Fig. 2a**). The indenothiophene-fused diphenylamine enhances molecular planarity and conjugation, broadening the absorption spectrum^23^. The incorporation of the strong benzothiadiazole acceptor shifts the IDM potentials toward more positive values, further stabilized by the enhanced delocalization afforded by indenothiophene. This results in a positive shift in the ground- and excited-state reduction potential (E^(0/−)^ and E*^(0/−)^) of IDM, making it a strong oxidant for the oxidation of H_2_O. Additionally, strategic modifications with aryl groups and negatively charged cyanoacrylic acid were incorporated to improve solubility. We analyzed the photophysical properties of IDM (**Supplementary Fig. S2**). IDM demonstrated strong absorbance in aqueous conditions within the green region (λ=513 nm, ε=17,350 M^-1^·cm^-1^). While many photocatalytic PL platforms have suffered from undesired photodamage and off-target labeling due to the use of UV-blue light, the absorption range of IDM enables its application with minimal perturbation. At the same time, IDM exhibits redox potentials that are sufficiently positive to oxidize water, a property attributed to its indenothiophene scaffold (**Supplementary Fig. S3**). Cyclic voltammetry revealed the ground- and excited-state reduction potentials of IDM^+^ and IDM* (E^(+/0)^=1.40 V and E*^(0/−)^=1.82 V vs. NHE at pH 7.4), which exceed the potential required for water oxidation to generate hydrogen peroxide (H_2_O_2_; E_ox_=1.34 V; **Fig. 2b(1)**). Furthermore, the oxidation potentials of IDM* and IDM^−^ (E*^(+/0)^=−0.72 V and E^(0/−)^=−0.30 V) are thermodynamically favorable for electron transfer to both oxygen (E_red_=−0.33 V; **Fig. 2b(2)**) and H_2_O_2_ (E_red_=0.38 V; **Fig. 2b(3)**), enabling the generation of O_2_^•−^ and •OH, respectively^24,25,26^.

**Fig. 2.**
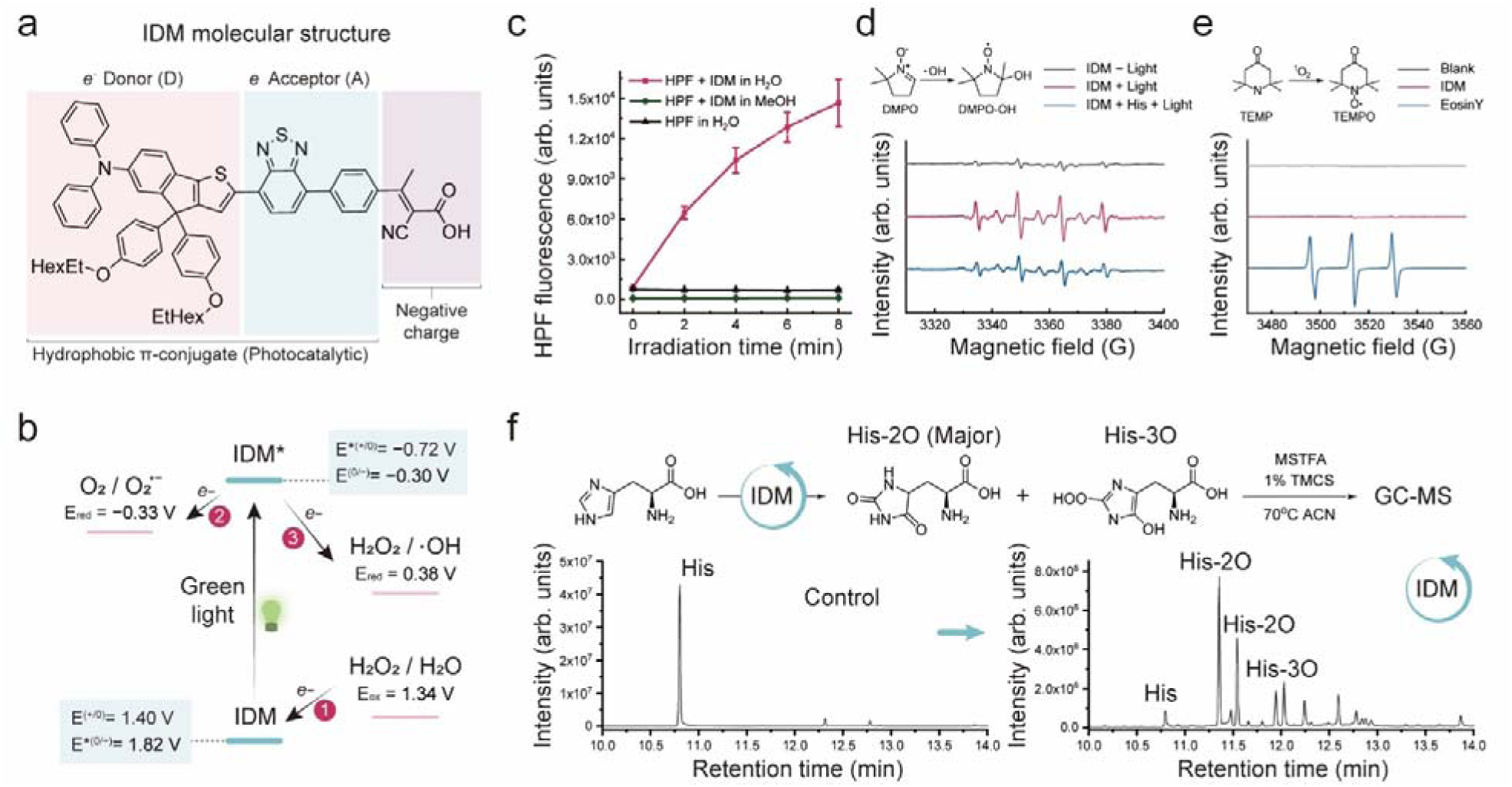
ROS generation by IDM photocatalysis for His dioxidation. **a** Molecular structure of IDM. **b** Energy diagram of electron transfer for ROS generation. The energy levels were described vs. NHE (at pH 7.4) **c** •OH generation assay with HPF and DMPO. In the HPF assay, •OH generation was measured by fluorescence changes of HPF solutions with irradiation energy (green LED, λ_max_ = 525 nm, 16.6 mW·cm^−2^). **d** In EPR spectroscopy, a spectrum of the spin adduct DMPO-OH was observed after IDM photocatalysis under aqueous conditions ([IDM] = 1 mM, [His] = 7 mM, [DMPO] = 100 mM, and green LED, 0.1 J·cm^−2^). **e** EPR spectra of ^1^O_2_ spin adducts (TEMPO) after IDM and Eosin Y photoactivation ([IDM] = [EosinY] = 0.2 mM, [TEMP] = 100 mM, and green LED 1.5 J·cm^−2^). **f** Chromatogram of GC-MS before and after IDM photocatalysis for histidine dioxidation. Reaction conditions: [IDM] = 0.5 mM, [His] = 5 mM in 500 µL aqueous solution, green light irradiation: 120 J·cm^−2^ (16.6 mW·cm^−2^ for 2 hours). After photocatalysis, the dioxidized histidine was further silylated using MSTFA for gas chromatography analysis. Data are presented as mean ± s.d. (*n* = 3 independent experiments). Source data are provided as a Source Data file.

Given the redox potential of excited IDM, IDM photocatalysis could drive both water oxidation and oxygen reduction. To examine redox reactions, we analyzed the generation of reactive oxygen species (ROS)—including O_2_^•−^, ^1^O_2_, H_2_O_2_, and •OH—using specific assays in aqueous media (Fig. 2c,d; **Supplementary Figs. S4–6**). H_2_O_2_ generation was measured via the *N,N*-diethyl-p-phenylenediamine (DPD)/horseradish peroxidase (HRP) system^27^. Green light activation (λ=525 nm) of IDM produced significant H_2_O_2_ levels (**Supplementary Fig. S4**), consistent with a two-electron water oxidation mechanism. Parallel experiments using HPF^22^, •OH-specific fluorescent probe, revealed a sharp increase in fluorescence upon IDM photoactivation (**Fig. 2c**), confirming substantial •OH generation. To verify •OH and ^1^O_2_ generation, we performed electron paramagnetic resonance (EPR) spectroscopy using *5,5*-dimethyl-1-pyrroline *N*-oxide (DMPO) and *2,2,6,6*-tetramethylpiperidine (TEMP) as spin traps^28, 29^. The EPR spectrum of IDM under photoactivation with DMPO (•OH-specific trap) showed distinct signals corresponding to the DMPO-OH adduct, confirming the production of •OH (**Fig. 2d**). Additionally, the DMPO-OH signal decreased in the presence of histidine, suggesting that the generated OH •reacts with histidine. In contrast, TEMP (^1^O_2_-specific trap) yielded no detectable signals under IDM photoactivation (**Fig. 2e**). The absence of ^1^O_2_ generation by IDM was further corroborated by the 9,10-anthracenediyl-bis(methylene)dimalonic acid (ABDA) assay (**Supplementary Fig. S5**)^30^. We calculated the singlet and triplet energy levels of IDM to rationalize the rare ^1^O_2_ generation of IDM photocatalysis. The result indicates that intersystem crossing of the excited IDM is restricted by a wide S_1_-T_1_ gap, and the low-lying T_1_ energy level is thermodynamically insufficient for excitation of ^1^O_2_. (**Supplementary Fig. S6**). To measure the O_2_ ^•−^ generation, we conducted EPR with DMPO in dimethyl sulfoxide (DMSO), which quenches •OH in a water-deficient environment. Photoactivated IDM generated a DMPO-OOH adduct signal (**Supplementary Fig. S7**), confirming O_2_ ^•−^ production via oxygen reduction. Together, these results demonstrate that IDM photocatalysis in aqueous conditions generates O_2_^•−^, H_2_O_2_, and •OH—but not ^1^O_2_.

Next, we examined whether IDM photocatalysis could oxidize histidine residues through a ^1^O_2_-independent mechanism. An aqueous histidine solution containing IDM was irradiated with green light for 2 hours. The mixture was then derivatized with *N*-trimethylsilyl-*N*-methyl trifluoroacetamide (MSTFA) and 1% trimethylchlorosilane (TMCS) to enhance volatility and thermal stability for GC-MS analysis (**Fig. 2f, Supplementary Fig. S8, S9**)^31^. The GC chromatogram revealed a sharp decrease in the histidine peak, accompanied by distinct chromatographic peaks corresponding to oxidized species. To confirm histidine di- and trioxidation via IDM photocatalysis, we further analyzed the reaction products using high-resolution mass spectrometry (HRMS). The results exhibited prominent dioxidation (+32 Da) and trioxidation (+48 Da) products (His-2O and His-3O; **Supplementary Fig. S10**). This confirms that IDM photocatalysis oxidizes histidine by generating reactive oxygen radicals rather than ^1^O_2_, implying that IDM photocatalysis could trigger histidine dioxidation with a distinct mechanism independent of the ^1^O_2_ pathway.

### Mechanism study of histidine dioxidation by IDM photocatalysis

To investigate the mechanism of histidine oxidation by IDM photocatalysis, we performed GC-MS analysis of histidine oxidation under hypoxic conditions (**Fig. 3a** and **Supplementary Fig. S11**). Although histidine dioxidation has been primarily linked to ^1^O_2_ in prior studies^9–12^, a distinct oxidative pathway operates here. Notably, no reaction occurred under hypoxic conditions, suggesting that oxygen molecules are essential for histidine oxidation. Considering O_2_^•−^ generation via IDM-mediated oxygen reduction, O_2_^•−^ can attack the electrophilic ε-carbon of the imidazole ring, forming a peroxyl intermediate^32, 33^. Together, these observations support that nucleophilic attack by O_2_^•−^ constitutes the initial step in histidine oxidation. Next, we performed IDM photocatalysis in H_2_^18^O and analyzed products via GC-MS to determine the involvement of •OH from water (**Fig. 3b** and **Supplementary Fig. S12**). Notably, His-2O and His-3O peaks exhibited a +2 Da mass shift, indicating that one oxygen atom in each product derives from water. These data support a model in which both O_2_^•−^ from oxygen and •OH from water participate in histidine oxidation, independent of ^1^O_2_-mediated pathways. Consequently, we propose a plausible dual-radical mechanism (**Fig. 3c**). Once O_2_^•−^ initiates oxidation by attacking the ε-carbon of histidine’s imidazole ring, subsequent •OH addition to the γ- and δ-carbons is possible. Given that the hydroxylation at the γ position breaks the aromaticity of the imidazole ring, hydroxylation to the δ-carbon could be favorable (His-3O-δ). The preferential δ□hydroxylation is supported by the mass spectrum of trioxidation products (His-3O) exhibiting penta-silylated m/z values (**Supplementary Fig. S9c**), since γ□trioxidation product (His□3O□γ) would be expected to be silylated at four different positions.

**Fig. 3.**
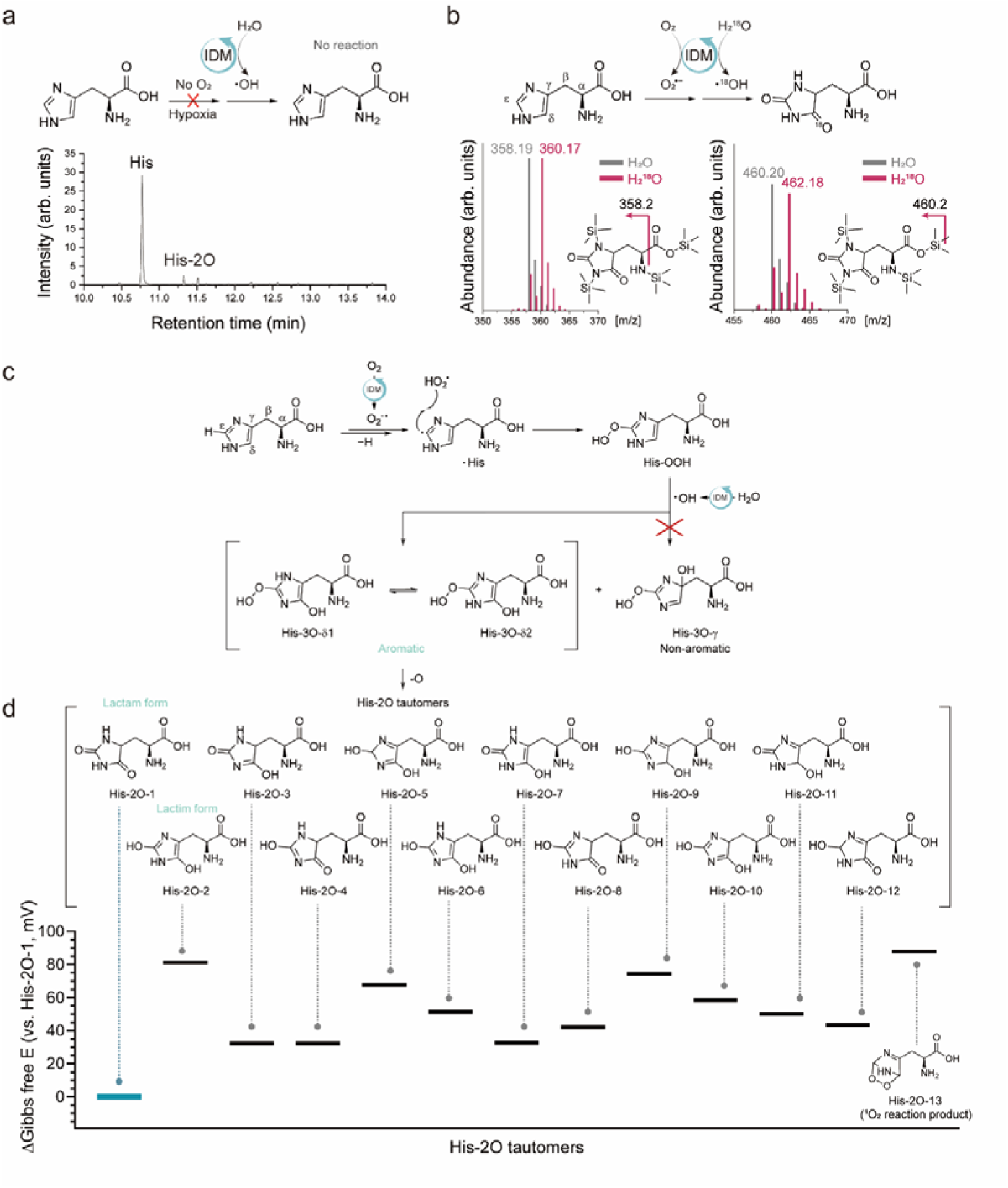
His dioxidation mechanism. **a** GC chromatogram of histidine oxidation in hypoxia. The reaction mixture was further silylated by MSTFA for gas chromatography. The mass spectra are provided in the Supplementary Information. [IDM] = 0.5 mM, [His] = 5 mM in 500 µL aqueous solution (Argon bubbled for 30 min), green light irradiation: 120 J·cm^−2^ (16.6 mW·cm^−2^ for 2 hours). **b** Histidine dioxidation with IDM photocatalysis in H_2_^18^O and mass spectra corresponding to His-2O in H_2_O (retention time: 11.653-11.705 min) and H_2_^18^O (retention time: 11.660-11.772 min) conditions. The reaction mixtures for each condition were further silylated by MSTFA. [IDM] = 0.5 mM, [His] = 5 mM in 500 µL normoxic H_2_O and H_2_^18^O. **c** Suggested mechanisms of water-mediated histidine dioxidation by IDM photocatalysis. **d** His-2O tautomers that can be produced by the mechanism and corresponding ΔG by DFT calculation (vs. His-2O-1, Lactam form). 1,4-cycloaddition product by ^1^O_2_ (His-2O-13) was calculated. DFT calculation was conducted using B3LYP/6-311G+G(d,p). Source data are provided as a Source Data file.

Furthermore, we calculated the Gibbs free energy of 12 possible His-2O tautomers to identify the most thermodynamically stable tautomer using DFT calculations (**Fig. 3d**). Among the 12 structures, His-2O-1, featuring double lactam structures, emerged as the most thermodynamically stable form consistent with lactam-lactim stabilization, which exceeds aromaticity-driven stabilization. To validate lactam-lactim formation, we purified the oxidized histidine mixture and performed FT-IR analysis. Notably, new peaks appeared at 1673, 1204, and 1027 cm□¹, corresponding to C=O stretching (lactam), C–OH stretching (lactim), and C–H out-of-plane bending (loss of aromaticity), respectively (**Supplementary Fig. S13**). Unlike the ^1^O_2_-driven mechanism, IDM photocatalysis can give rise to up to 12 tautomerically stabilized His□2O variants, which function as thermodynamically stable intermediates rather than transient endoperoxides formed in ^1^O_2_□driven mechanisms (**Fig. 3d**). Therefore, •OH-mediated histidine oxidation has the potential to enable intracellular protein mapping without exogenous labeling probes, as the lactam derivatives persist long enough to serve as intermediates for downstream enrichment and analysis.

### Histidine-specific labeling by IDM photocatalysis

We next evaluated the stability of His-2O modifications in intracellular environments. To determine whether His-2O persists on proteins in cell environments, we performed a global proteomics workflow after IDM photocatalysis to measure oxidative modifications (**Fig. 4a**). Based on the LC-MS/MS results, we searched for common •OH□induced oxidative modifications as well as His□2O on a proteome□wide scale. Among the oxidative modifications, peptide spectrum matches (PSMs) for His-2O specifically increased (fold change >12) in the presence of IDM (**Fig. 4b**), indicating that intracellular His-2O is stable under cellular conditions and throughout the proteomics workflow. Importantly, IDM photocatalysis selectively oxidizes histidine over other amino acids.

**Fig. 4.**
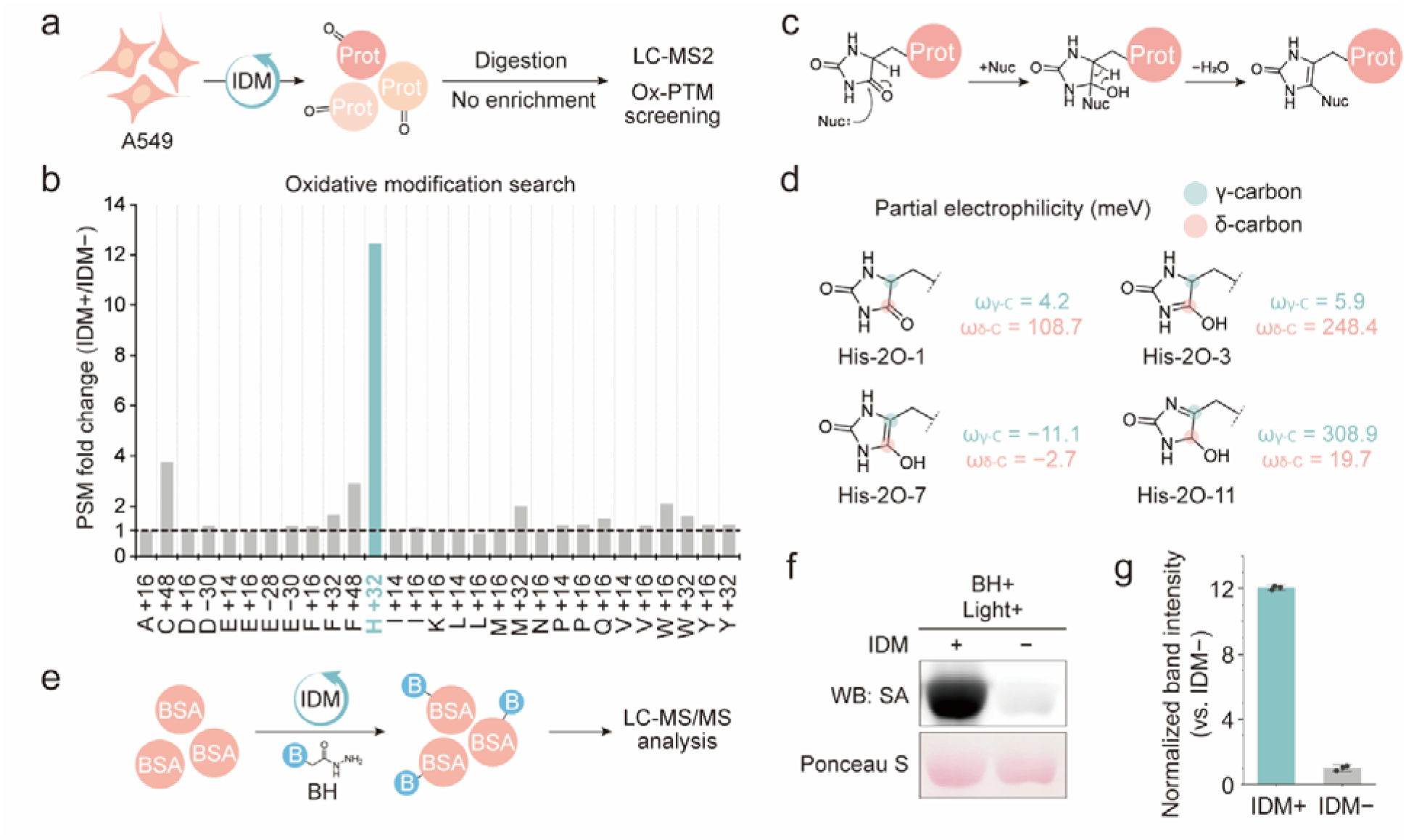
Labeling dioxidized histidine with hydrazide. **a** Brief scheme of global proteomics to investigate histidine specificity of IDM photocatalysis within cells. A549 cells were incubated with 10 µM IDM followed by green light irradiation (16.6 mW·cm^−2^ for 30 min). **b** PSM fold changes (IDM+/IDM−) for various oxidative modifications (hydroxyl radical-induced oxidative modifications) after IDM photocatalysis. **c** Reaction scheme of dioxidized histidine (His-2O-1, lactam form) with nucleophiles. **d** DFT calculating partial electrophilicity of His-2O tautomers. The γ-carbon and the δ-carbon were marked in green and red, respectively. **e** Scheme for dioxidized histidine labeling with BSA. **f** BSA labeling assay in the presence or absence of IDM. Biotin-hydrazide-modified proteins are detected by streptavidin-HRP blotting. Data are shown for three biological replicates. Reaction condition: [IDM]=0.4 mM, [BSA]=1 mg/ml, [Biotin-hydrazide]=1 mM. **g** quantified band intensity of **f**. (n=3). Source data are provided as a Source Data and Source Data Proteomics file.

Next, we investigated suitable chemical handles targeting His-2O to enrich labeled proteins or peptides for identification. These nucleophilic labeling probes can be added during cell lysis, enabling biotinylation for enrichment (**Fig. 4c**). To identify the electrophilic site for labeling, we calculated the partial electrophilicity of His-2O tautomers using DFT calculations (**Fig. 4d** and **Supplementary Fig. S14**). Partial electrophilicity refers to the electron-accepting ability of specific atoms of the structure, enabling the prediction of active sites. In the major tautomer, His-2O-1, the δ-carbon is identified as the highest partial electrophilic site on the imidazole ring, suggesting that this position is the preferred site for nucleophilic attack. Furthermore, all His-2O tautomers that were thermodynamically close to His-2O-1 (within 40 mV in Gibbs free energy) contained at least one highly electrophilic carbon atom (≥ 100 meV), except for His-2O-7. Together, these calculations identify plausible nucleophilic labeling sites on the carbon of His□2O and support targeting His□2O with additional nucleophilic probes.

To verify the hydrazide-driven His-2O enrichment biochemically, we employed bovine serum albumin (BSA). IDM-mediated histidine dioxidation in BSA and subsequent biotin hydrazide (BH) conjugation enable the streptavidin bead enrichment of only biotin-tagged proteins and derived peptides (**Fig. 4e**). Streptavidin-HRP western blot analysis showed strong biotinylation signals exclusively in IDM-treated samples (**Fig. 4f, g**), confirming protein-level labeling. LC-MS/MS further identified His-2O and His-3O modifications (+32/+48 Da) and biotinylation (+270/+272 Da) at several histidine positions of BSA (**Supplementary Fig. S15, S16,** and **S17**). Collectively, these findings demonstrate that IDM photocatalysis enables histidine mapping through dioxidation followed by hydrazide-based tagging and selective enrichment.

### PD-Map and PF-Map identify proteins related to intracellular vesicle trafficking

As the IDM molecular structure contains an anionic cyanoacrylic acid, negatively charged plasma membranes are expected to limit passive diffusion. Additionally, the hydrophobic indenothiophene and the cyanoacrylic acid side chain might trigger interaction with random membrane proteins, facilitating internalization into endosomal vesicles during endocytosis. To confirm IDM internalization, we incubated A549 cells with IDM and LysoTracker (**Supplementary Fig. S18**) to assess colocalization. The images showed a clear dot pattern and partial colocalization with LysoTracker, suggesting that IDM is localized to vesicle trafficking pathways. Additionally, we examined the cytotoxicity of IDM photocatalysis with 3-(4,5-dimethylthiazol-2-yl)-2,5-diphenyltetrazolium bromide (MTT) assay (**Supplementary Fig. S19**). In the presence and absence of green light irradiation, we did not observe significant cytotoxicity across the 2–32 µM concentration range, suggesting that IDM photocatalysis is suitable for histidine labeling without affecting cell viability.

Next, we leveraged IDM for photo-PL to map the proteome involved in intracellular vesicle trafficking. Taking advantage of IDM’s properties, including histidine-specific labeling via dual-radical processes, low cytotoxicity, and biocompatible green-light absorption, we established two complementary workflows: probe-dependent proximal protein mapping (PD-Map) and probe-free proximal protein mapping in live cells (PF-Map). In the PD-Map, A549 cells were treated with IDM and biotin hydrazide (BH), then irradiated with green light (λ=525 nm) to enable intracellular biotinylation. Using confocal microscopy, we imaged biotinylated proteins with a streptavidin-conjugated fluorophore and observed significant spots in the presence of IDM and light irradiation (**Supplementary Fig. S20**). For LC-MS/MS analysis, proteins were digested, and peptides were enriched using streptavidin beads (**Fig. 5a**). Due to the co-incubation with high concentrations of hydrazide probes, this method provides immediate hydrazide labeling of His-2O in live cells.

**Fig. 5.**
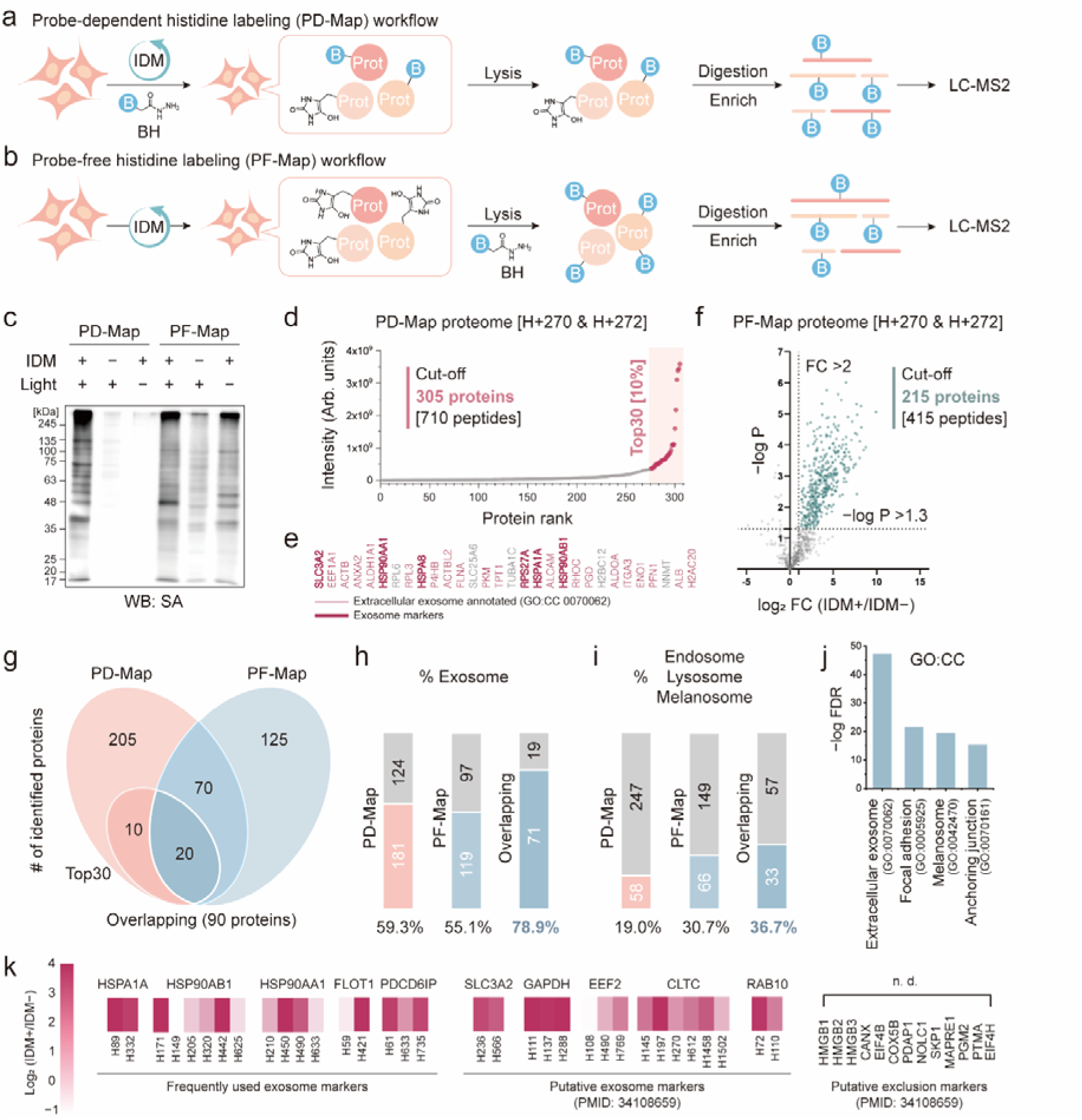
Protein mapping in intracellular vesicle trafficking. **a** Probe-dependent proximal protein labeling (PD-Map) and **b** Probe-free proximal protein labeling (PF-Map) workflow using hydroxyl radical-generating IDM photocatalysis. **c** Comparison of PD-Map and PF-Map biotin-hydrazide labeling upon photoactivation. Cells treated with 10 µM IDM were irradiated with 525 nm light and subjected to biotin-hydrazide labeling either before (PD) or after (PF) lysis, followed by western blot detection using streptavidin-HRP. **d** Intensity-based ranking of the PD-Map–detected proteome. Proteins identified by mass spectrometry were ranked according to their intensity values, and the top 10% (30 of 305 proteins; 710 peptides) were defined as the high-intensity group (shaded area). **e** Protein list of the top 30 ranked proteins of PD-Map. Entries annotated as extracellular exosome (GO:CC 0070062) are colored in red, and established exosome markers are indicated in bold. **f** Volcano plot of PF-Map proteome (H+270 and H+272LJDa). Peptides with a *p*-valueLJ<LJ0.05 and Fold Change (IDM+/IDM−) >LJ2 were defined as PF-Map proteome. **g** Venn diagram representing the number of identified proteins of PD-Map, PF-Map, and overlapping proteins. **h** The ratio of Extracellular exosome (GO:0070062) annotated proteins and **i** the ratio of Endosome, Lysosome, or Melanosome annotated proteins to total PD-, PF-Map, and overlapping proteome. **j** False discovery rate (FDR) of GO:CC terms for the overlapping proteome. **k** Heat map (Fold Change; IDM+/IDM− of PF-Map) of detected histidine for frequently used exosome markers, previously reported putative exosome markers, and putative exclusion markers (PMID:34108659). Source data are provided as a Source Data Proteomics file.

In contrast, the PF-Map method involved incubating cells with IDM alone, followed by light-triggered histidine dioxidation, enabled by the stability of His□2O. Subsequently, the cells were lysed with BH for biotinylation, and the biotinylated proteins were digested and enriched using the same method described above (**Fig. 5b**). We then analyzed protein lysates from each method by western blots using streptavidin-HRP (**Fig. 5c**). The blot showed a high signal-to-noise ratio in PD-Map, likely because the hydrazide is rarely activated by green light in the absence of IDM photocatalysis. Although PF-Map exhibits a higher background signal due to the post-lysis reaction, the method avoids exposing live cells to high BH concentrations, potentially reducing both physiological disturbance and spatial bias arising from the inaccessibility of specific vesicles. Together, these considerations suggest that PD-Map prioritizes high-resolution proximal protein labeling at the cost of probe-induced bias, whereas PF-Map can recover vesicle-associated subproteomes that PD-Map underestimates, indicating that the two methods are mechanistically complementary.

For PD-Map, we selectively extracted peptides exhibiting the expected histidine modification (H + 270 or H + 272) from their MS/MS fragmentation spectra. We defined 710 peptides (corresponding to 305 proteins) that met the cut-off criteria (non-zero median values across triplicates) as the PD-Map proteome (**Fig. 5d**). We identified and listed the top 30 ranked proteins (**Fig. 5e**), and 25 proteins among them were exosome-annotated proteins (GO:CC 0070062). Notably, 6 proteins have been reported as exosome markers. Next, we analyzed 787 histidine-biotinylated peptides (H+270 or H+272) identified by the PF-Map. Among these, we defined the 415 peptides and corresponding 215 proteins that satisfied the cut-off criteria (log_2_FC > 2 and *p* < 0.05, FC: IDM+/IDM−) as the PF-Map proteome (**Fig. 5f**). We then further investigated the PD-Map and PF-Map proteomes and identified 90 overlapping proteins, which include 20 proteins of the top 30 ranked proteins of the PD-Map proteome (**Fig. 5g**). Interestingly, the overlapping proteome showed a high proportion of exosome-annotated proteins (78.9%) compared to the PD-Map and PF-Map proteomes (59.3% and 55.1% respectively) (**Fig. 5h**). Additionally, the proteins of other vesicles (endosome, melanosome, and lysosome) were also enriched in the overlapping proteome (**Fig. 5i**). Furthermore, gene ontology (GO) enrichment analysis of the overlapping proteome highlighted strong associations with extracellular exosomes but also focal adhesions and the melanosome, supporting that PD□Map and PF□Map together can map proteins involved in vesicle trafficking pathways (**Fig. 5j**).

Peptide-level enrichment enabled precise identification of modified histidine residues. As the histidine imidazole ring is chemically versatile, the solvent-accessible histidine often contributes protein functions. To test the accessibility of identified histidines, we calculated the solvent-accessible surface area (SASA) of labeled histidine in the PF-Map proteome. Interestingly, the δ-carbon of oxidized histidine exhibited significantly higher SASA values than the atoms of unmodified histidines (**Supplementary Fig. S21**). This tendency is consistent across other histidine atoms, suggesting that water□mediated dioxidation favors solvent□accessible sites. We next identified labeled histidines of previously reported exosome markers among the overlapping proteomes (**Fig. 5k**)^34^. The overlapping proteome successfully detected many putative exosome markers previously suggested in studies, whereas 13 proposed putative exclusion markers were not found, further validating the specificity of PD/PF-Map. Furthermore, we identified specific histidines proximal to the IDM cargo during vesicle trafficking.

### PF-Map reveals vesicle trafficking-related proteome undetected in PD-Map

In PD□Map, live-cell labeling requires BH to access His-2O sites, so vesicular proteins that are poorly accessible to BH are likely underrepresented. Such underrepresentation implies that even functionally important vesicular proteins can remain largely invisible to conventional probe□dependent PL workflows, thereby narrowing the apparent scope of vesicle trafficking-related proteomes. Of the 215 proteins identified by PF-Map, 125 were not significantly detected in PD-Map. We hypothesize that these PF-Map-specific proteins may reflect differences in probe accessibility within the cellular environments. To further characterize this PF□Map-specific subproteome, we constructed the Circos plot to visualize their interaction network and performed GO term enrichment analysis (**Fig. 6a**). The Circos plot revealed a densely interconnected protein–protein interaction network, indicating that these 125 proteins form a coherent functional module enriched in transport (GO:0006810)- and localization (GO:0051641)-related processes consistent with vesicle trafficking. For example, PF-Map identified Rab5 and Rab11^35, 36^, which play central roles in vesicle recycling and trafficking. Furthermore, SEC31A is an essential component of the COPII coat complex that mediates vesicle budding from the endoplasmic reticulum (ER) to the Golgi apparatus^37^, suggesting a potential crosstalk between COPII-dependent secretory trafficking and endosomal/exosomal pathways. Together, these analyses suggest that PF-Map captures an additional vesicle trafficking–related subproteome that is underrepresented in the PD-Map, likely reflecting spatial bias in intracellular BH distribution.

**Fig. 6.**
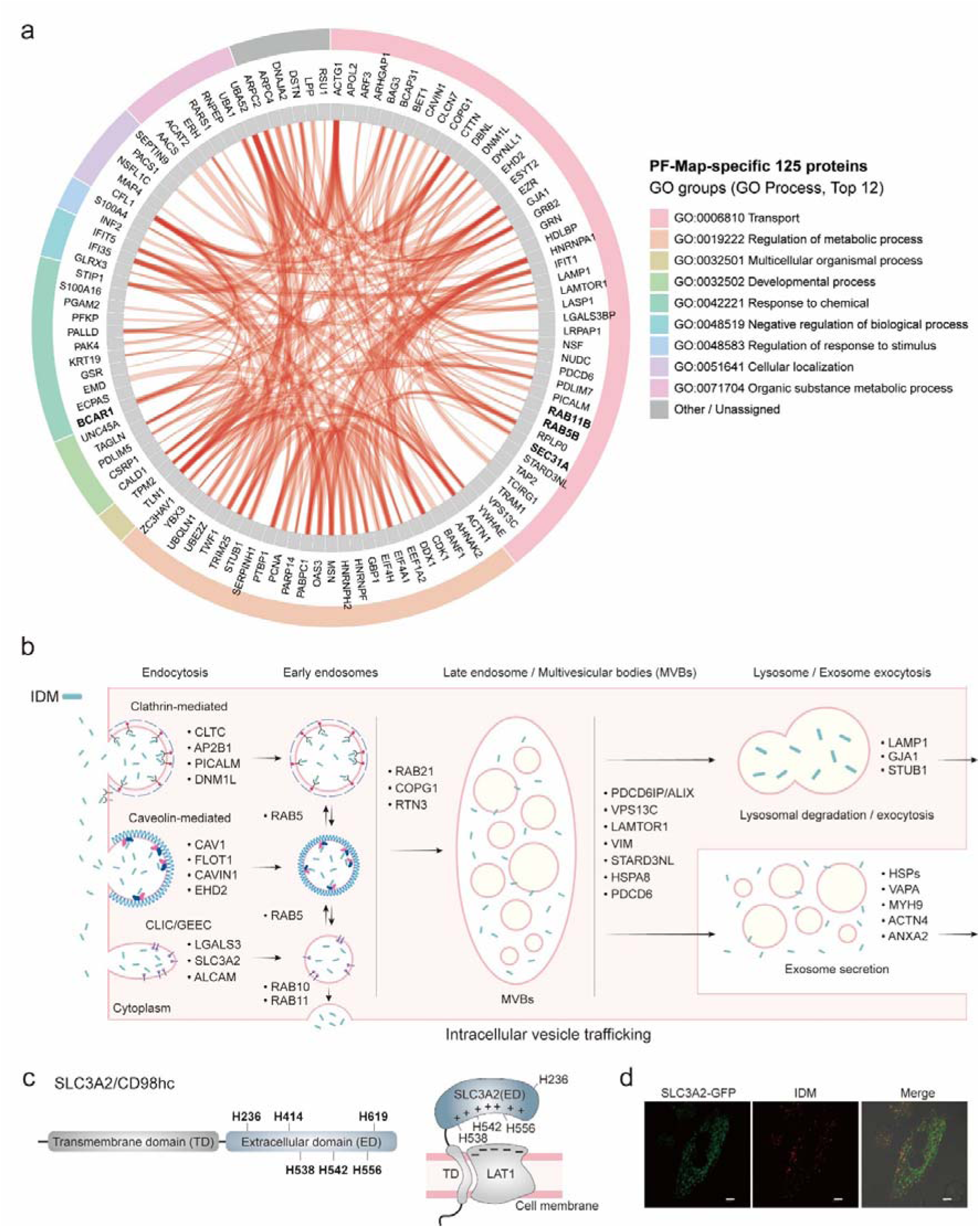
PF-Map analysis to uncover a vesicle trafficking–related proteome. **a** The Circos plot to reveal protein–protein interaction network and GO term classification of 125 proteins uniquely identified in the PF-Map. **b** Suggested proteome related to intracellular exosome trafficking of IDM cargo. The PF-Map proteome was categorized into Endocytosis, Early endosome, Late endosome/MVBs, and Lysosome/Exosome. **c** Identified histidines of SLC3A2. H538, H542, and H556 are located at positively charged sites that interact with LAT1. **d** Co-localization image with SLC3A2-GFP (green) and IDM (red) using A549 cells. scale bar = 10 µm.

Based on PF-Map, we outlined the intracellular vesicle trafficking pathway in A549 cells by grouping the identified proteins into four stages: endocytosis, early endosomes, late endosomes/multivesicular bodies (MVBs), and exosomes (**Fig. 6b**). PF-Map revealed multiple endocytic entry routes, including clathrin-dependent, caveolar, and clathrin-independent CLIC/GEEC pathways. Proteins that regulate vesicle trafficking were robustly detected in subsequent stages. In the late endosome/MVB stage, we observed proteins such as PDCD6IP/ALIX, VPS13C, HSPA8, and PDCD6, which coordinate cargo sorting and vesicle maturation. Notably, both PD-Map and PF-Map robustly enriched SLC3A2 (also known as CD98 heavy chain; CD98hc), a transmembrane protein that plays a crucial role in the transport of bulky aromatic amino acids by conjugating with LAT1^38, 39^. All the identified histidines are in the extracellular domain, and H538, H542, and H556, in particular, are located on a positively charged interaction site with LAT1 (**Fig. 6c**). Accordingly, we verified the co-localization of SLC3A2 and IDM using confocal microscopy. The images showed that intracellular SLC3A2-GFP fluorescence partially overlapped with the IDM signal, suggesting that SLC3A2 is consistent with a role in endocytosis-mediated internalization of small-molecular cargo (**Fig. 6d**). By capturing proteins across early endosomes, late endosomes/multivesicular bodies, and exosomes, PF□Map maps vesicle trafficking over multiple stages rather than focusing on a single vesicle class. This broad coverage could reduce the dependence of spatial proteomics on probe□accessible vesicle subsets and thus provide a less biased view of vesicular proteomes by eliminating the need for exogenous capture probes during live-cell proximity labeling.

## Discussion

In this study, we establish a redox-guided strategy to harness endogenous water and oxygen to generate radical precursors for proximity labeling and apply this principle in two complementary photo-PL workflows, PD-Map and PF-Map. We developed an indenothiophene-based organic photocatalyst, IDM, that generates •OH, O_2_^•−^, and H□O□ but not detectable ^1^O_2_ under green-light irradiation. GC–MS analyses combined with DFT calculations support a mechanism in which histidine dioxidation yields thermodynamically stable lactam structures (His-2O) that persist through lysis and proteomic workflows. In contrast to conventional photo-PL strategies that couple oxidative modification with immediate probe trapping in live cells, IDM oxidizes histidines via water oxidation, and His-2O is subsequently tagged post-lysis with hydrazide probes, thereby decoupling oxidative labeling from live-cell probe accessibility. We applied probe-free proximal protein mapping in live cells (PF-Map) to protein mapping along intracellular vesicle trafficking, a pathway comprising highly heterogeneous vesicles. Leveraging the same photocatalyst, IDM, in two orthogonal workflows, comparative proteomics showed that PD-Map and PF-Map share a core vesicle trafficking proteome enriched in established exosome markers, while PF-Map additionally captures a coherent subproteome of 125 proteins (e.g., Rab5, Rab11, SEC31A) that are underrepresented in PD-Map. PD-Map, in turn, provides high signal-to-noise proximal labeling and a stringent core vesicle marker reference for interpreting PF-Map–specific expansions. Together, these complementary PF-Map and PD-Map workflows establish a versatile photo-PL toolbox for protein mapping in dynamic subcellular compartments.

Given histidine’s essential roles in enzymatic catalysis and protein-protein interactions^40–42^, it has been regarded as an attractive target for photo-PL^14, 43^. Functional histidines are dynamically protonated and deprotonated to sense microenvironmental pH and catalyze biomolecular reactions, thereby contributing to diverse bioprocesses across different organelles^44, 45^. To capture a snapshot of histidines engaged in a specific bioprocess, the PF-Map strategy can be used as a chemical tool to minimize microenvironmental perturbations and spatial biases caused by exogenous capture probes during live-cell proximity labeling. PF-Map may also be applicable to in vivo biological systems, where the accessibility of exogenous chemical probes is limited^46, 47^, because the absorption range of IDM (500–630 nm) overlaps with the tissue-penetrant optical window. Although methods for precisely localizing IDM to specific cellular compartments remain to be developed, we anticipate that PF-Map will provide a promising approach for mapping proteins in complex bioprocesses. From a chemical perspective, this study demonstrates that photocatalysts with precisely tuned redox potentials can harness water oxidation to generate radical species for endogenous protein labeling without requiring exogenous capture probes during live-cell proximity labeling. This design principle, stable oxidative modification followed by post-lysis chemical addressing, may inform future strategies for spatial proteomics and other proximity-dependent chemical biology applications.

## Methods

### Synthesis & Characterization

The methods for organic synthesis of photocatalysts, IDM, are described in Supplementary Information. All chemicals used in organic synthesis were purchased from Sigma Aldrich, Alfa Aesar, Tokyo Chemical Industry, Combi-Blocks, and SAMCHUN. In the synthetic process, the newly synthesized chemical compounds were analyzed by ^1^H, ^13^C NMR (Agilent 400MR-DD2 NMR spectroscopy), FT-IR (Varian Cary 620/670 FT-IR spectrometer), and HRMS (Bruker maXis^TM^ HD Ultra-high-resolution Q-TOF LC-MS/MS system, The Cooperative Laboratory Center of Pukyong National University, Republic of Korea).

### Photophysical properties (Abs, PL, and CV)

The ground-state redox potentials of TP-1 and TP-2 were determined by cyclic voltammetry (CV) using a three-electrode setup. A glassy carbon electrode drop-cast with the photocatalyst dye (TP-1 or TP-2) served as the working electrode, an Ag/AgNO□ electrode (0.1 M TBAPF_6_ in acetonitrile) as the reference electrode, and a platinum wire as the counter electrode. The electrolyte solution consisted of 0.1 M TBAPF_6_ in acetonitrile. CV measurements were conducted with a step potential of 1 mV and a scan rate of 10 mV·s□¹. The oxidation and reduction potentials were referenced to the normal hydrogen electrode (NHE) after calibration. For TP-1, the E^(0/−)^ was −1.20 V vs. NHE, and the E^(+/0)^ was 1.08 V vs. NHE. For TP-2, E^(0/−)^ = −1.47 V and E^(+/0)^ = 0.96 V vs. NHE. The redox properties of TP-3 were measured under aqueous conditions to reflect physiological relevance. A glassy carbon electrode was used as the working electrode, Ag/AgCl (saturated KCl solution) as the reference electrode, and a platinum wire as the counter electrode. The electrolyte was 1× PBS solution (pH = 7.4), and 20 μL of 5 mM TP-3 stock solution was dissolved directly in the electrolyte. CV was performed with Estep = 1 mV and a scan rate of 10 mV·s□¹. All potentials were converted to the NHE scale for comparison. For TP-3, E^(0/−)^ = −0.52 V and E^(+/0)^ = 1.29 V vs. NHE. To calculate excited-state redox potentials, previously reported E^0/0^ values were used^21^. Note that only TP-3 exhibits a more positive excited-state reduction potential (E*^(0/−)^ = 1.57) than the two-electron water oxidation potential (Eox = 1.34 V).

Aqueous solutions of IDM (20 µM, H_2_O:DMF = 995:5 v/v) were prepared for measuring the absorbance and photoluminescence of IDM. The UV-visible spectrometer (SHIMADZU UV-2600 240V EN, Japan) and fluorescence spectrometer (ISS PC1 photon counting spectrofluorometer, USA) were utilized. The absorption and fluorescence spectra were normalized to the maximum value in the visible range. Then, we determined the wavelength at the cross-point of the absorbance and fluorescence curves to set E^0/0^ at 585 nm (2.12 eV). The photocatalytic redox potentials were calculated using method^48^.

The excited-state redox potentials of IDM were calculated from its oxidation and reduction potentials measured by three-electrode cyclic voltammetry (CV). It was performed in 1 XPBS using a glassy carbon electrode drop-cast with IDM as the working electrode, Ag/AgCl as the reference electrode, and a platinum wire as the counter electrode. The CV was measured with a step potential of 1mV and a scan rate of 20 mV·s^−1^.

In the reductive quenching cycle, excited IDM has the proper potential to oxidize water. Thus, photoinduced electron transfer can occur between excited IDM and water molecules, thereby reducing IDM fluorescence. To this end, the 20 μM IDM solution (in acetonitrile) was prepared to measure fluorescence changes as a function of H_2_O content. The fluorescence of IDM was measured as the H_2_O content increased from 0% to 10% at 2% intervals.

### Reactive oxygen species (ROS) assay

The hydrogen peroxide generation by IDM photoactivation was measured by the horseradish peroxidase (Sigma Aldrich, USA) and N, N-diethyl-*p*-phenylenediamine (DPD) (Sigma Aldrich, USA). First, stock solutions of peroxidase (1 mg·mL-1 in DI water) and DPD (50 mM in 1 M H2SO4) were prepared. Then, the three 50 μM IDM solutions (in normoxic PBS and Ar-bubbled PBS) were irradiated by the green LED (λ_max_ = 525 nm, 66.7 mW·cm^-2^) (HepatoChem Inc., USA). During the light exposure, assay samples were obtained at 30-minute intervals up to 150 minutes. 200 L of each sample solution was dissolved in 224 L of DI water, and 80 μL of sodium phosphate buffer (pH 6) was added to each sample solution. Then, 10 μL of DPD and peroxidase stock solution were added. Right after that, the absorbance of each sample solution at 551 nm was measured. The results were obtained from three distinct experimental samples.

Hydroxyl radicals were detected using an HPF assay. HPF (Invitrogen) is an indicator of hydroxyl radicals and peroxynitrite. The hydroxyl radical eliminates the hydroxyphenyl group of HPF, and HPF is subsequently converted to the fluorescent form (fluorescein). Thus, we prepared aqueous IDM (5 μM) and HPF solutions (5 μM) and measured fluorescence at 515 nm. In this assay, the final concentration of IDM was 5 μM. Furthermore, green LED (λ_max_ = 525 nm) (HepatoChem Inc., USA) was used to photoactivate IDM (16.6 mW·cm^-2^ for 2 min = 2 J·cm^-2^). A microplate reader (SpectraMax M5e, USA) was used to measure fluorescein fluorescence. Results were obtained using three distinct experimental samples.

Additionally, •OH generation by photocatalysis of TP-1, TP-2, and TP-3 were measured by HPF assay. Stock solutions of TP-1, TP-2, and TP-3 (5 mM) and HPF (10 mM) were prepared in DMF. These were diluted 1,000-fold into 2 mL of distilled water. The reaction mixtures were then irradiated with 525 nm LED light for 0, 2, 4, 6, or 8 min (λ_max_ = 525 nm, 16.6 mW·cm^−2^). After irradiation, 100 μL of each sample was transferred to a 96-well black plate, and the fluorescence intensity of fluorescein was recorded on a microplate reader (Ex 485 nm, Em 515 nm).

Singlet oxygen generation was examined by the 9,10-anthracenediyl-bis(methylene)dimalonic acid (ABDA) assay. 100 mM ABDA stock solution (1000X) was supplemented with 5 µM Eosin Y (EY) and IDM solution. Solutions were irradiated with a green LED (λ_max_= 525 nm, 16.6 mW·cm^−2^) for 0, 2, 4, and 6 minutes. Then, the ABDA absorbance at 400 nm was detected by a microplate reader. The ABDA absorbance decay of each sample was recorded as Abs_ABDA, 400nm_ − Abs_sample, 400nm_ to correct the baseline.

### Electron paramagnetic resonance spectroscopy to measure ROS generation

EPR spectroscopy was performed to measure singlet oxygen generation under room temperature using 4-hydroxy-2,2,6,6-tetramethylpiperidine (4-OH-TEMP) as a trap for singlet oxygen to yield 4-hydroxy-2,2,6,6-tetramethylpiperidine 1-oxyl (4-OH-TEMPO) free radical. A 100 mM stock solution of 4-OH-TEMP in water (50 μL) and a 10 mM stock solution of IDM and Eosin Y in DMF (1 μL) were mixed in a 1.5ml microtube. Then, the sample solutions were irradiated with a green LED (λ_max_= 525 nm, 1.5 J·cm^−2^) for 6 minutes. Subsequently, the samples were transferred to an EPR capillary tube for EPR spectroscopy.

EPR spectroscopy with a 5,5-dimethyl-1-pyrroline *N-*oxide (DMPO) as a trap to yield DMPO-OH was employed to identify the hydroxyl and superoxide radicals. DMPO solutions with IDM were prepared ([IDM] = 1 mM and [DMPO] = 100 mM) in microtubes, irradiated with weak green light (λ_max_ = 525 nm, 0.1 J·cm^−2^), and the EPR spectra were measured. To verify the effect of histidine on the generated hydroxyl radicals, we prepared a reaction solution containing DMPO, IDM, and histidine ([IDM] = 1 mM, [His] = 7 mM, [DMPO] = 100 mM), and subsequently measured the EPR spectra.

To measure superoxide radicals, we used EPR spectroscopy to detect DMPO-OOH signals in anhydrous, oxygen-purged DMF. A 1 M stock solution of DMPO in DMF (5 μL) and a 10mM stock solution of IDM in DMF (1 μL) were mixed with an oxygen-purged DMF solution (24 μL) and transferred into a 1.5ml microtube. The diluted mixture was then irradiated with a 525 nm green LED for 5 minutes (1 J·cm^−2^) and transferred to an EPR capillary tube for EPR spectroscopy.

EPR measurements were performed at Ulsan National Institute of Science and Technology (UNIST) UCRF. X-band (9.86□GHz) EPR spectra were derived using a Bruker EMX Plus 6/1 spectrometer equipped with a dual-mode cavity (ER 4116DM). The spectra were obtained using the following experimental parameters: microwave frequency, 9.86□GHz; microwave power, 3.99□mW; modulation amplitude, 1G; time constant, 10.24□ms; 5 scans.

### Gas chromatography-Mass spectroscopy (GC-MS) for identification of histidine oxidation

Histidine oxidation was performed prior to identifying products through gas chromatography-mass spectrometry (GC-MS) (Agilent, GC/MS 8890A/5977C, USA). A reaction mixture (0.5 mM IDM and 5 mM amino acids in H_2_O:DMF (95:5, v/v) was prepared and irradiated with green light (λ_max_ = 525 nm, 16.6 mW/cm^2^) for 2 hours. Following irradiation, the solvent was evaporated entirely, and the residue was dried under vacuum. The resulting materials were dissolved in 800 μL of acetonitrile and 200 μL of MSTFA + 1 % TMCS. The mixture was incubated at 70 □ for 20 min for silylation. The silylated sample was analyzed by GC-MS using an HP-5ms column (30 m X 0.25 mm X 0.25 μm). Helium was used as the carrier gas in split mode. The inlet temperature was set to 240 °C. The oven was initially held at 100 ^°C, then^ ^ramped^ ^to^ ^300^ ^°C^ ^at^ ^a^ ^heating^ ^rate^ ^of^ ^15^ ^°C^ min^−1^.

Isotope-labeling experiments were conducted to validate histidine oxidation via hydroxyl radicals generated from water. The photocatalytic oxidation of histidine was performed using a reaction mixture (0.5 mM IDM and 5 mM amino acids in H218O:DMF = 95:5, v/v) under green-light irradiation (λmax = 525 nm, 16.6 mW/cm2) for 2 hours. Both sample preparation and the GC-MS measurement method followed the general histidine oxidation product analysis.

### FT-IR analysis of oxidized histidines

For FT-IR analysis of oxidized histidine structures, histidine was first oxidized by IDM photocatalysis. The reaction mixture (0.05 mM IDM and 5 mM histidine in H_2_O:DMF = 1:200, v/v, total 12 mL) was irradiated with a 525 nm LED light for 6 hours. The resulting crude mixture was purified by high-performance liquid chromatography (HPLC) (Waters, Modular system, USA). The mobile phase was 0.1% trifluoroacetic acid in HPLC-grade methanol with 18.2 ΩM cm^−1^ water, and the stationary phase was a C18 column. The flow rate was 2 mL min^−1^, the column temperature was about 25 □, and a UV detector set at 230 nm was used. After each purified product was collected, the solvent was removed by evaporation. Finally, FTIR was measured using diamond crystal attenuated total reflection (ATR) geometry on a PerkinElmer Spectrum Two. They were performed from 4000 cm^−1^ to 400 cm^−1^ with 16 scans.

### Density functional theory (DFT)

Gibbs free energy and electrophilicity index were calculated using density functional theory (DFT) calculations with the Gaussian 09W program. All calculations were performed at the B3LYP/6-311G(d,p) basis set. Geometry optimization and frequency calculations were carried out for the ground state of histidine dioxidized tautomers, resulting in Gibbs free energy.

Global electrophilicity index (ω) was calculated using the formula^49, 50^:

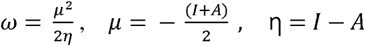

where I and A denote the ionization potential and electron affinity, respectively. To obtain these values, the total electronic energies of the neutral, one-electron-oxidized, and one-electron-reduced states were calculated. Specifically, I was calculated as the energy difference between the oxidized and neutral states, and A as the energy difference between the neutral and the reduced states.

Local electrophilicity index was computed by multiplying global electrophilicity by the Fukui function (f_k_) of each interested atom.

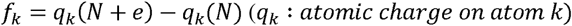

This value can be obtained from Natural Population Analysis (NPA) within the Natural bond orbital (NBO) analysis. These calculations were also performed at B3LYP/6-311G(d,p) level of theory.

### Proteomics for identification of global oxidative modifications

A549 cells were cultured in 6-well plates for 24 h and treated with 10 µM IDM in DMEM for an additional 24 h. Triplicate wells were treated with IDM, and three wells were left untreated as controls. Cells were washed three times with DPBS, incubated with 2 mL DPBS, and irradiated with 525 nm LED light at room temperature for 30 min. After irradiation, cells were harvested, lysed in 2% SDS in TBS supplemented with protease inhibitor cocktail, and proteins were precipitated with acetone. The resulting pellets were reconstituted in 8 M urea in 50 mM ammonium bicarbonate (ABC), and protein concentration was determined by BCA assay. A total of 70 µg protein per sample was loaded on SDS-PAGE and electrophoresed at a constant current of 15 mA until the dye front migrated approximately 1 cm.

The gel was stained and each lane was excised into five gel pieces (1 × 1 mm). Gel pieces were sequentially destained with 200 µL of methanol/50 mM ammonium bicarbonate (1:1, v/v) at 56 °C for 3 min with gentle agitation, and the process was repeated twice until the gel became colorless. The destained pieces were then washed once with 200 µL of 40% acetonitrile in water at 56 °C for 3 min, followed by dehydration with 200 µL of 100% acetonitrile under the same conditions. After complete dehydration, the solvent was removed, and the gel pieces were air-dried before reduction and alkylation.

For reduction, 25 mM DTT in 50 mM ammonium bicarbonate was added and incubated at 56 °C for 20 min, followed by alkylation with 100 mM iodoacetamide in 50 mM ammonium bicarbonate for 30 min in the dark at room temperature. And then, the gel pieces were dried and digested overnight at 37 °C with trypsin (enzyme-to-protein ratio of 1:50, w/w) in 50 mM ABC containing 10% acetonitrile. Peptides were extracted with 50% acetonitrile/0.1% formic acid followed by 99.9% acetonitrile/0.1% formic acid, and the combined extracts were dried in a SpeedVac.

Peptide cleanup was performed using C18 StageTips (10 µL bed volume). Tips were conditioned with 50% acetonitrile and 0.1% TFA, equilibrated with 0.1% TFA, and loaded with samples dissolved in 0.1% TFA. Bound peptides were washed with 0.1% TFA containing 5% acetonitrile and eluted with 0.1% formic acid in 80% acetonitrile. Eluates were dried by SpeedVac, reconstituted in 0.1% formic acid, and stored at −20 °C until LC–MS/MS analysis.

The raw MS data were converted to mzML format using MSConvert (v3.0.25035). To globally characterize diverse oxidative modifications on amino acids, a modification-tolerant search was performed using MODplus (v1.02) against the UniProt protein database^51^. The search parameters were defined as follows: a precursor mass tolerance of ±10 ppm with 13C isotope errors of –1, 0, 1, or 2; a fragment mass tolerance of ±0.4 Da; and trypsin specificity with a minimum of one enzymatic terminus. Missed cleavages were permitted without restriction. Carbamidomethylation of cysteine was set as a fixed modification. Variable modifications included common MS modifications predefined in MODplus, histidine dioxidation, and fast photochemical oxidation of proteins (FPOP)-related modifications (Supplementary Table S1)^52, 53^. The search allowed an unlimited number of modifications per peptide within a mass shift range of –150 to +350 Da. A decoy-based strategy was employed for false discovery rate (FDR) estimation. All peptide-spectrum matches (PSMs) were subsequently rescored via Percolator^54^, and final identifications were filtered at 1% FDR.

### Western blot to verify BSA labeling

For in vitro BSA labeling, total 5 mg of BSA at a final concentration of 1mg/ml, was mixed with 0.4 mM IDM and 1 mM biotin-hydrazide in 50 mM ammonium acetate buffer to final volume 250 uL.

The protein samples were then exposed to a 525 nm LED (HepatoChem Inc.) for 30 minutes. After this, the proteins were precipitated using 5-6 times the volume of -20°C acetone and incubated at -20°C for a minimum of 2 hours. Subsequently, the protein pellet was reconstituted in 400 μL of an 8 M Urea (Sigma, U5378) in 50 mM ammonium bicarbonate (ABC; Sigma, A6141) buffer. The samples were then prepared for Western blot analysis or mass sampling as needed.

### In vitro BSA labeling

5mg of BH labeled BSA, precipitated and solubilized in 400 μL of 8 M urea in 50mM ABC buffer, was reduced with DTT, alkylated with IAA, and digested with trypsin. Peptides were enriched using streptavidin magnetic beads (100 μL per sample), eluted with 80% acetonitrile containing 0.1% formic acid and 0.2% TFA, dried by SpeedVac, and analyzed by LC–MS/MS.

To explore the types of modifications present in the sample, a blind search was performed using MODplus, enabling the detection of unknown modifications as mass shifts (delta masses). In blind search mode, the precursor ion mass tolerance was set to be flexible to allow compensation for isotope errors with automatic correction, while a fragment ion mass tolerance of 0.02 Da was applied. Peptides were assumed to be potentially modified within a mass shift range of −150 to +350 Da. The search was conducted without enzyme specificity, and an unlimited number of modifications per peptide was permitted. Finally, all MS/MS spectra assigned with mass shifts were subjected to manual inspection to verify the accuracy and localization of the mass shifts.

### Co-localization confocal imaging

Confocal laser scanning microscopy (CLSM) was utilized to identify the subcellular localization of IDM. One or two days before the experiment, A549 cells were seeded on a coverglass-bottom confocal dish. One day after cell seeding, the cells were incubated with IDM (10 μM in media:DMF = 200:1, v/v) for 24 hours and LysoTracker Green DPD (LysoG; 50 nM) for 30 minutes. Then, the co-localization images were obtained in the CO_2_ incubator at 37 °C under a humidified atmosphere and 5% CO_2_ by Carl Zeiss LSM980 with a 63X lens. (LysoG: λ_excitation_ = 488 nm, emission gain range: 500-530 nm; IDM: λ_excitation_ = 561 nm, emission gain range: 564-693 nm).

### PD-Map LC-MS/MS preparation

A549 cells were split into three 150 mm pi culture dishes and cultured to obtain triplicate samples. The following day, the cells were treated with 10 µM IDM in DMEM (Gibco) for overnight. On the subsequent day, the cells were washed twice with DPBS and pre-incubated with 250 μM biotin-hydrazide in DPBS for 30 minutes, followed by irradiation with a 525 nm LED (HepatoChem Inc.) for 30 minutes. After washing the cells three times with DPBS, cells were collected to 1.5 mL microcentrifuge tubes and stored at -80°C until lysis.

For protein extraction, a 2% SDS (Sigma, 436143) solution containing 1x protease inhibitor in 1x TBS (25 mM Tris, 0.15 M NaCl, pH 7.2) was used. The extraction process involved sonication with a Bioruptor (diagenode) for 30 minutes, repeated 2-3 times at 4°C. To remove residual chemicals or lipids, the sample volume was mixed with 5-6 times -20°C acetone and incubated at -20°C for at least 2 hours. The mixture was then centrifuged at 13,000 × g for 10 minutes at 4°C, separating the precipitated proteins. The protein pellet was dissolved in 200 μL of 8 M Urea in 50 mM ammonium bicarbonate (ABC) buffer.

Protein concentration was determined using the BCA (bicinchoninic acid) assay. Next, 0.5 mg of protein samples were denatured by incubating at 37°C and 650 rpm for 1 hour in a thermomixer (Eppendorf). The samples were reduced at 37°C and 650 rpm for 1 hour using 10 mM dithiothreitol (Sigma, 43819), followed by alkylation with 40 mM iodoacetamide (Sigma, I1149) at 37°C and 650 rpm for 1 hour. The samples were then diluted 8-fold with 50 mM ABC buffer, and 1 M CaCl_2_ was added to reach a final concentration of 1 mM. TPCK-Trypsin (Thermo Fisher 20233, USA) digestion (50:1 w/w ratio) was carried out in the thermomixer at 650 rpm and 37°C for 20 hours, and insoluble material was removed by centrifugation at 13,000 × g for 15 minutes.

Streptavidin-conjugated beads (100 μL; Pierce) were washed three times with 2 M Urea in 1x TBS and then mixed with the samples for enrichment at room temperature for 1 hour. The flow-through fraction was removed, and the beads were washed twice with 2 M Urea in 50 mM ABC, once with 50 mM ABC, and finally once with pure water. Subsequently, elution was carried out at 60°C and 650 rpm using 150 μL of elution buffer (80% acetonitrile, 0.2% trifluoroacetic acid, 0.1% formic acid). The eluted upper fraction was collected in a new tube, and this elution process was repeated at least three times and combined in one tube. The eluted solution was dried using a Speed-Vac (Eppendorf) and stored at -20°C until it was injected into the mass spectrometry instrument.

### PF-Map LC-MS/MS preparation

A549 cells were seeded on 150-mm dishes and cultured for 24 h prior to treatment. Cells were incubated with 10 µM IDM in DMEM for an additional 24 h, and triplicate samples were treated with IDM while three untreated dishes served as controls. After incubation, cells were washed three times with DPBS, replenished with 10 mL fresh DPBS, and irradiated with 525 nm LED light at room temperature for 30 min.

Cells were then harvested, and pellets were lysed in 500 µL of 2% SDS in TBS containing protease inhibitor cocktail and 0.5 mM biotin-hydrazide, followed by ultrasonication. Lysates were subjected to dialysis using Slide-A-Lyzer™ MINI Dialysis Devices (10 K MWCO, Thermo Fisher Scientific, Cat# 88404) three times, concentrated using Amicon filters, and subjected to acetone precipitation. After that , approximately 1 mg of total protein remained per sample. Proteins were reduced with DTT, alkylated with IAA, and digested with trypsin prior to enrichment using 100 µL streptavidin magnetic beads per sample. Bound and peptides were eluted with 80% acetonitrile containing 0.1% formic acid and 0.2% TFA, then dried by SpeedVac prior to LC–MS/MS analysis.

### LC-MS/MS measurement

The resulting digested peptides were analyzed by LC-MS/MS. All mass analyses were performed on a Q Exactive Plus orbitrap mass spectrometer (Thermo Fisher Scientific, MA, USA) equipped with a nanoelectrospray ion source. Peptides were separated on a 75 μm inner diameter microcapillary column packed with 50 cm of RSLC C18 resin (2μm, 100 Å, PepMAP, Thermo Fisher Scientific). For each analysis, separation was achieved using a 190min gradient of 3 to 97% acetonitrile in 0.1% formic acid at a flow rate of 0.3 nl/min. MS2 analysis consisted of high-energy collision-induced dissociation (HCD) with the following settings: resolution, 17,500; AGC target, 1 × 105; isolation widov 2 m/z; NCE, stepped: 27; and maximum injection time, 100 ms. For MS/MS analysis, the precursor ion scan MS spectra (m/z 400–2000) were acquired in the Orbitrap at a resolution of 70 000 at m/z 400 with an internal lock mass. The 15 most intensive ions were isolated and fragmented by high-energy collision-induced dissociation.

### PD-Map/PF-Map analysis

The raw MS data were converted to mzML format using MSConvert, and the converted files were searched against the UniProt protein database using MSFragger (v3.8) through FragPipe (v20.0). The search parameters were as follows: enzyme specificity was set to strict trypsin with up to two missed cleavages and a requirement of two enzymatic termini; precursor mass tolerance was 10 ppm and fragment mass tolerance was 0.4 Da; 13C isotope errors of 0, 1, or 2 were allowed. Fixed modification was carbamidomethylation of cysteine residues. Variable modifications included methionine oxidation, protein N-terminal acetylation, histidine oxidations (+31.99 Da and +47.985 Da), and histidine biotinylations (+254.084 Da, +270.079 Da, +272.094 Da, and +288.089 Da). A maximum of three variable modifications per peptide was allowed, regardless of modification type. A decoy-based search strategy was employed, and all PSMs were filtered at 1% FDR using Percolator. Quantification and match-between-runs (MBR) was performed using IonQuant.

### SASA calculation for histidine residues

UniProt amino acid sequences of all MS-detected proteins were analyzed in R to identify possible tryptic peptides containing histidine residues. The selection criteria allowed no missed cleavages and peptide lengths between 6 and 40 amino acids. Solvent-accessible surface areas (SASA) of histidine residues were calculated from AlphaFold protein models using Biopython’s rolling-ball algorithm, with a probe radius of 1.4 Å and 960 surface points per atom.

### Plasmid construction

The coding sequences of EGFP–ALIX and SLC3A2–EGFP were cloned into pLenti-CMV vectors. For EGFP–ALIX, the EGFP and ALIX fragments (from pLVX-ALIX-GFP, Addgene #138582) were fused by overlap PCR using AccuPower® PCR PreMix (Bioneer) with the following cycling conditions: 95 °C for 5 min; 35 cycles of 95 °C 30 s, 63 °C 30 s, 68 °C 1 min per kb; final extension 68 °C 10 min. The fused fragment was digested with BamHI and SalI and inserted into pLenti-CMV-cGAS-HA (Addgene #130910) digested with the same enzymes. For SLC3A2–EGFP, the SLC3A2 coding sequence was amplified from pDONR221-SLC3A2_STOP (Addgene #161379) and subcloned into the pLenti-CMV vector so that EGFP was fused to the C-terminus of SLC3A2. The insert and vector were digested with XbaI and SalI and ligated using T4 DNA ligase. All constructs were propagated in E. coli DH5α and selected on LB plates containing ampicillin (100 µg mL□¹). All constructs were verified by Sanger sequencing to confirm the correct DNA sequence after cloning.

### Lentivirus infection

Lentiviral particles were produced in HEK293FT cells (6-well format, ∼70–80% confluency) using PEI transfection. Per well, pLenti construct (1,000 ng), psPAX2 (750 ng), and pMD2.G (250 ng) (total 2.0 µg DNA) were mixed with PEI (linear, 25 kDa; DNA:PEI 1:3 w/w) in serum-free DMEM, incubated for 15 min at room temperature, and added dropwise to the cells. At 24 h post-transfection, culture supernatants were collected, clarified, and passed through a 0.45 µm syringe filter to obtain the virus solution. A549 cells in 6-well plates were transduced by adding 1 mL of the virus solution per well. After 2 h incubation at 37 °C, 1 mL fresh DMEM was added to each well (final 2 mL/well). The following day, cells were passaged and selected with puromycin (1 µg mL□¹) in DMEM for 5–7 days until non-transduced control cells were fully eliminated.

### Imaging SLC3A2-GFP

A549 cells infected with SLC3A2–GFP were seeded onto glass coverslips placed in 12-well plates and cultured overnight. On the following day, the culture medium was replaced with DMEM containing IDM (10 μM), and cells were incubated overnight. Cells were then fixed with 4% paraformaldehyde (PFA) at room temperature for 15 min, washed twice with DPBS, and subjected to FV3000 confocal microscope imaging.

### Imaging of biotinylated proteins

A549 cells were seeded onto glass coverslips placed in 12-well plates and cultured overnight, followed by IDM incubation as indicated. On the next day, cells were pre-incubated with biotin–hydrazide (250 μM in DPBS) for 30 min, followed by green LED irradiation (525 nm) for the indicated durations (0, 10, 20, or 30 min). Cells were washed with DPBS and fixed. Fixed cells were permeabilized with 0.1% Triton X-100 in PBS at room temperature for 10 min, blocked with 2% BSA at room temperature for 1 h, and stained with streptavidin–Alexa Fluor 488 (Invitrogen; 1:3000 dilution in PBS) at room temperature for 1 h. After washing, samples were imaged using a FV3000 confocal microscope.

## Data availability

The mass spectrometry proteomics data generated in this study will be deposited in the PRIDE repository via the ProteomeXchange Consortium and will be made publicly available upon publication. The data supporting the findings of this study are available from the corresponding author upon reasonable request.

## Acknowledgments

This work was supported by the National Research Foundation of Korea (NRF) grant funded by the Korean government (MEST) (RS-2025-00520347 to T.-H.K. and RS-2026-25493835 to H.W.R) and by the InnoCORE program (GIST InnoCORE KH0830) of the Ministry of Science and ICT. S.N. was supported by the Korea Basic Science Institute (C612110, S610000). Instrumentation and technical support were provided by UNIST Central Research Facilities (UCRF).

## Author contribution

C.L., J.K.L., and C.Y. contributed equally to this work. All authors reviewed the manuscript. C.L. and T.-H. K. conceived this study and wrote the manuscript. C.L. and J.K.L. conducted mechanistic studies. J.K.L. synthesized all photocatalysts. J.K.L. designed and analyzed chemical experiments, including GC-MS analysis and DFT calculations. C.Y. performed and analyzed biological experiments, including proteomics sampling, confocal imaging, and immunoblotting. B.G.K. measured LC-MS/MS data for proteomics. G.Y. conducted ROS generation assay, EPR analysis, and cell viability test. J.K. performed the solvent-accessible surface area analysis. S.N. analyzed all the proteomics data. H.-W.R. contributed to study design, data interpretation, and critical revision of the manuscript. T.-H.K. supervised all aspects of this study.

