## Supplementary Figs 1-21, Supplementary Table 1 for "Water oxidation-driven histidine dioxidation enables probe-free proximity labeling"

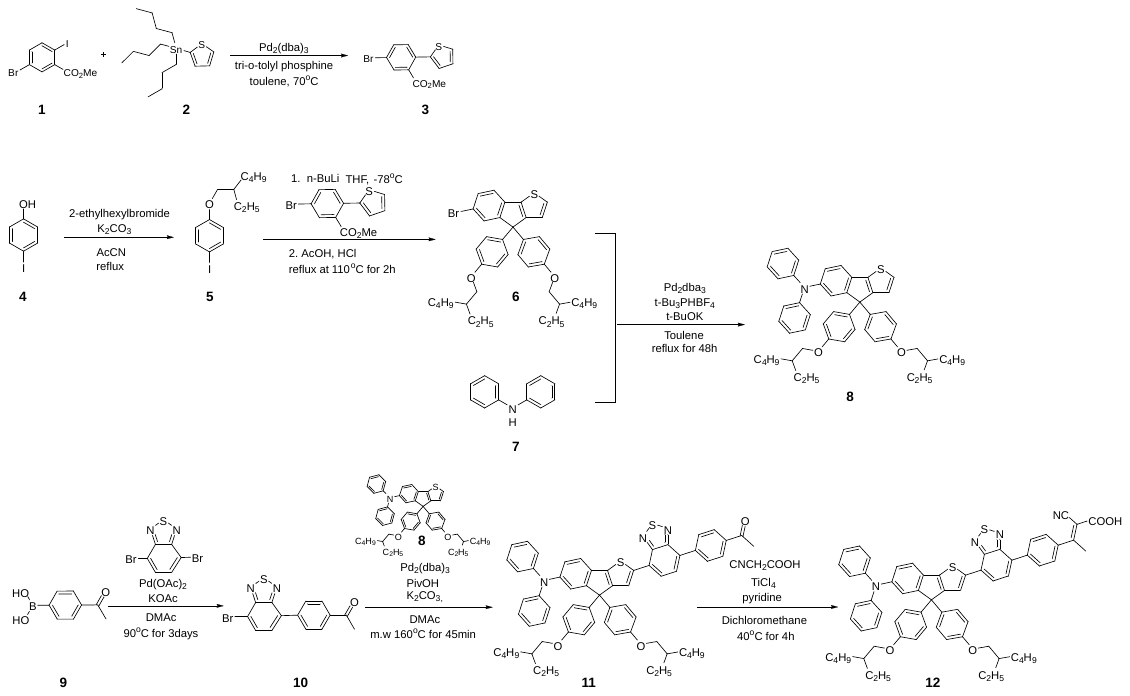

**Scheme 1.** Synthesis of IDM

Compounds TP-1, TP-2, and TP-3 were synthesized following reported procedures^1^. For IDM synthesis, compound 5 was prepared according to the literature methods^2^.

**Methyl 5-bromo-2-(thiophen-2-yl)benzoate (3)**

In a two-neck round-bottom flask, methyl 5-bromo-2-iodobenzoate (1.01 g, 2.97 mmol), tris(dibenzylideneacetone)dipalladium(0) (92 mg, 0.10 mmol), and tri-*o*-tolylphosphine (83 mg, 0.30 mmol) were added to toluene (15 mL) under a nitrogen atmosphere. After stirring for a few minutes, tributylstannyl thiophene (1.11 g, 2.97 mmol) was added, and the reaction mixture was refluxed at 110 °C overnight. Upon completion, the reaction mixture was worked up with dichloromethane (100 mL) and water. The organic layer was dried over anhydrous MgSO₄, filtered, and concentrated under reduced pressure using a rotary evaporator. The crude product was purified by column chromatography using a hexane/dichloromethane mixture as the eluent, affording compound 2 as a slightly yellow liquid. ^1^H NMR (400 MHz, CDCl_3_) *δ* (ppm): 7.86 (s, 1H), 7.62 (d, *J* = 8 Hz, 1H), 7.37 (dd, *J* = 4 Hz, 2H), 7.07­–7.02 (m, 2H), 3.75 (s, 3H). ^13^C NMR (400 MHz, CDCl_3_) δ (ppm): 167.59, 140.67, 133.95, 133.11, 132.57, 132.34, 132.30, 127.34, 126.60, 126.32, 121.60, 52.51.

**1-((2-ethylhexyl)oxy)-4-iodobenzene (5)**

Compound **4** and potassium bicarbonate were dissolved in degassed acetonitrile under nitrogen. Ethylhexylbromide was injected into the RBF. The solution was heated at 85 ^o^C for 16 h. The resulting mixture was extracted with dichloromethane and washed with brine. The organic layers were dried over MgSO_4_ and concentrated under reduced pressure using a rotary evaporator. The desired product was purified by column chromatography with 100% hexane. It is a transparent liquid. ^1^H NMR (400 MHz, DMSO-d_6_) *δ* (ppm): 7.53 (d, *J* = 8 Hz, 2H), 6.73 (d, *J* = 8 Hz, 2H), 3.77 (d, *J* = 8 Hz, 2H), 1.63–1.60 (m, 1H), 1.33–1.23 (m, 8H), 0.85–0.82 (m, 6H).

**6-bromo-4,4-bis(4-((2-ethylhexyl)oxy)phenyl)-4H-indeno[1,2-*b*]thiophene (6)**

Compound **5** (6.71 g, 20.19 mmol) was dissolved in argon-purged tetrahydrofuran under an inert condition in a round-bottom flask. At −78 ^o^C, n-butyllithium was slowly injected into the solution. After 30 minutes, compound **3** (2 g, 6.73 mmol) dissolved in THF was slowly injected into the reaction mixture. The temperature was gradually increased to room temperature overnight. Upon completion, it was quenched by adding water, and the resulting mixture was extracted with dichloromethane and washed with brine. The organic layers were dried over MgSO_4_ and concentrated under reduced pressure using a rotary evaporator. These crude products were used as reactants, dissolving in degassed acetic acid. The 1 mL of hydrogen chloride was injected dropwise, and the temperature was elevated to 110 °C. The reaction mixture was refluxed overnight. After the reaction, the process of quenching and extraction was identically conducted. The desired product was purified by column chromatography using a n-hexane/dichloromethane (2/1, v/v) eluent. The transparent liquid product was obtained (3.23 g, 72.7%). ^1^H NMR (400 MHz, CDCl_3_) δ (ppm): 7.44 (d, *J* = 3.2 Hz, 1H), 7.41 (dd, *J* = 8 Hz, 1H), 7.32 (d, *J* = 4 Hz, 1H), 7.31 (d, *J* = 8 Hz, 1H), 7.09 (d, *J* = 8 Hz, 4H), 6.98 (d, *J* = 8 Hz, 1H), 6.77 (d, *J* = 8 Hz, 4H), 3.79 (d, *J* = 8 Hz, 4H), 1.70–1.67 (m, 2H), 1.31–1.29 (m, 8H), 0.92–0.88 (m, 12H). ^13^C NMR (400 MHz, DMSO-d_6_) δ (ppm): 158.22, 156.91, 156.19, 139.09, 136.19, 135.89, 130.95, 129.19, 128.88, 123.59, 121.65, 118.83, 114.76, 78.18, 70.14, 62.19, 39.08, 30.36, 28.86, 23.73, 22.95, 14.38, 11.33. FTIR (neat, cm−1): 2956, 2925, 2858, 1607, 1507, 1457, 1291, 1246, 1176, 1113, 1033, 822, 731, 662, 523. HRMS (LC/Q-TOF) calc. for C_39_H_46_BrO_2_S = 657.2407. Found: m/z =657.2407 [(M+H)^+^]. Mass error ≈ 0 ppm.

**4, 4-bis(4-((2-ethylhexyl)oxy)phenyl)-N, N-diphenyl-4H-indeno[1,2-b]thiophen-6-amine (8)**

In the glove box, Pd2(dba)3 (3 mg, 0.003 mmol), tri-tert-butylphosphonium tetrafluoroborate (2 mg, 0.007 mmol), and potassium tert-butoxide (112 mg, 1.0 mmol) were prepared in a two-neck round-bottom flask and sealed with two rubber septa. Compound **6** (57 mg, 0.33 mmol) and compound **7** (200mg, 0.30 mmol) were dissolved in degassed anhydrous toluene and injected into the round-bottom flask. The reaction mixture was refluxed for 48 hours at 110 °C. Upon completion, it was cooled down to room temperature and stirred for 30 minutes. The resulting mixture was extracted with dichloromethane and washed with brine. The organic layers were dried over MgSO_4_ and concentrated under reduced pressure using a rotary evaporator. The crude product was purified by column chromatography using a n-hexane/dichloromethane (9/1, v/v) eluent. The slightly yellow liquid product was obtained (210 mg, 92.5%). ^1^H NMR (400 MHz, Acetone-d6) δ (ppm): 7.44 (s, 1H), 7.42 (d, *J* = 4 Hz, 1H), 7.28 (t, *J* = 8 Hz, 4H), 7.12 (d, 1H), 7.10 (d, *J* = 4 Hz, 1H), 7.06–6.99 (m, 10H), 6.96 (dd, *J* = 4 Hz, 1H), 6.80 (d, *J* = 8 Hz, 4H), 3.84 (d, *J* = 4 Hz, 4H), 1.70–1.67 (m, 2H), 1.33–1.31 (m, 8H), 0.93–0.87 (m, 12H). ^13^C NMR (400 MHz, Acetone-d6) δ (ppm): 209.10, 158.22, 155.84, 155.61, 147.72, 145.77, 140.19, 136.48, 131.89, 129.29, 128.73, 127.62, 123.86, 123.21, 122.95, 122.84, 122.44, 119.81, 114.08, 69.94, 39.35, 30.38, 28.89, 23.66, 22.83, 13.50, 10.56. FTIR (neat, cm−1): 3035, 2957, 2926, 2858, 2227, 1695, 1587, 1506, 1488, 1470, 1420, 1378, 1342, 1276, 1246, 1176, 1154, 1031, 821, 752, 697, 661, 628, 521, 413. HRMS (LC/Q-TOF) calc. for C_51_H_58_NO_2_S = 748.4188. Found: m/z = 748.4183 [(M+H)^+^]. Mass error ≈ −0.7 ppm.

**4-(7-bromobenzo[*c*][1,2,5]thiadiazol-4-yl)benzaldehyde (10)**

All reactants Pd_2_(OAc) (90 mg, 0.4 mmol), 4,7-dibromobenzo[c][1,2,5]thiadiazole (6 g, 20.42 mmol), (4-formylphenyl)boronic acid (compound **9**, 2.24 g, 13.62 mmol), and potassium acetate (2 g, 20.42 mmol) were added in a round-bottom flask. Anhydrous dimethylacetamide was injected, and the temperature was increased to 90 °C. After 72 hours, the reaction mixture was cooled to room temperature and quenched with water. The resulting mixture was extracted with sufficient dichloromethane and washed with brine. The organic layers were dried over MgSO_4_ and concentrated under reduced pressure using a rotary evaporator. The crude product was purified by column chromatography using a n-hexane/dichloromethane (1/2, v/v) eluent. The product was obtained as a yellow solid after trituration with dichloromethane and n-hexane, yielding 3.16 g (69.6 %). ^1^H-NMR (400 MHz, CDCl_3_) *δ* (ppm): 8.21 (d, *J* = 4 Hz, 2H), 8.19(d, *J* = 4 Hz, 2H), 8.10 (d, *J* = 8 Hz, 1H), 8.05 (d, *J* = 8 Hz, 1H) 2.75 (s, 3H). ^13^C-NMR (400 MHz, CDCl_3_) *δ* (ppm): 197.63, 153.85, 152.82, 141.06, 136.79, 132.66, 132.16, 129.34, 128.74, 128.67, 114.31, 26.75. FTIR (neat, cm^−1^): 3081.81, 2961.05, 2922.56, 2351.62, 1730.94, 1671.06, 1601.25, 1553.27, 1526.38, 1504.68, 1478.99, 1434.41, 1410.42, 1361.86, 1355.13, 1327.86, 1269.46, 1191.26, 1151.34, 1118.19, 1015.42, 961.20, 941.20, 933.24, 888.74, 853.04, 837.55, 826.18, 785.92, 767.65, 735.88, 649.15, 628.56, 622.09, 600.17, 590.78, 558.54, 514.42, 507.00, 501.77, 487.89, 460.01. HRMS (LC/Q-TOF) calc. for C­_14_H_9_BrN_2_­OS= 331.9619. Found: *m/z* =331.9621 [(M+H)^+^]. Mass error = −0.7 ppm.

**4-(7-(6-(diphenylamino)-4,4-bis(4-((2-ethylhexyl)oxy)phenyl)-4H-indeno[1,2-*b*]thiophen-2-yl)benzo[*c*][1,2,5]thiadiazol-4-yl)benzaldehyde (11)**

All reactants, compound **8** (477 mg, 0.64 mmol), compound **10** (213 mg, 0.64 mmol), Pd_2_(dba)_3_ (30 mg, 0.032 mmol), Pivalic acid (65 mg, 0.64 mmol), and K_2_CO_3_ (310 mg, 2.24 mmol) were prepared with argon-purged anhydrous N, N-dimethylacetamide in a microwave tube under inert conditions. The mixture was reacted at 160 °C for 45 minutes using a microwave. Upon completion, the mixture was extracted with dichloromethane and washed with brine. The organic layers were dried over MgSO_4_ and concentrated under reduced pressure using a rotary evaporator. The crude product was purified by column chromatography using a n-hexane/dichloromethane (1/1, v/v) eluent. The product was obtained as a violet solid after trituration with dichloromethane and methanol, yielding 180 mg (24.1%). ^1^H NMR (400 MHz, Acetone-d_6_) δ (ppm): 8.29 (s, 1H), 8.20–8.17 (m, 2H), 8.15–8.12 (m, 3H), 7.96 (d, *J* = 8 Hz, 1H), 7.56–7.54 (d, *J* = 8 Hz, 1H), 7.31 (t, *J* = 8 Hz, 4H), 7.16–7.13 (m, 5H), 7.08–7.02 (m, 6H), 7.00 (dd, *J* = 4 Hz, 1H), 3.85 (d, *J* = 8 Hz, 4H), 2.64 (s, 3H), 1.69–1.66 (m, 2H), 1.44–1.28 (m, 16H), 0.91–0.86 (m, 12H). ^13^C NMR (400 MHz, Acetone-d6) δ (ppm): 209.09, 159.35, 156.42, 155.73, 152.20, 147.58, 146.57, 141.41, 141.19, 136.20, 131.24, 130.55, 129.37, 129.25, 128.98, 128.85, 128.34, 127.49, 124.63, 124.20, 123.87, 123.17, 122.69, 120.43, 114.22, 69.97, 68.31, 39.34, 25.95, 23.65, 22.81, 13.47, 10.54. FTIR (neat, cm−1): 2957, 2926, 2858, 2228, 1682, 1603, 1541, 1507, 1491, 1470, 1420, 1407, 1357, 1265, 1247, 1177, 1117, 1031, 898, 875, 825, 753, 697, 677, 629, 602, 526, 486, 474, 453, 443, 427, 421, 411. HRMS (LC/Q-TOF) calc. for C_65_H_66_N_3_O_3_S_2_ = 1000.4540. Found: m/z = 1004.4533 [(M+H)^+^]. Mass error ≈ 0.7 ppm.

**E-2-cyano-3-(4-(7-(6-(diphenylamino)4,4-bis(4-((2-ethylhexyl)oxy)phenyl)-4H-indeno[1,2-*b*]thiophen-2-yl)benzo[*c*][1,2,5]thiadiazol-4-yl)phenyl)but-2-enoic acid (12, IDM)**

Compound **11** (70 mg, 0.07 mmol) and cyanoacetic acid (120 mg, 1.4 mmol) were dissolved in argon-purged dichloromethane under an inert condition in a round-bottom flask. The solution was cooled to 0 °C, and titanium tetrachloride (0.15 mL, 1.4 mmol) was slowly added. After stirring for 30 minutes, pyridine (0.11 mL, 1.4 mmol) was injected, and the mixture was stirred for another 30 minutes. The reaction temperature was then gradually raised to 40 °C and maintained for 4 hours. Upon completion, the mixture was cooled to room temperature and quenched with water. The organic layer was extracted with ethyl acetate, washed with brine, and dried over MgSO_4_. It was concentrated under reduced pressure using a rotary evaporator. The crude product was purified by column chromatography using a dichloromethane/methanol (10/1, v/v) eluent. The IDM was obtained as a dark pink solid after trituration with ethyl acetate and methanol, yielding 44 mg (62.9 %). ^1^H NMR (400 MHz, CDCl_3_/DMSO-d_6_ (*v/v* = 1:1)) δ (ppm): 8.10 (s, 1H), 8.00 (d, *J* = 8 Hz, 2H), 7.81 (d, *J* = 8 Hz, 1H), 7.53 (d, *J* = 8 Hz, 2H), 7.39 (d, *J* = 8 Hz, 2H), 7.20–7.16 (m, 4H), 7.03–7.01 (d, *J* = 8 Hz, 4H), 6.98–6.93 (m, 7H), 6.90–6.88 (d, *J* = 8 Hz, 1H), 6.71 (d, *J* = 8 Hfz, 4H), 3.72 (d, *J* = 4 Hz, 4H), 2.50 (s, 3H), 1.59–1.57 (m, 2H), 1.35–1.18 (m, 16H), 0.84–0.78 (m, 12H). ^13^C NMR (400 MHz, CDCl_3_/DMSO-d_6_ (*v/v* = 1:1)) δ (ppm): 158.10, 158.08, 156.17, 155.40, 153.63, 152.22, 147.37, 147.35, 146.29, 142.37, 142.28, 141.32, 136.15, 136.1, 131.09, 129.46, 129.40, 129.01, 128.87, 128.61, 127.83, 127.64, 126.76, 126.00, 124.20, 123.23, 121.74, 120.56, 119.45, 118.19, 114.35, 79.11, 70.14, 62.34, 39.18, 30.39, 28.94, 23.95, 22.95, 14.25, 11.25. FTIR (neat, cm−1): 3421, 2956, 2926, 2858, 2252, 2127, 1589, 1540, 1505, 1488, 1470, 1382, 1282, 1245, 1176, 1116, 1051, 1024, 1006, 898, 875, 823, 757, 697, 676, 630, 567, 526, 485, 454, 427, 421, 417, 411. HRMS (LC/Q-TOF) calc. for C_67_H_67_N_4_O_2_S_2_ = 1023.4700. Found: m/z = 1023.4707 [(M−CO_2_+H)^+^]. Mass error ≈ 0.7 ppm.

**Characterization of compound 3**

**^1^H NMR**

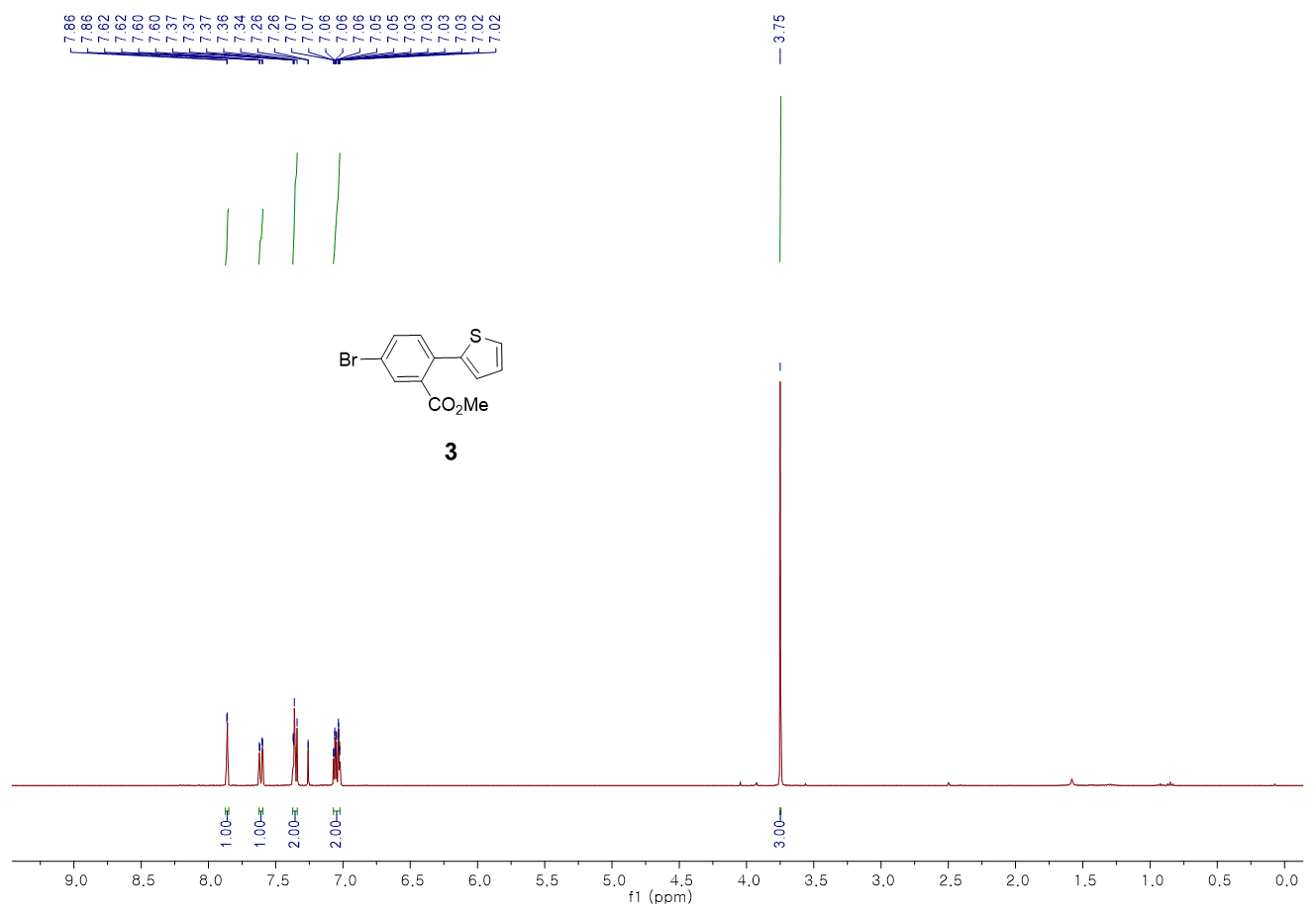

**^13^C NMR**

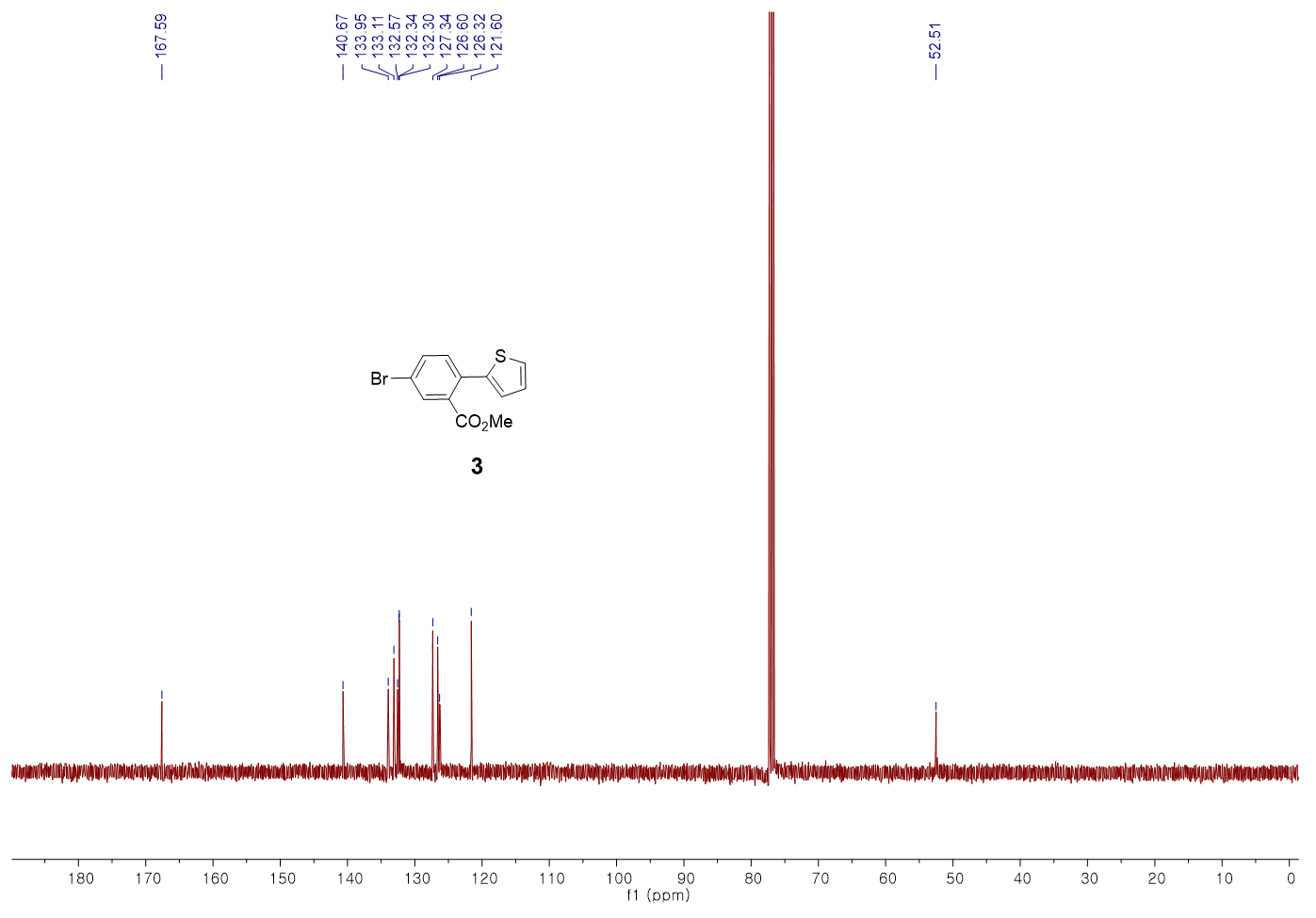

**Characterization of compound 6**

**^1^H NMR**

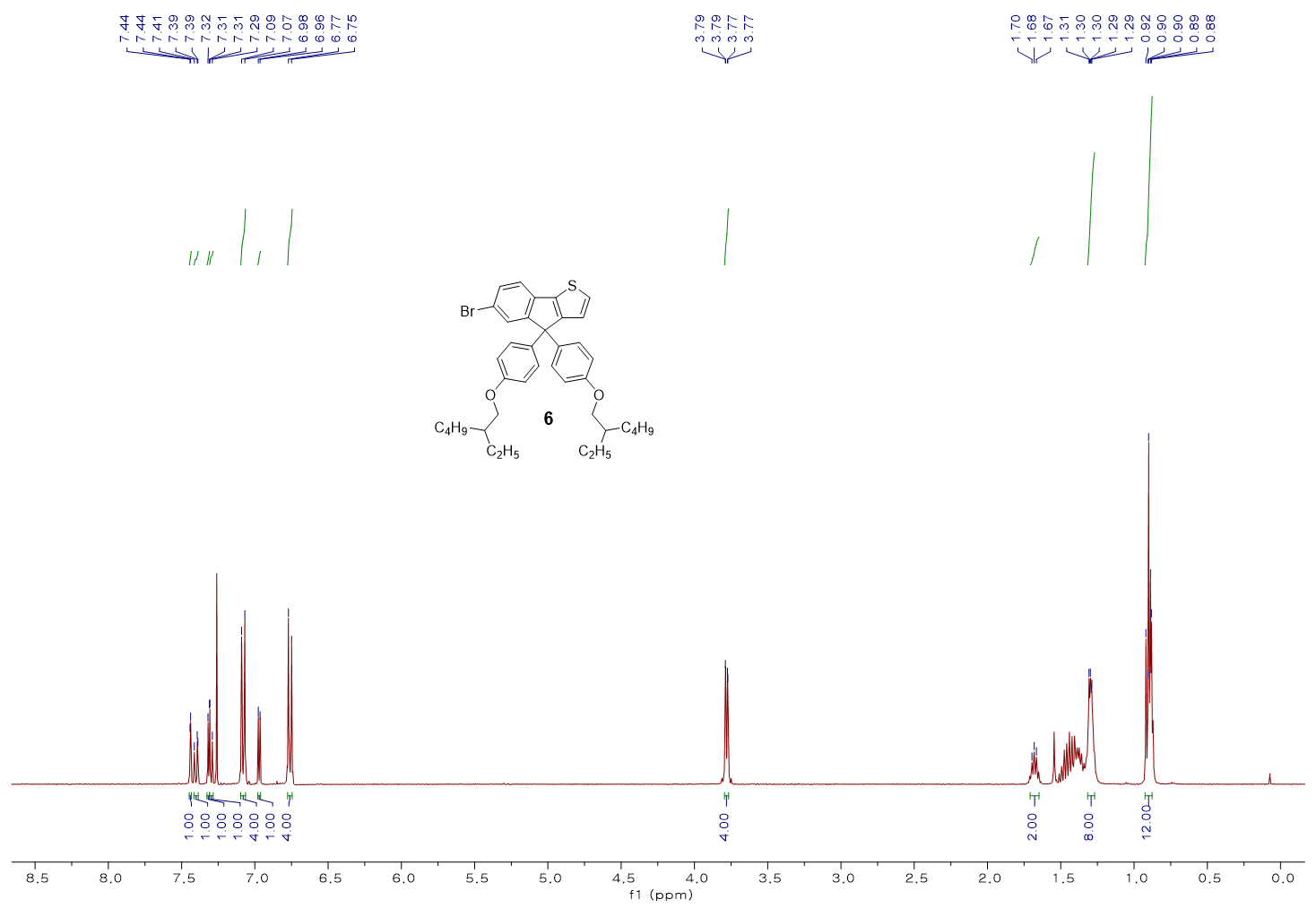

**^13^C NMR**

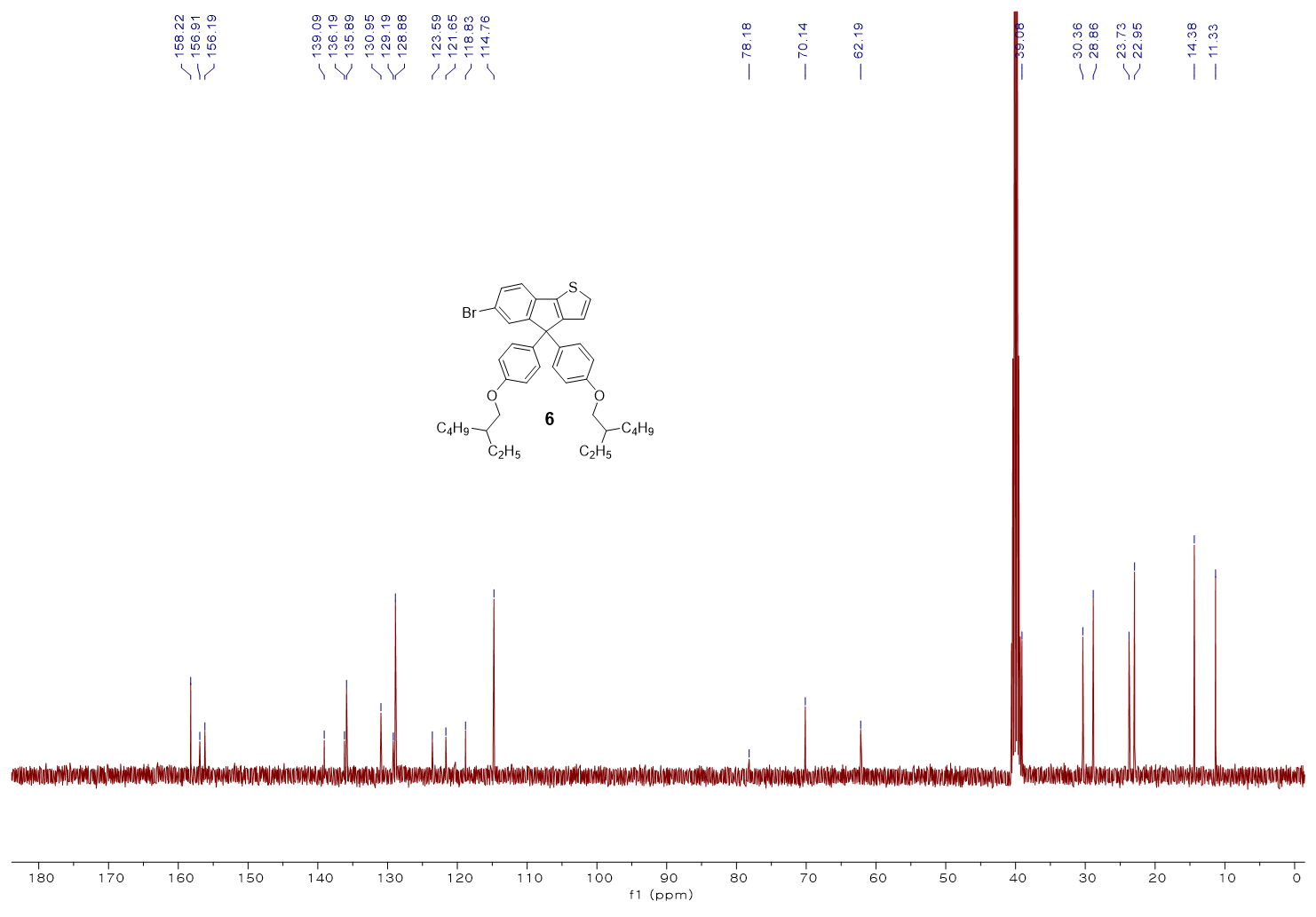

**FT-IR**

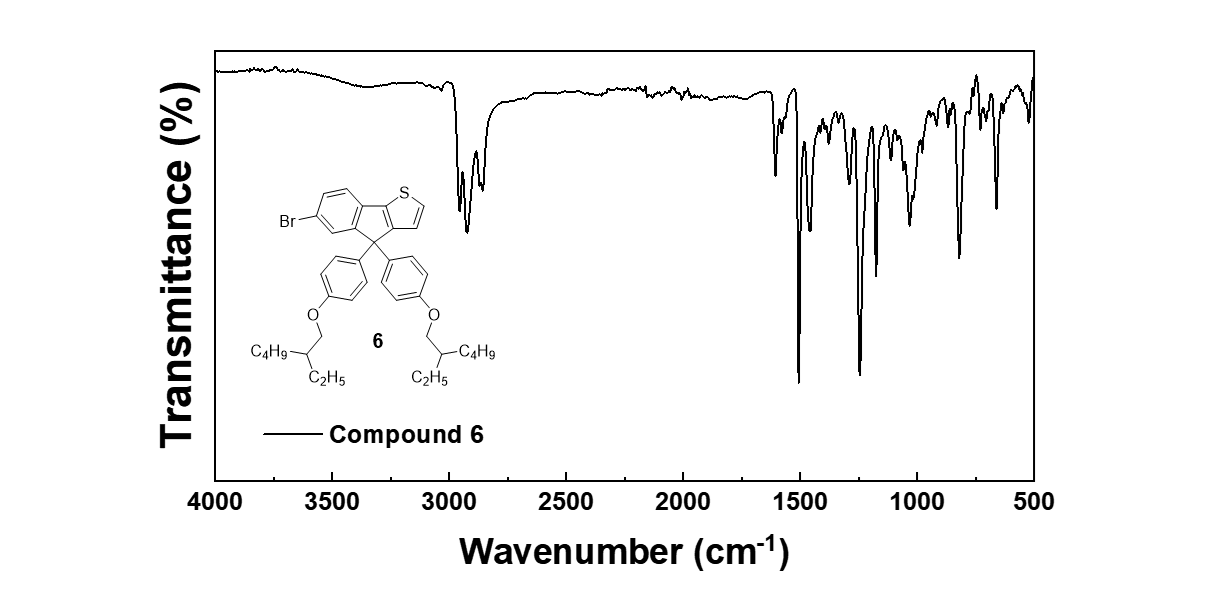

**HRMS**

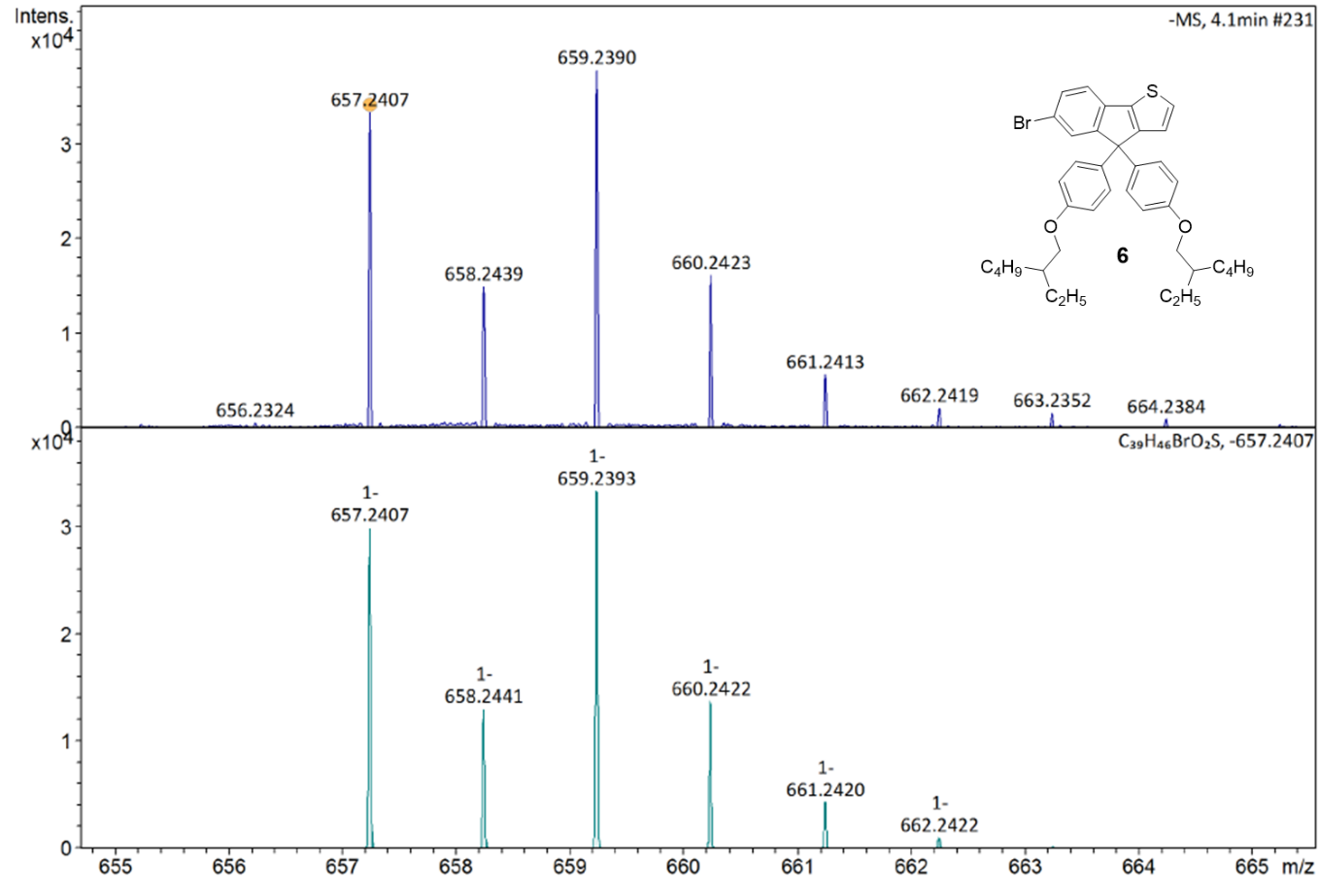

**Characterization of compound 8**

**^1^H NMR**

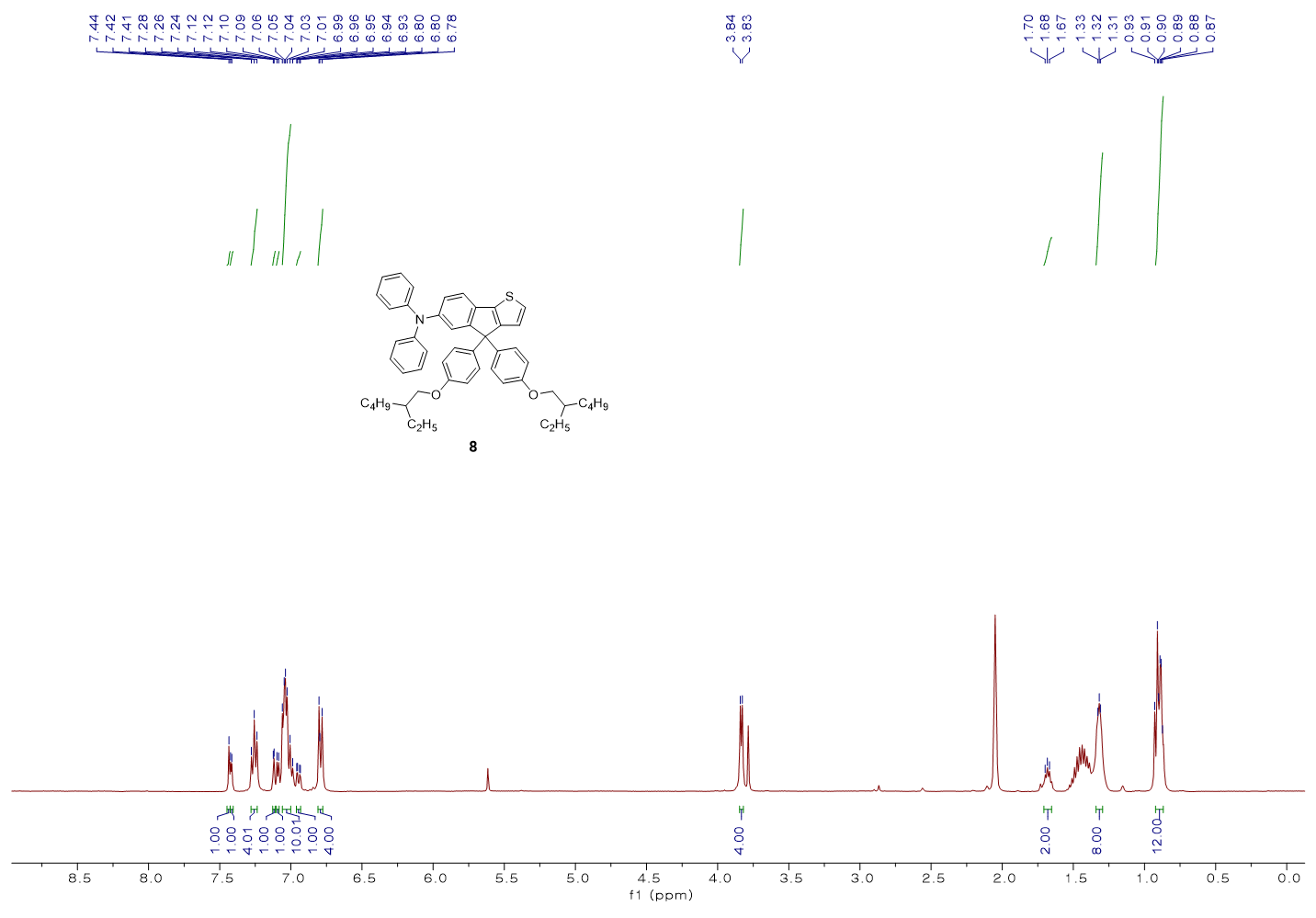

**^13^C NMR**

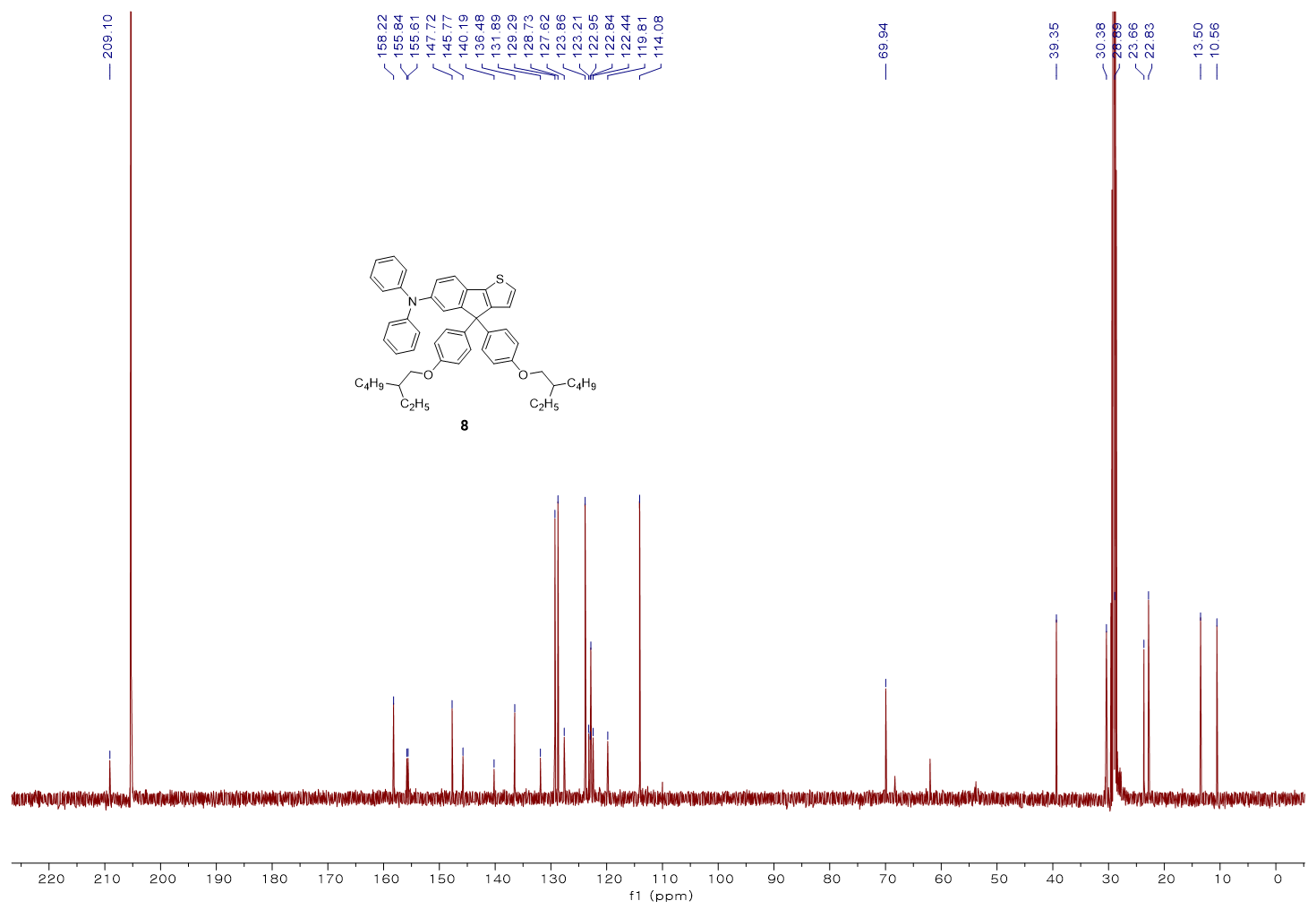

**FT-IR**

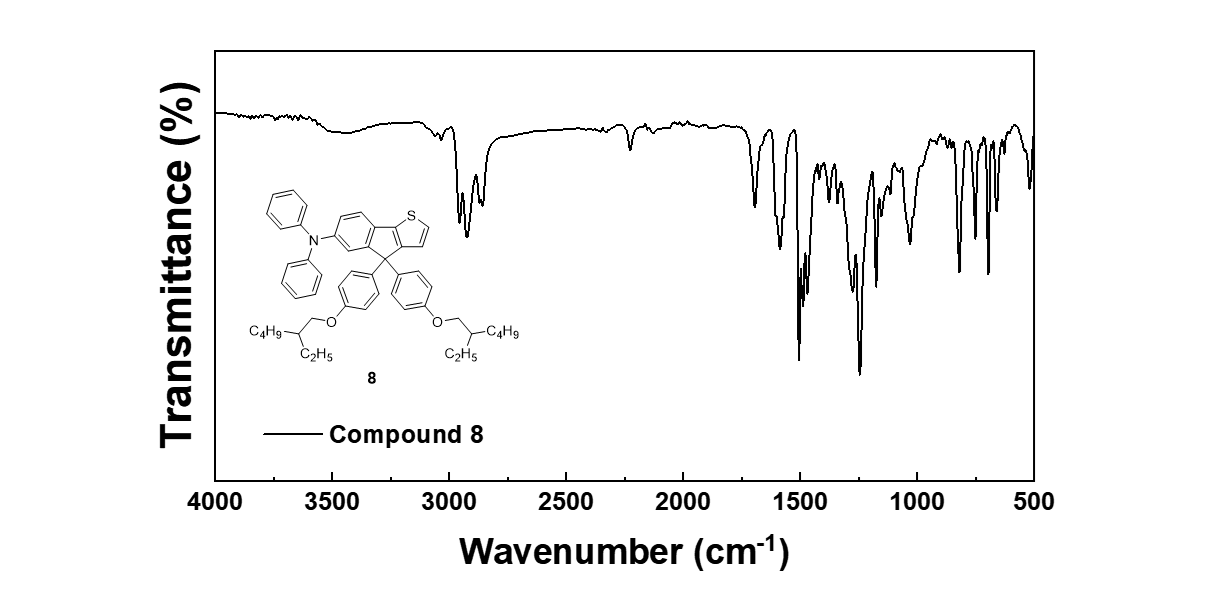

**HRMS**

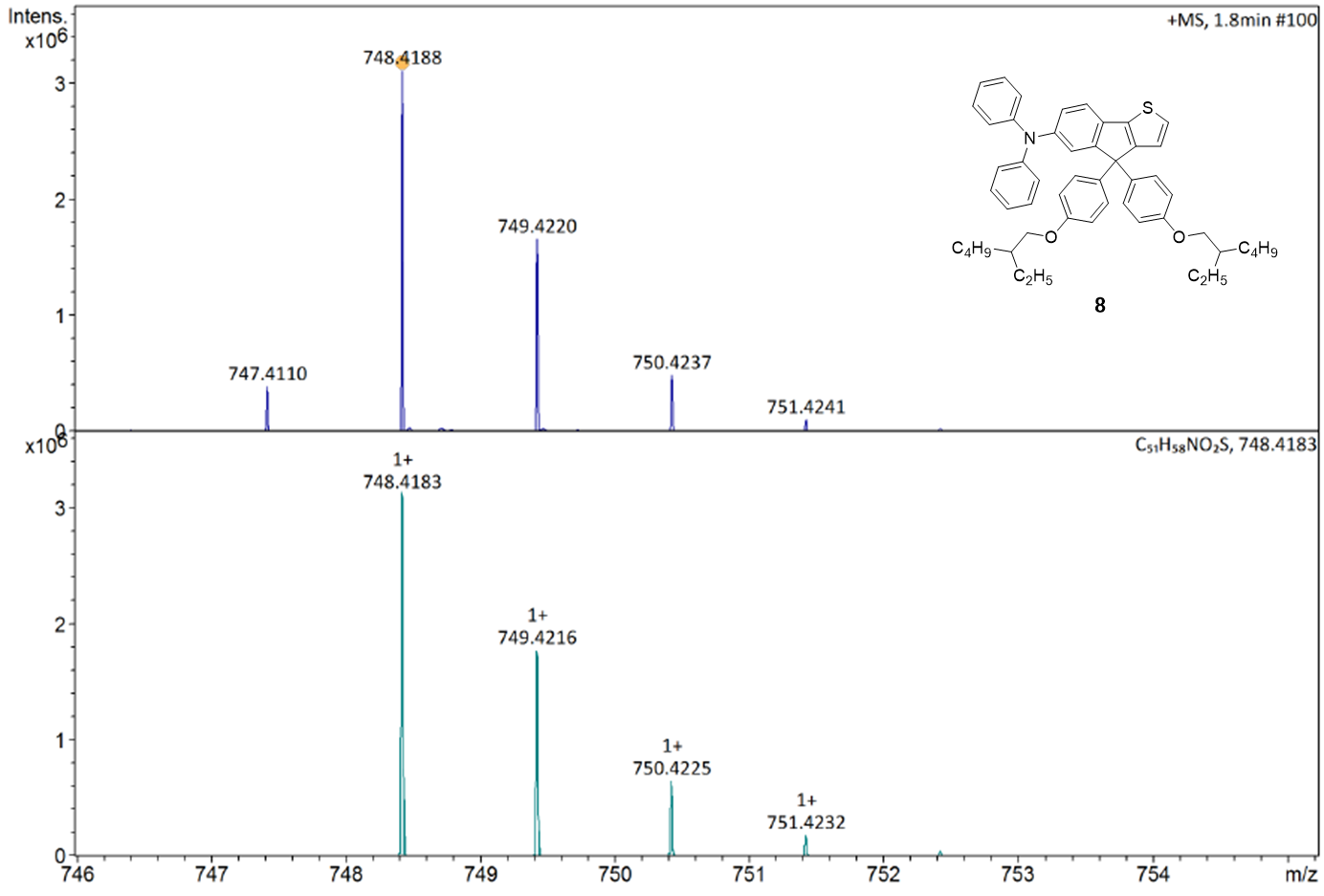

**Characterization of compound 10**

**^1^H NMR**

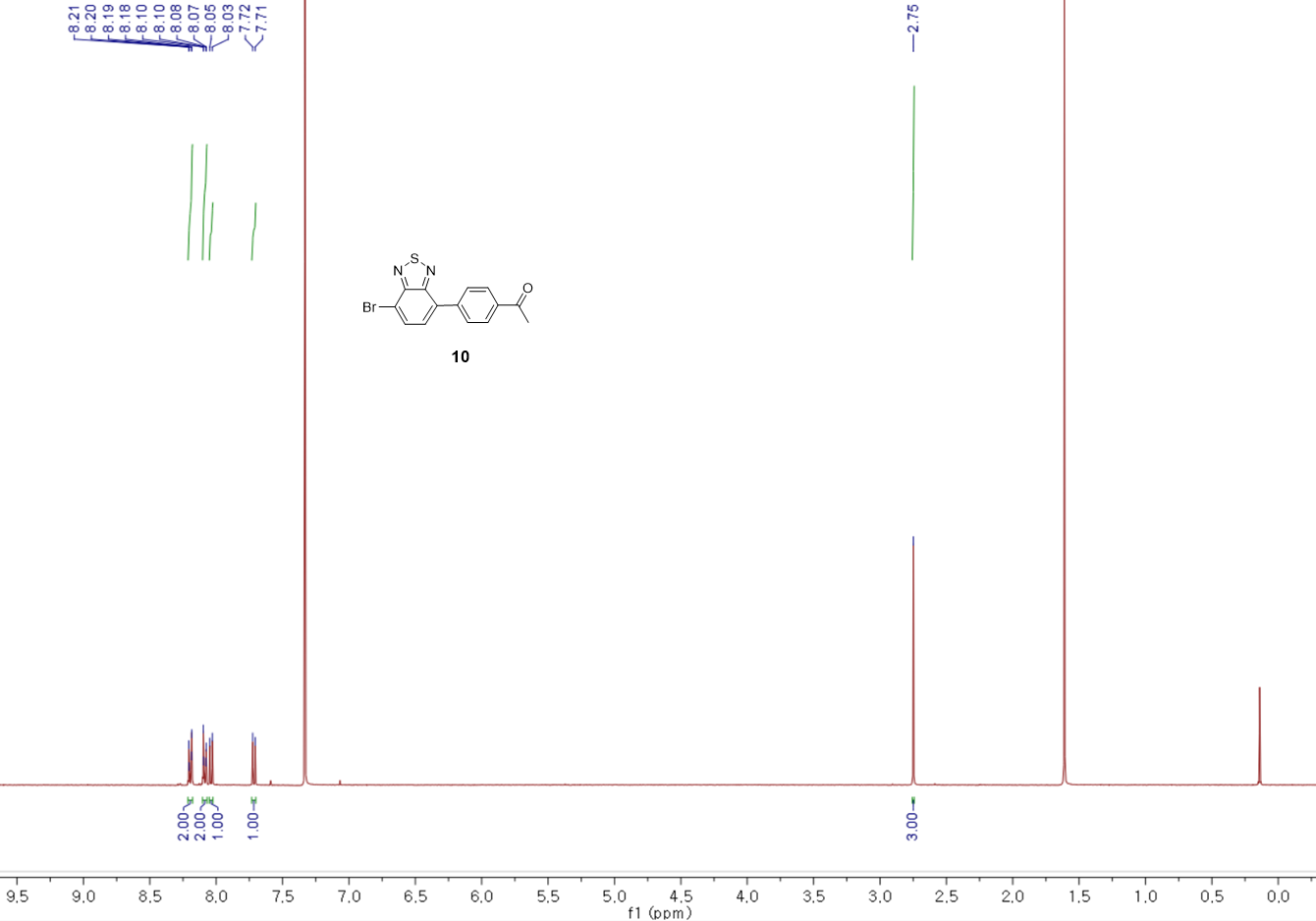

**^13^C NMR**

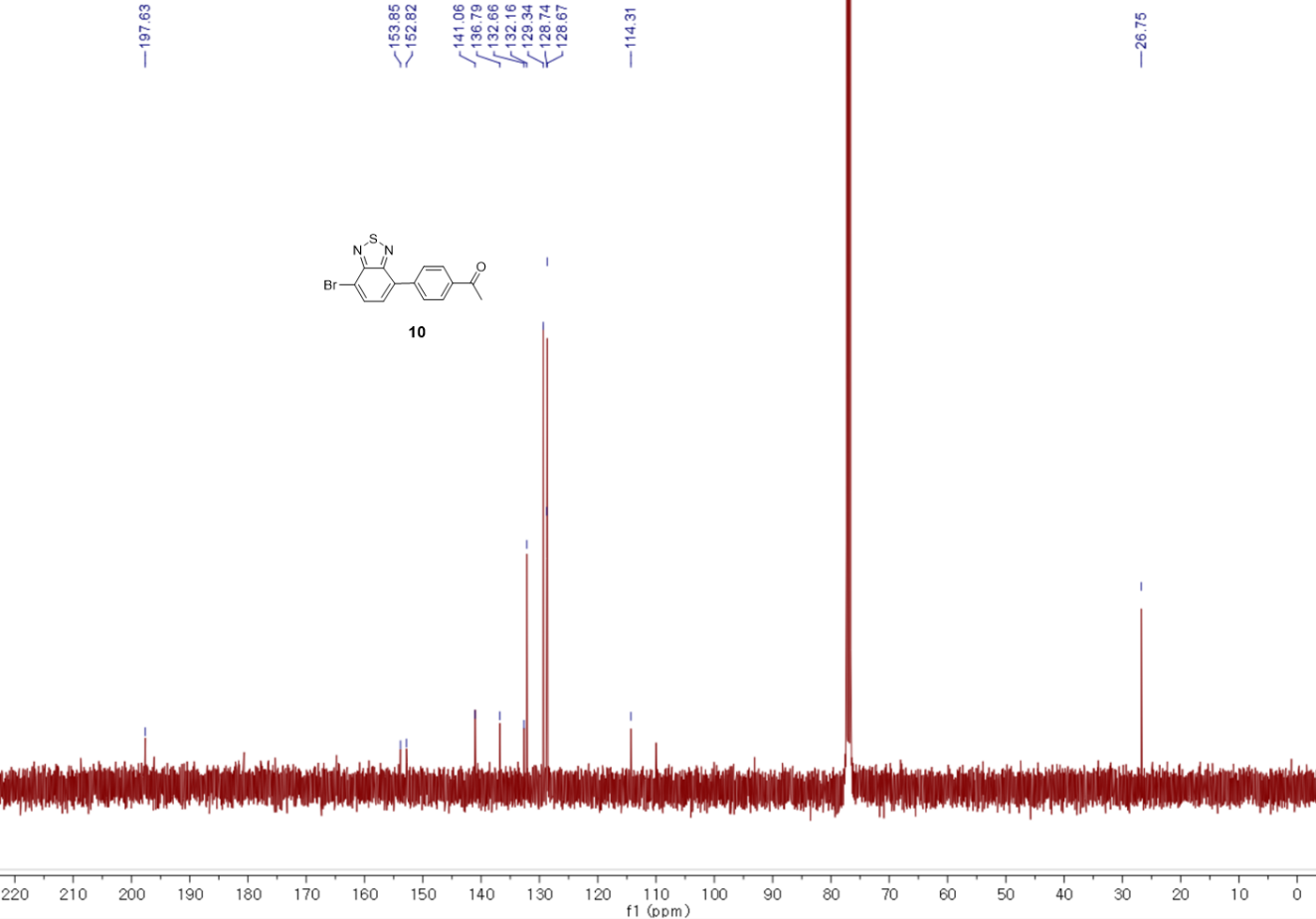

**FT-IR**

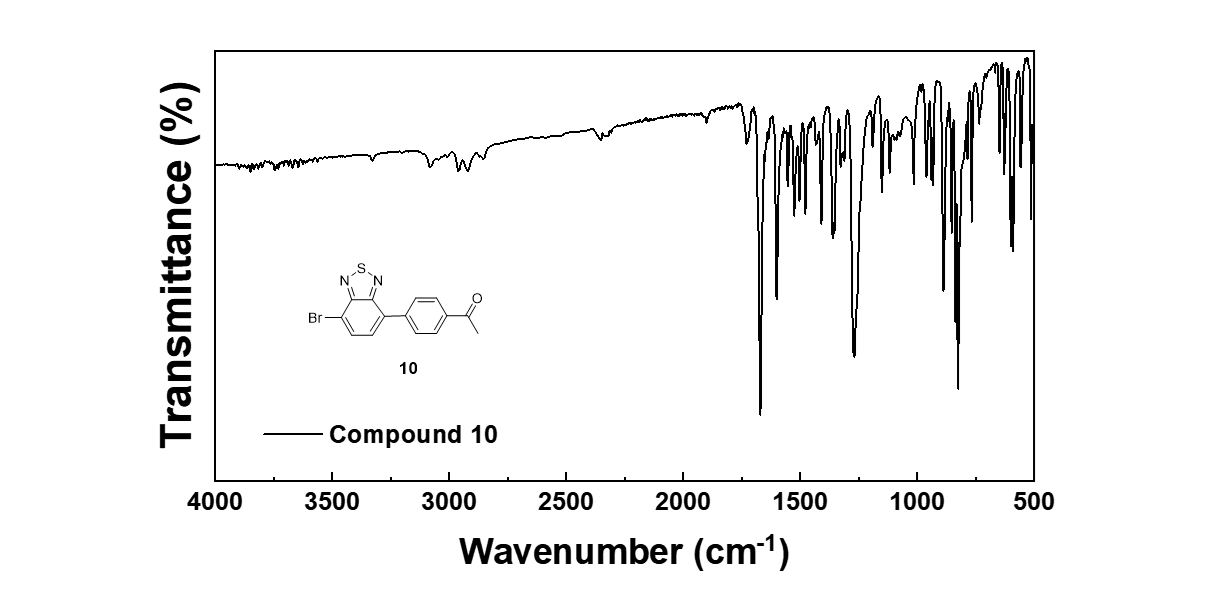

**HRMS**

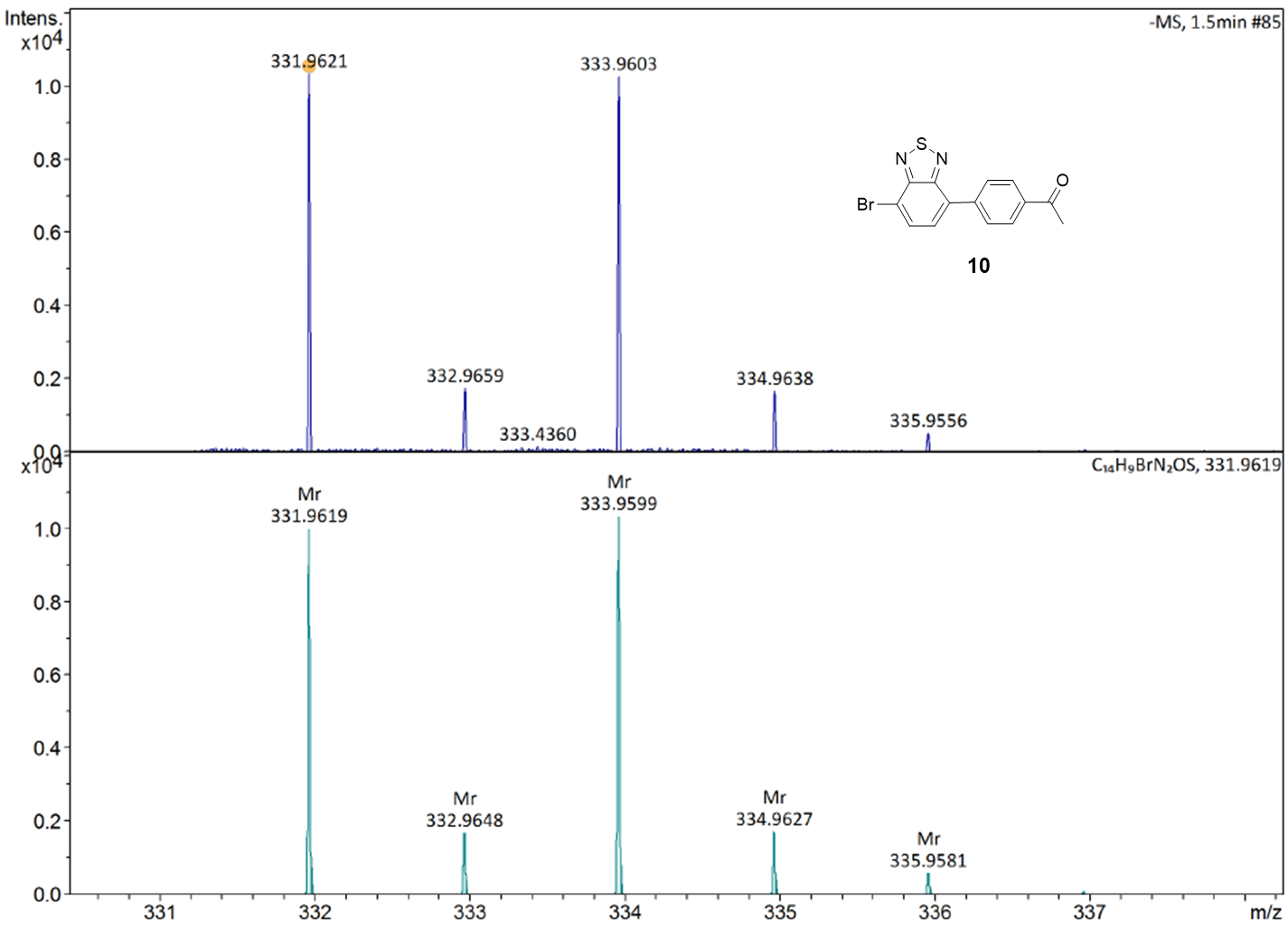

**Characterization of compound 11**

**^1^H NMR**

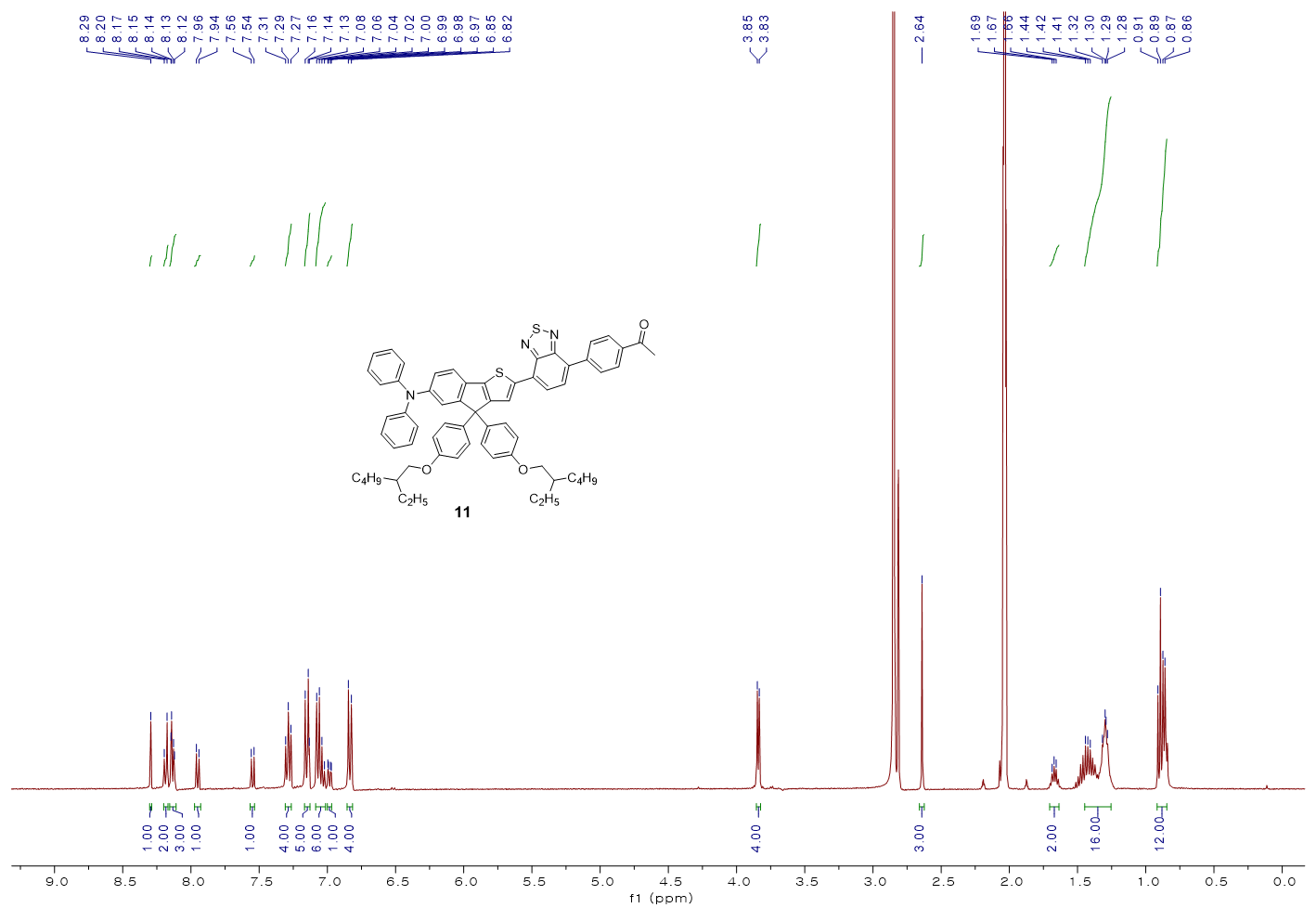

**^13^C NMR**

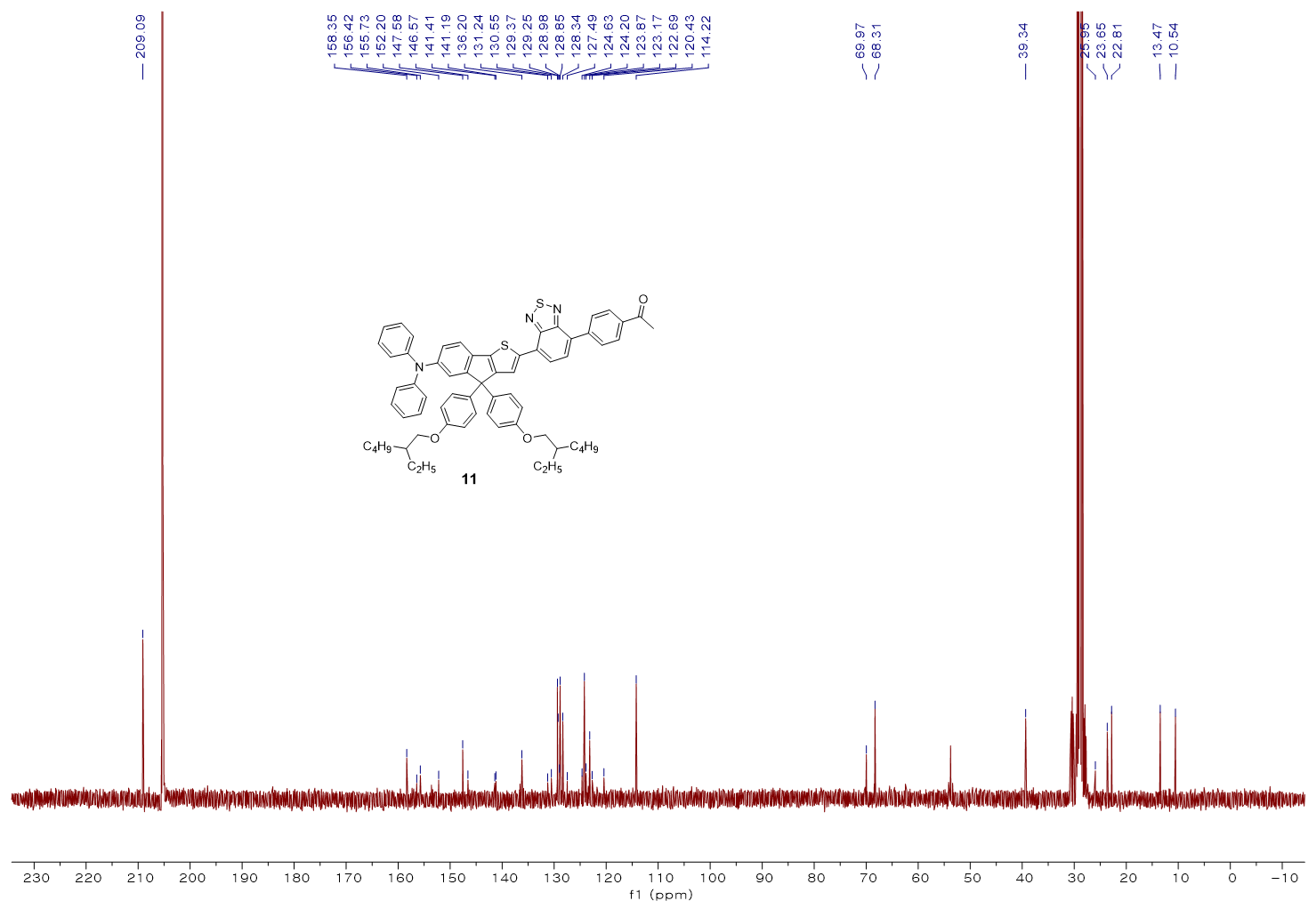

**FT-IR**

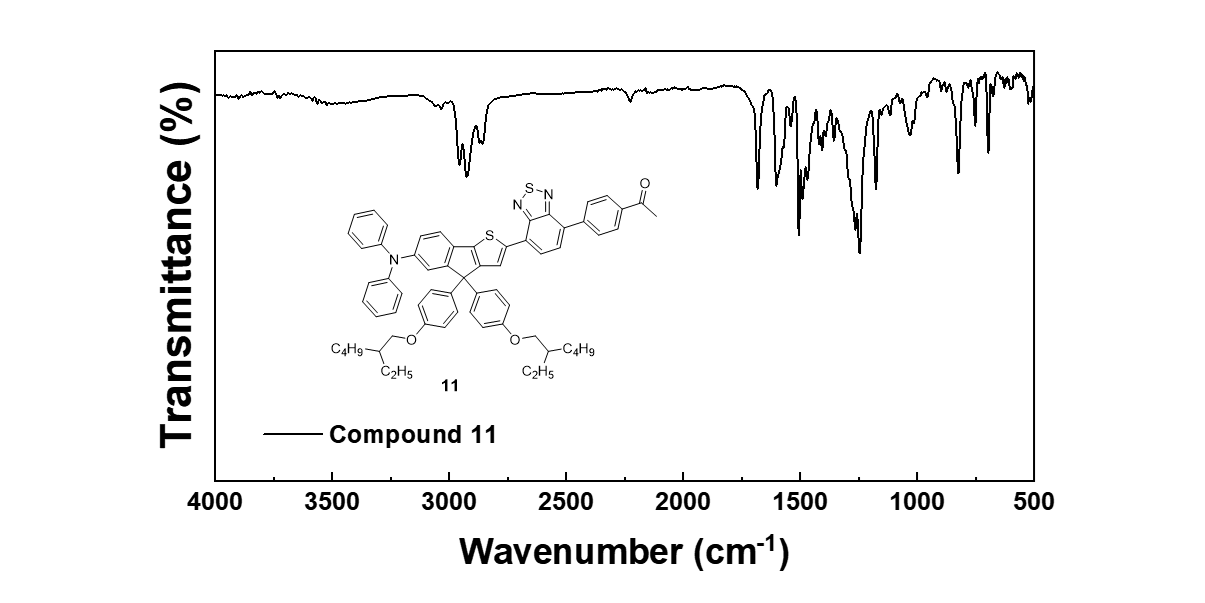

**HRMS**

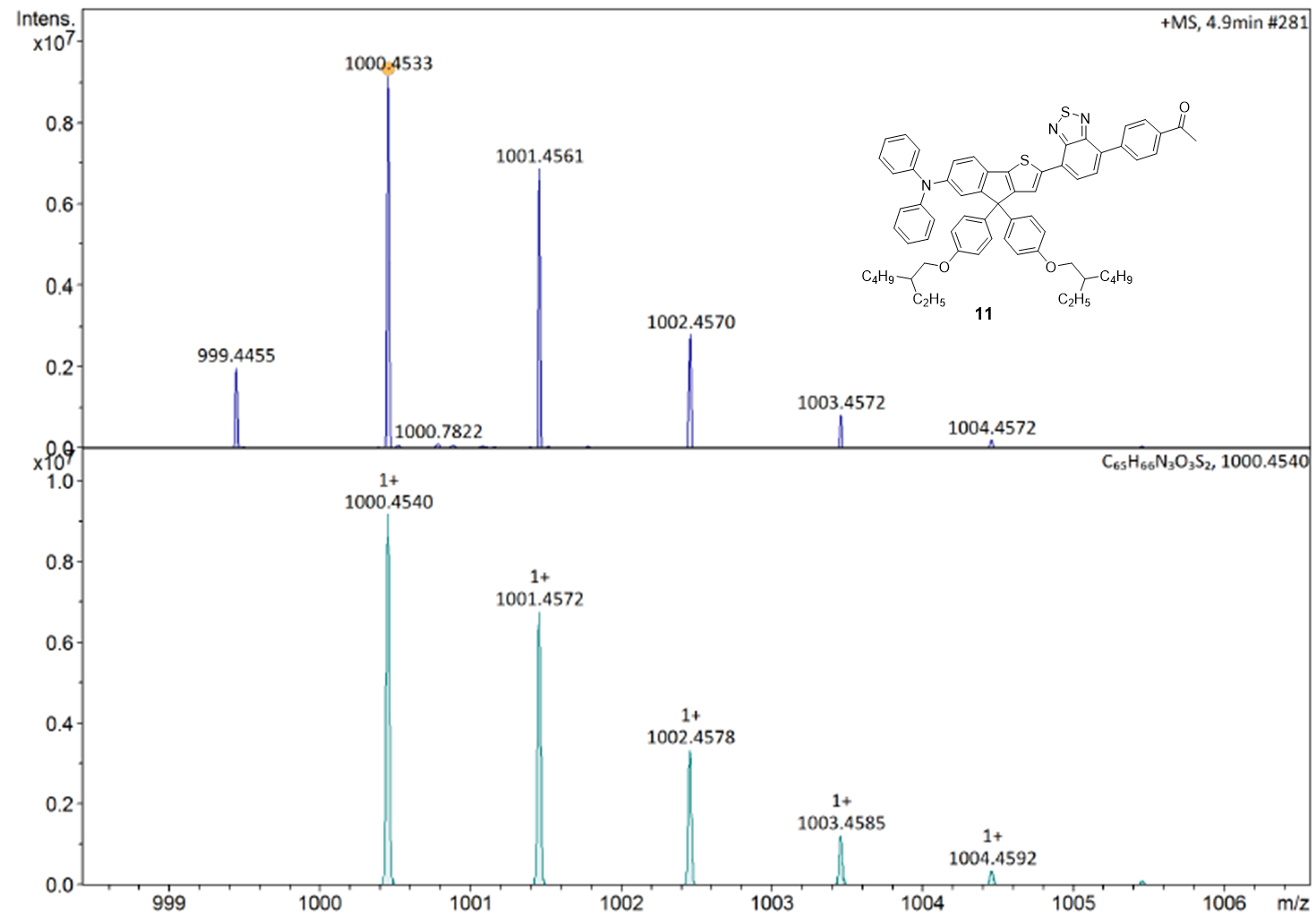

**Characterization of compound 12**

**^1^H NMR**

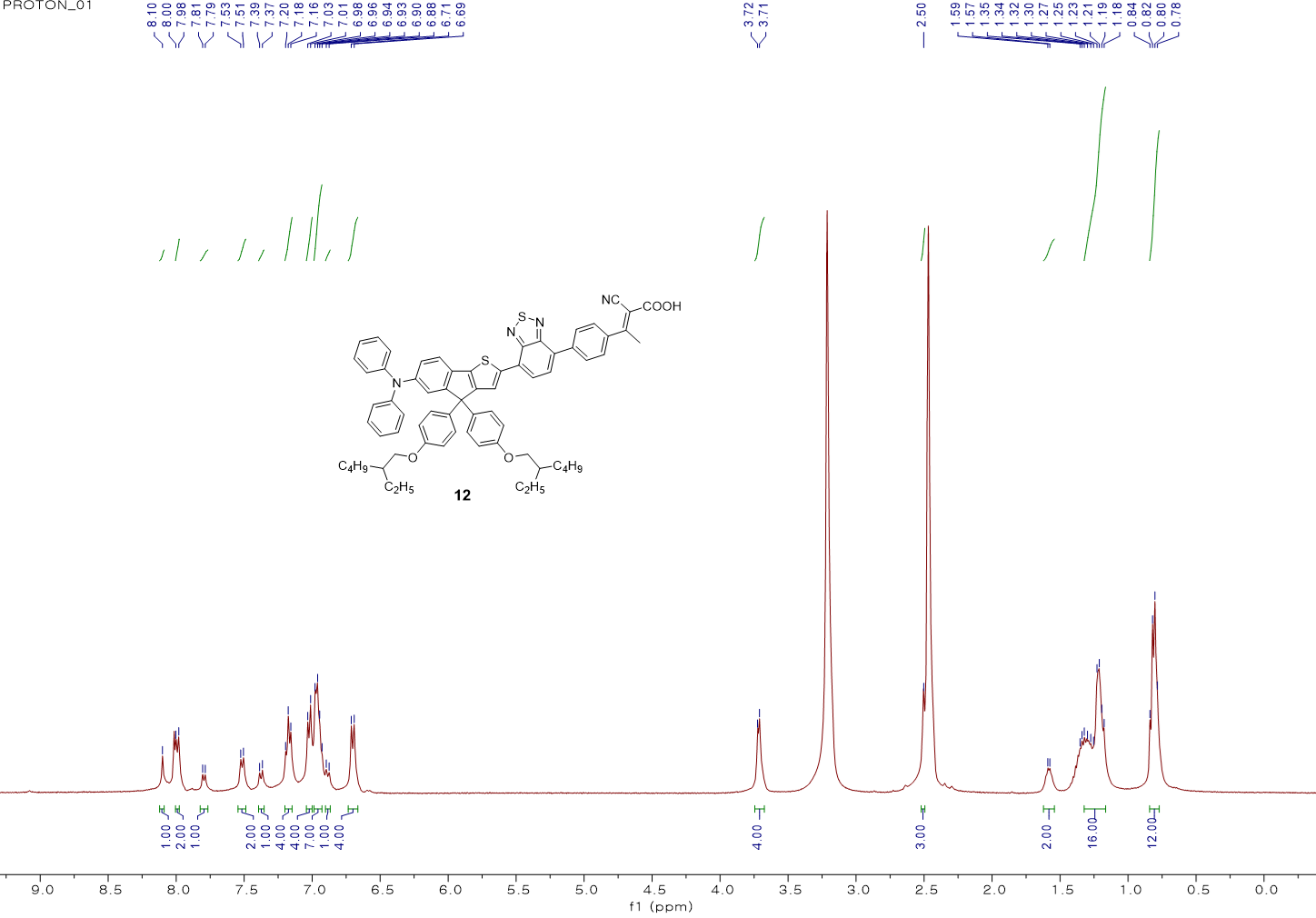

**^13^C NMR**

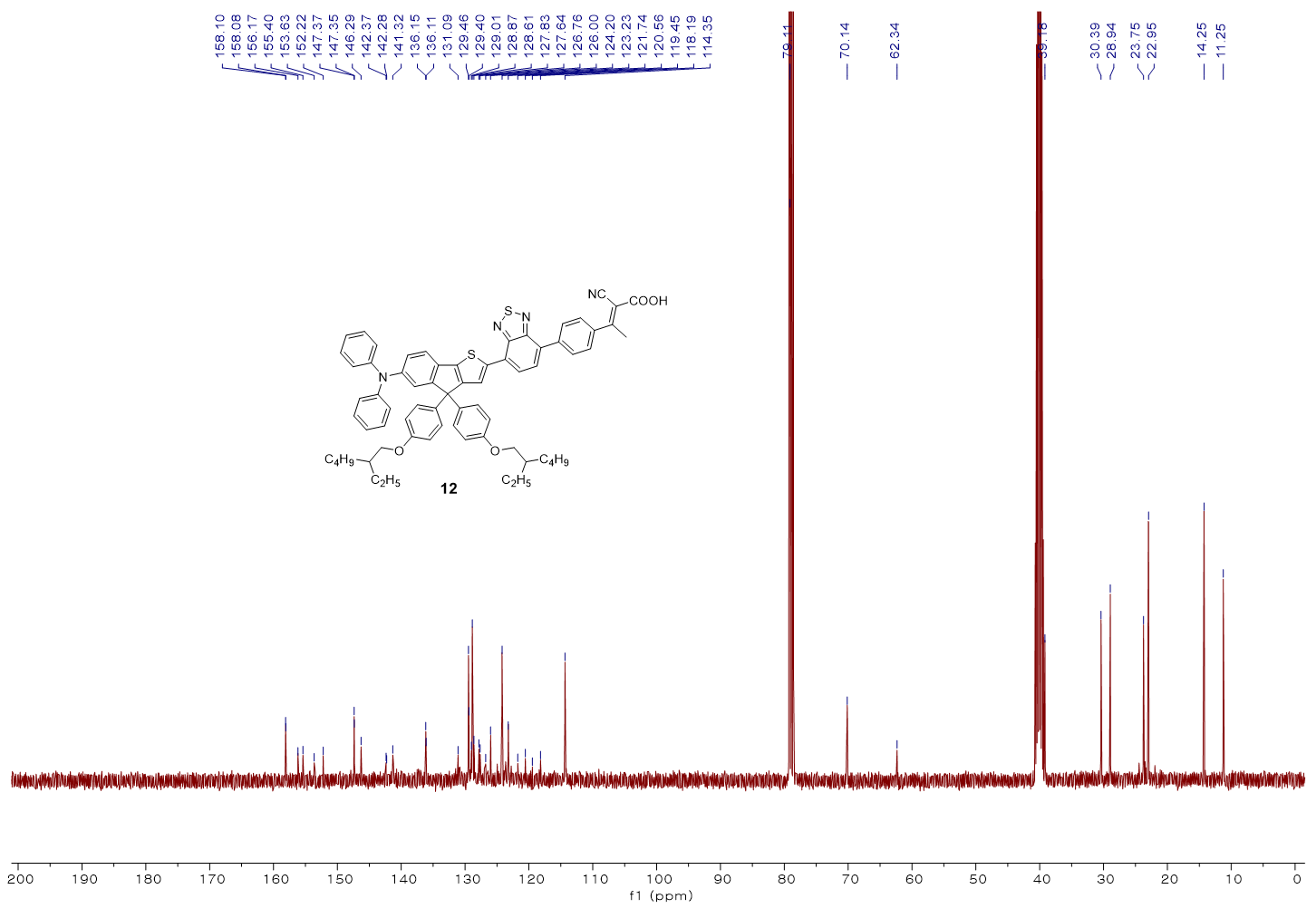

**FT-IR**

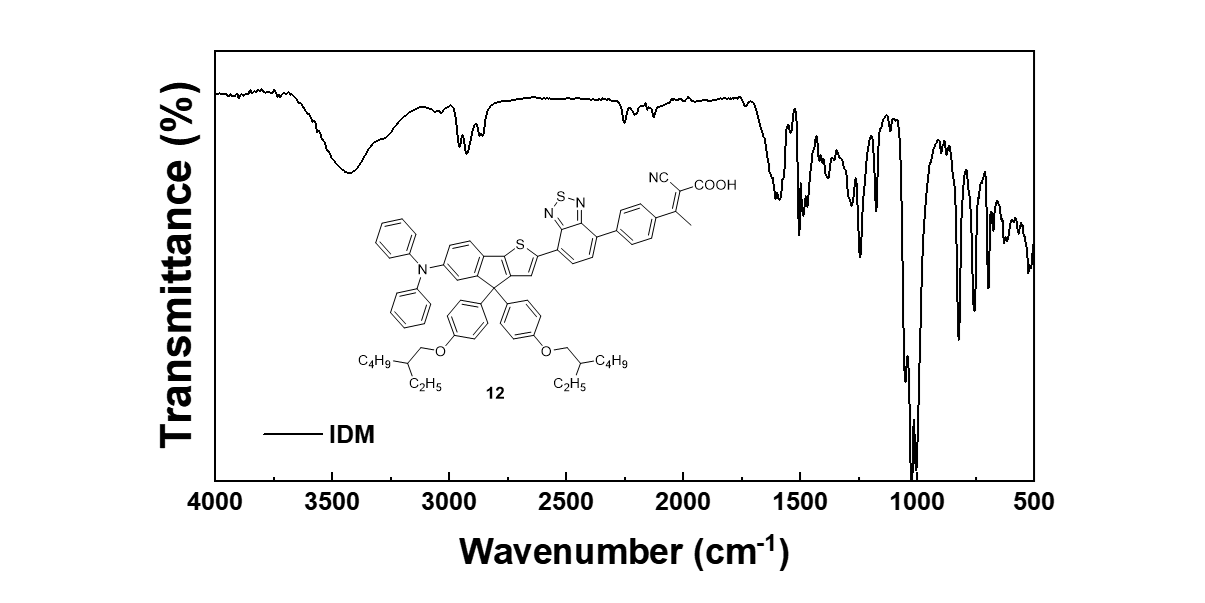

**HRMS**

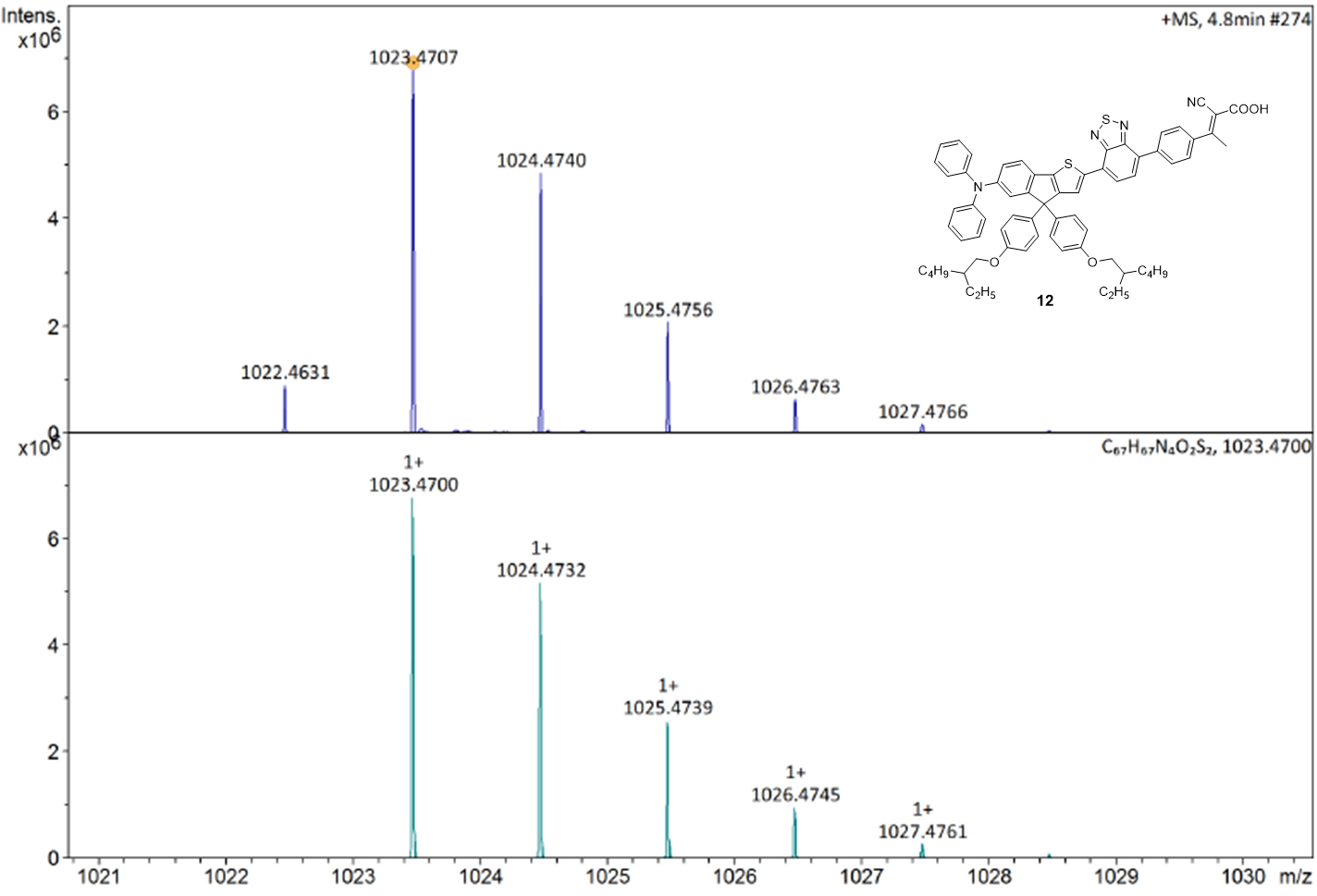

**Supplementary table**

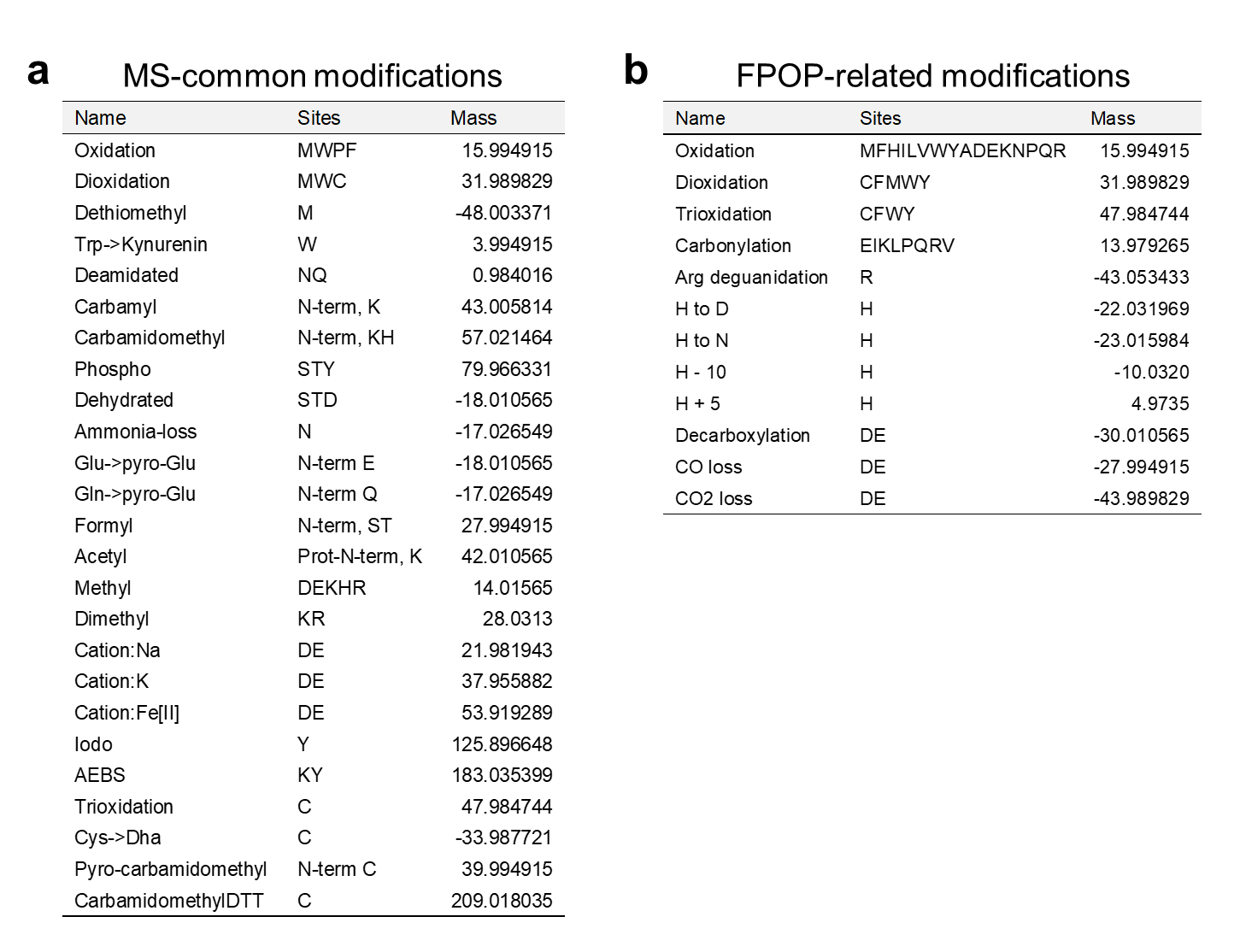

**Supplementary Table 1. List of modifications considered in the global oxidative modification search. a** MS-common modifications provided by MODplus. **b** Modifications caused by cell FPOP. The overlapping ones between **a** and **b** were only considered once (as FPOP-related modifications) during the search.

**Supplementary figures**

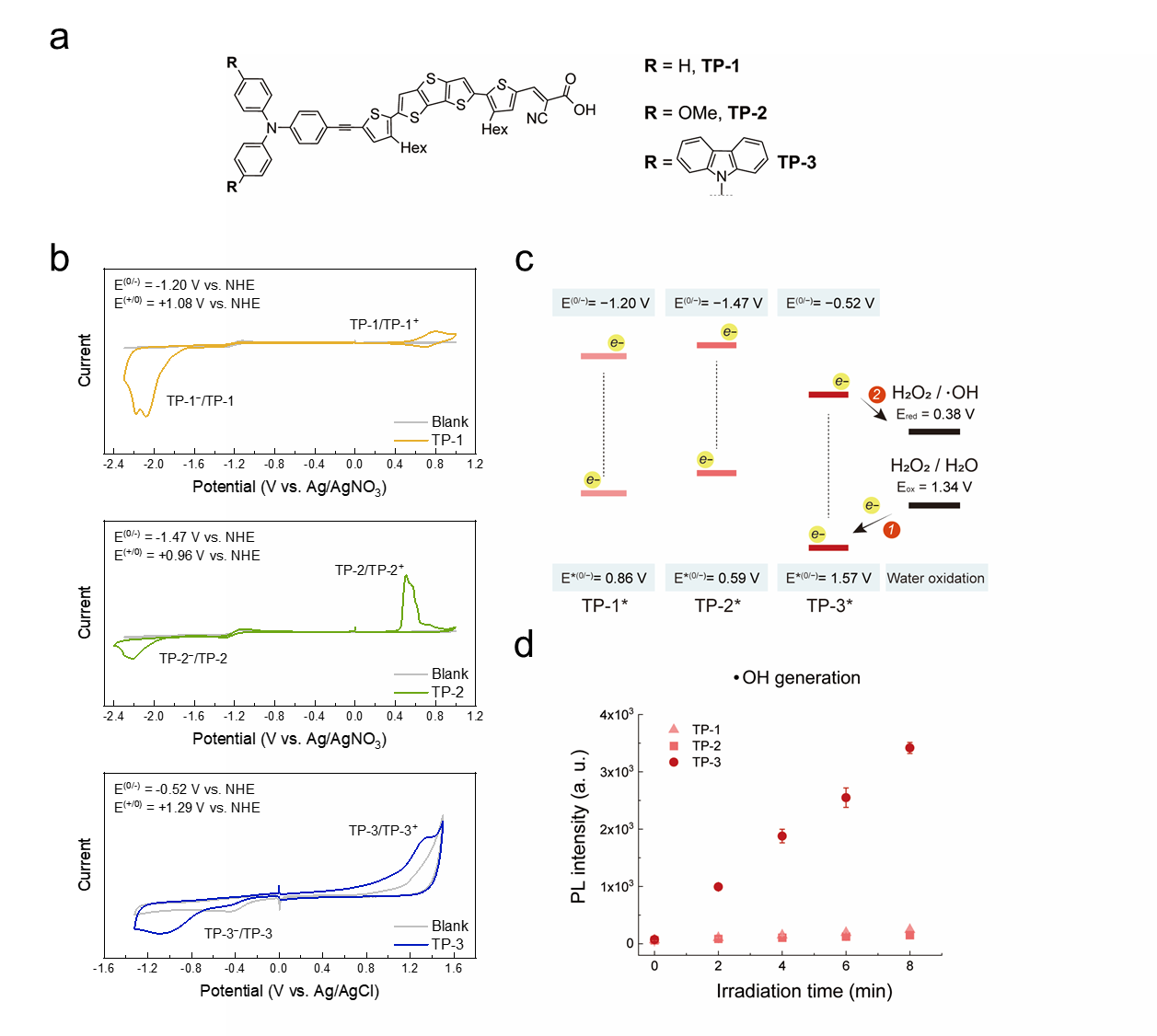

**Supplementary Fig. 1. Redox potentials and photocatalytic •OH generation of TP-1, TP-2, and TP3. a** Molecular structure of TP-1, TP-2, and TP-3. **b** Cyclic voltammetry for measuring ground-state redox potentials of TP-1, TP-2, and TP-3. **c** Energy diagram of electron transfer for water oxidation-driven hydroxyl radical generation. The energy levels were described vs. NHE (at pH 7.4). Excited-state redox potentials were calculated from the ground-state redox potentials. **d** •OH production was measured by an HPF assay using photocatalysts (5 µM) and HPF (10 µM) under green LED irradiation for 0, 2, 4, 6, and 8 minutes (λmax = 525 nm, 16.6 mW·cm−2).

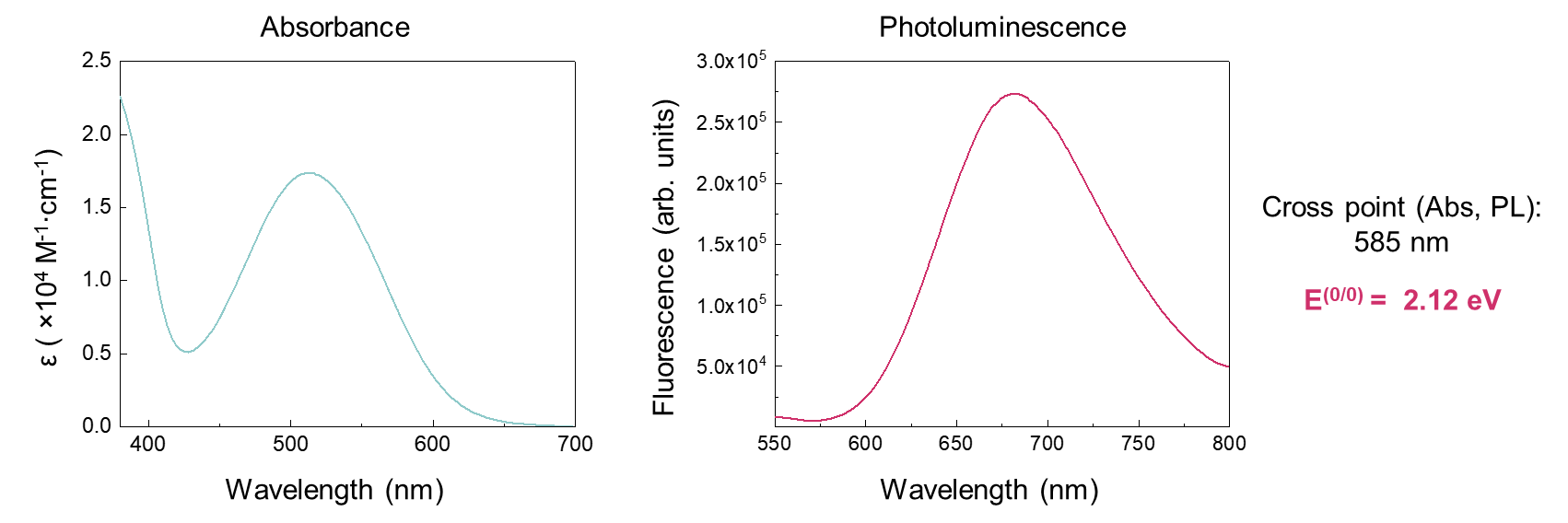

**Supplementary Fig. 2. Photophysical properties of IDM.** Absorbance and Photoluminescence of IDM. E^(0/0)^ was calculated from the cross point of the absorbance and photoluminescence spectra.

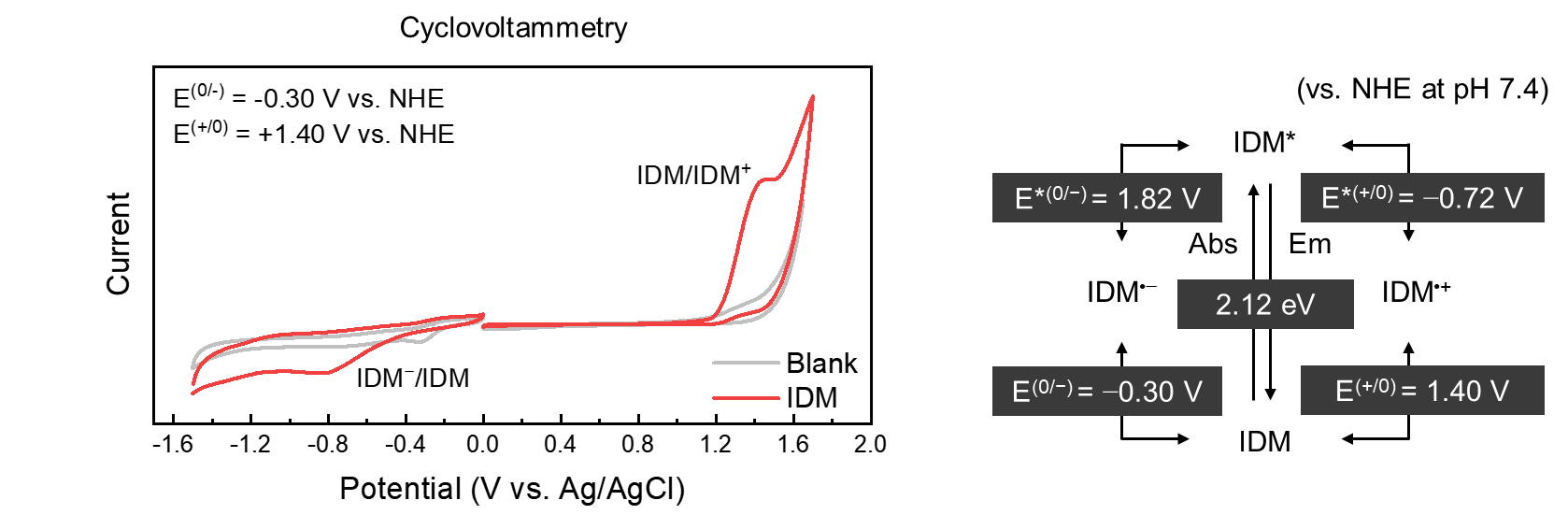

**Supplementary Fig. 3. Redox potentials of IDM and excited IDM.** Cyclic voltammetry curves of IDM and corresponding redox potentials.

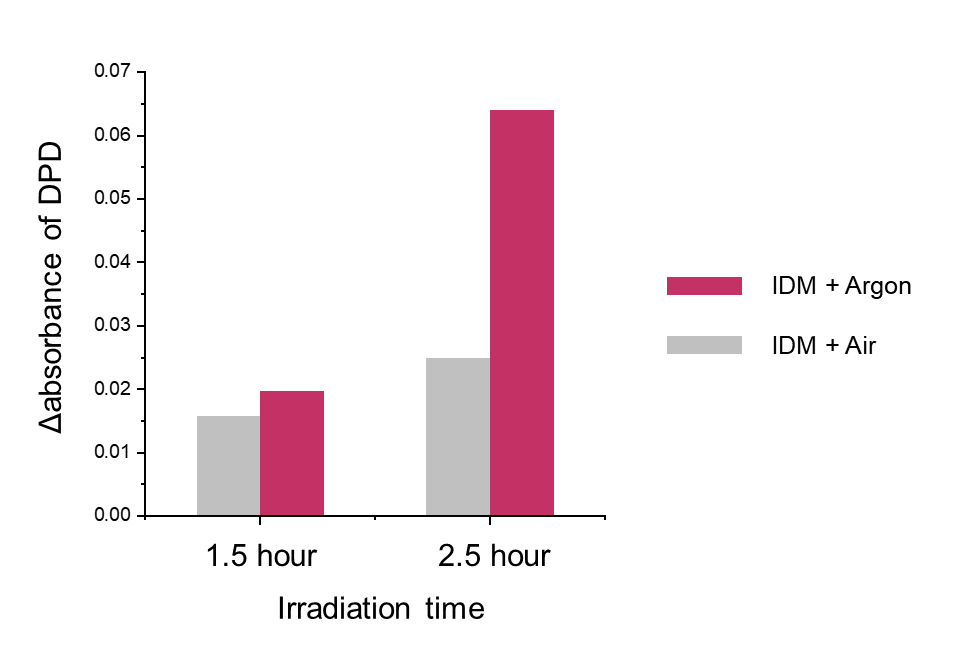

**Supplementary Fig. 4. Hydrogen peroxide generation by IDM photocatalysis.** DPD assay with horseradish peroxidase. IDM (50 μM) solution in normoxic (Air) and hypoxic (Ar-bubbled) conditions was irradiated by the green LED (λ_max_ = 525 nm, 66.7 mW·cm^‑2^) for 90 and 150 min. After photocatalysis, 200 µL of each sample was further incubated with 20 µg·mL^−1^ of peroxidase and 1 mM of DPD to measure a change in the absorbance of DPD.

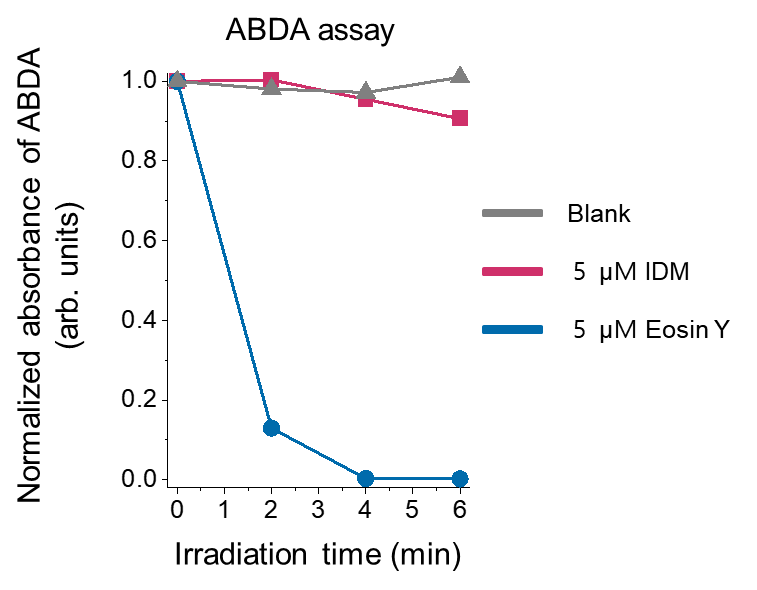

**Supplementary Fig. 5. Singlet oxygen generation assays.** ^1^O_2_ generation assay using ABDA absorbance decrease. The absorbance of ABDA (λ = 400 nm) was normalized to non-irradiated conditions, and the normalized ABDA absorbance was measured with irradiation time (green LED, λ_max_ = 525 nm, 16.6 mW·cm^−2^). The decrease in the absorbance of ABDA represents ^1^O_2_ generation.

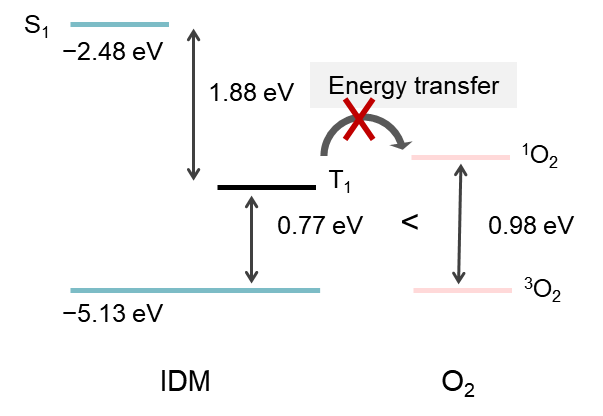

**Supplementary Fig. 6. Singlet and triplet energy diagrams.** Calculated singlet and triplet energy level diagram of IDM and energy transfer for singlet oxygen (1O2) generation by DFT calculation at the B3LYP/6-311G(d,p) level of theory with CPCM (water) model.

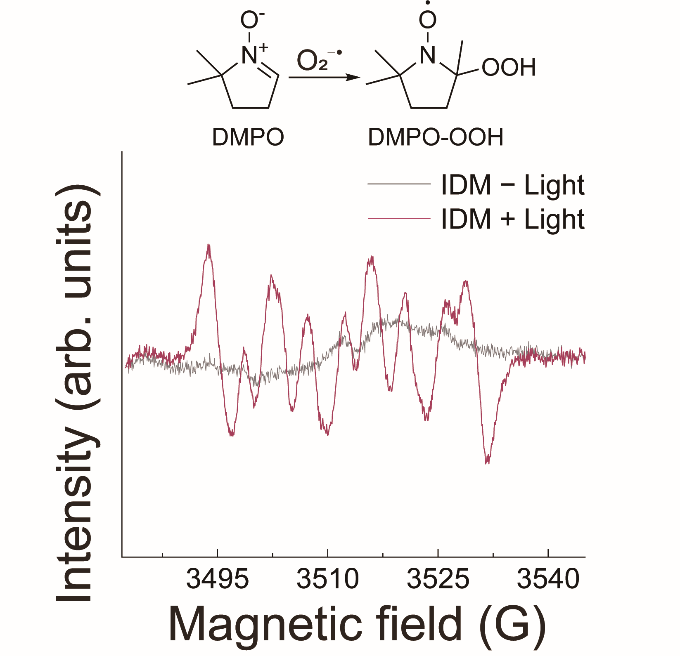

**Supplementary Fig. 7. Superoxide radical generation by IDM photocatalysis.** EPR spectra of O_2_^•−^ spin adducts (DMPO-OOH) after IDM photoactivation (green LED 1.0 J∙cm^−2^). Reaction conditions: [IDM] = 333 µM, [DMPO] = 167 mM.

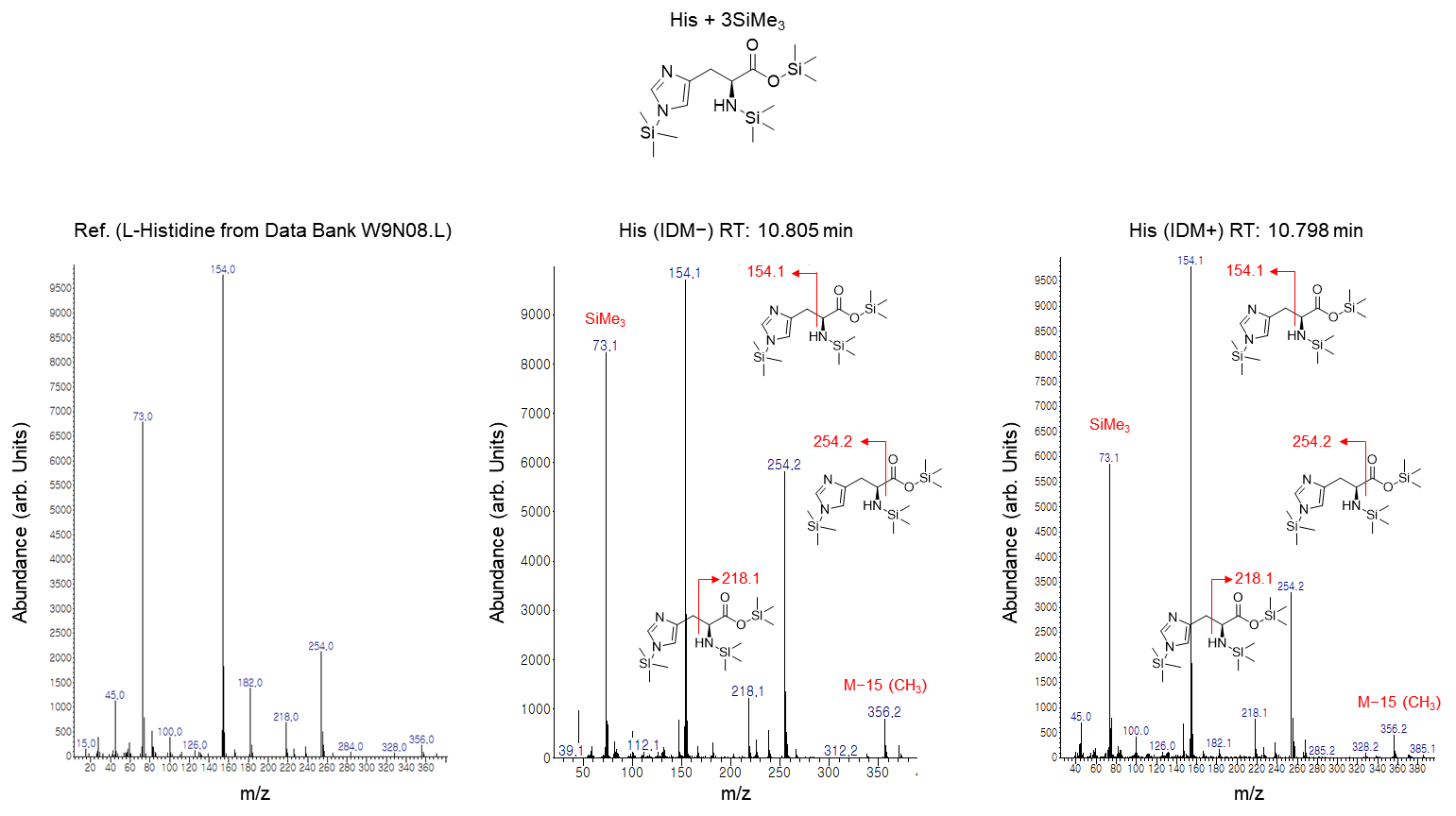

**Supplementary Fig. 8. GC-MS analysis for histidine oxidation by IDM photocatalysis.** GC-MS spectra of silylated histidine in Fig. 2e. The histidine peak was identified by reference spectrum from Data Bank Mass Spectra W9N08.L. Reaction conditions: [IDM] = 0.5 mM, [His] = 5 mM in 500 µL aqueous solution, green light irradiation: 120 J∙cm^−2^ (16.6 mW·cm^−2^ for 2 hours). After photocatalysis, dioxidized histidine was further silylated by MSTFA for gas chromatography.

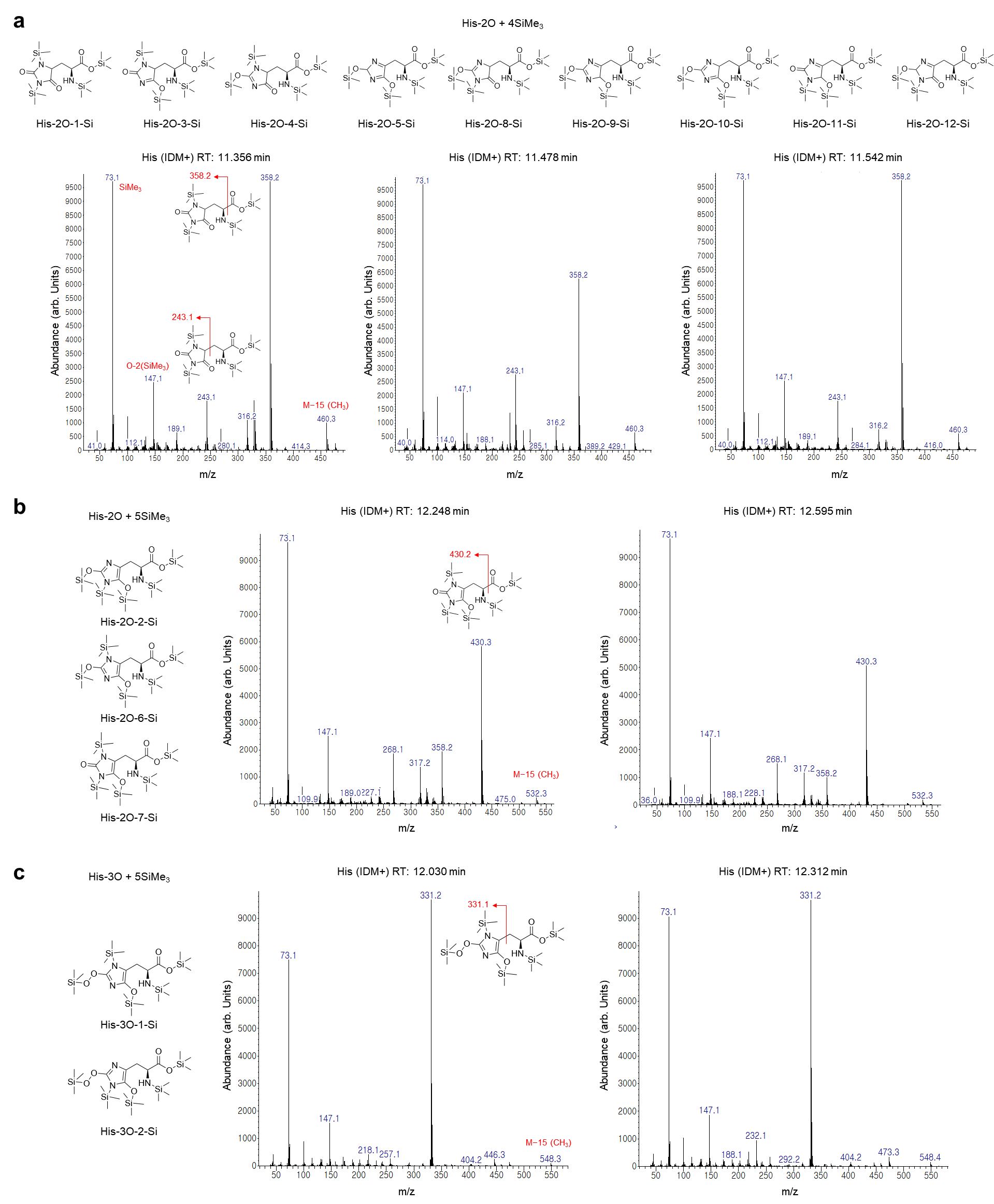

**Supplementary Fig. 9. GC-MS analysis for dioxidized and trioxidized histidine.** GC-MS spectra of silylated His-2O and His3O in Fig. 2e. **a** GC-MS spectra corresponding to His-2O with tetra-silylation. The three separated peaks of chromatogram after IDM photocatalysis (retention time: 11.356, 11.478, and 11.542 min) can be matched to 9 different His-2O tautomers. **b** Two peaks (retention time: 12.248 and 12.595 min) can be matched to 3 different His-2O tautomers with penta-silylation. **c** Two separated chromatogram peaks (retention time: 12.030 and 12.312 min) are matched to His-3O tautomers with penta-silylation. Reaction conditions: [IDM] = 0.5 mM, [His] = 5 mM in 500 µL aqueous solution, green light irradiation: 120 J∙cm^−2^ (16.6 mW·cm^−2^ for 2 hours). After photocatalysis, dioxidized histidine was further silylated by MSTFA for gas chromatography. Only one tautomer was described in the mass spectra, for example, which does not mean that the specific tautomers were assigned.

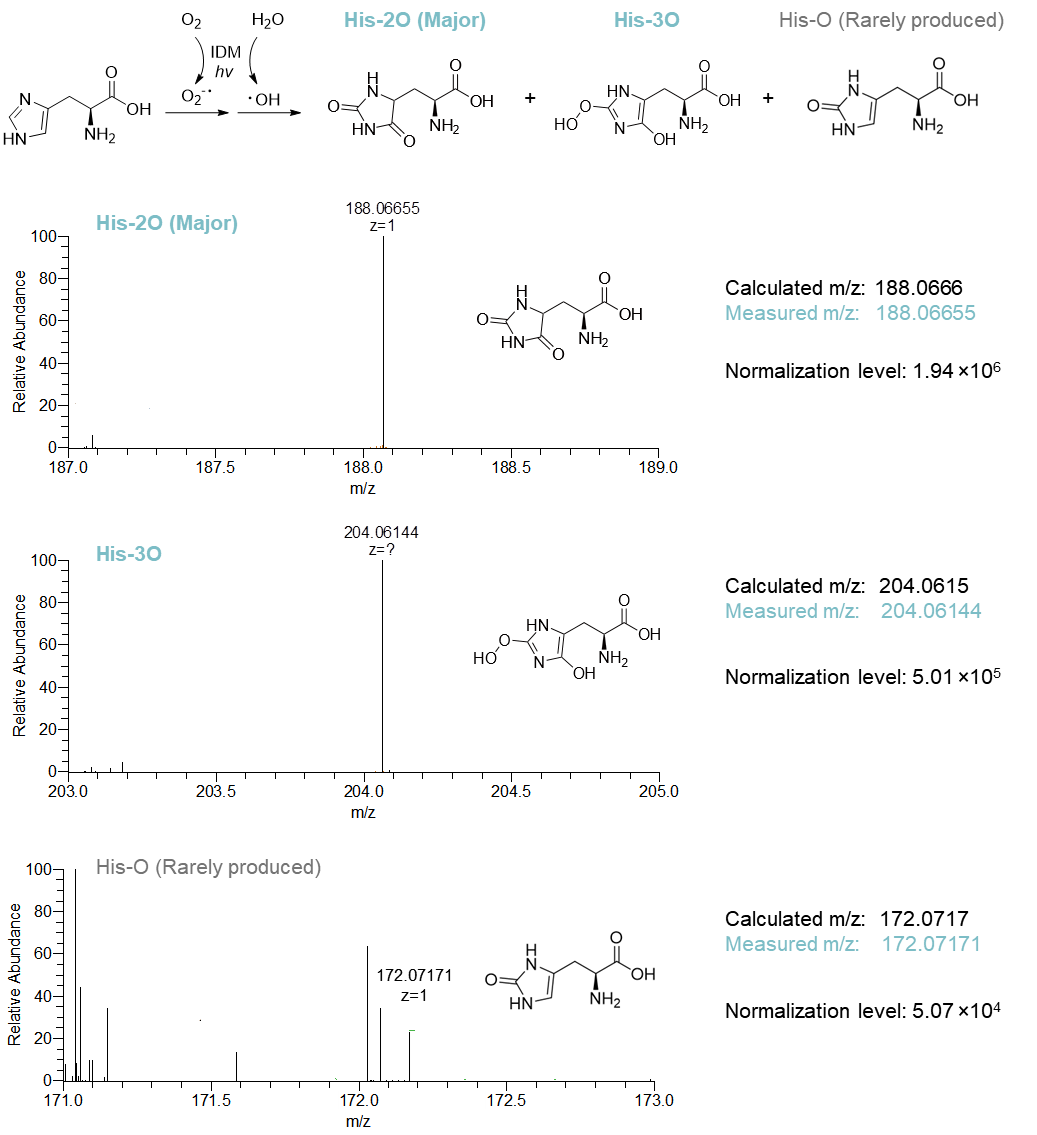

**Supplementary Fig. 10. HRMS analysis for histidine oxidation by IDM photocatalysis.** Product verification using HRMS. The reaction mixture of IDM photocatalysis with histidine was analyzed in positive mode (in MeOH with 1% formic acid). Scan ranges were defined by expected m/z values±2 of His-O (172.0717), His-2O (188.0666), and His-3O (204.0615). The relative abundance of each spectrum was normalized by the Normalization level. Reaction conditions: [IDM] = 0.5 mM, [His] = 5 mM in 500 µL aqueous solution, green light irradiation: 120 J∙cm^−2^ (16.6 mW·cm^−2^ for 2 hours).

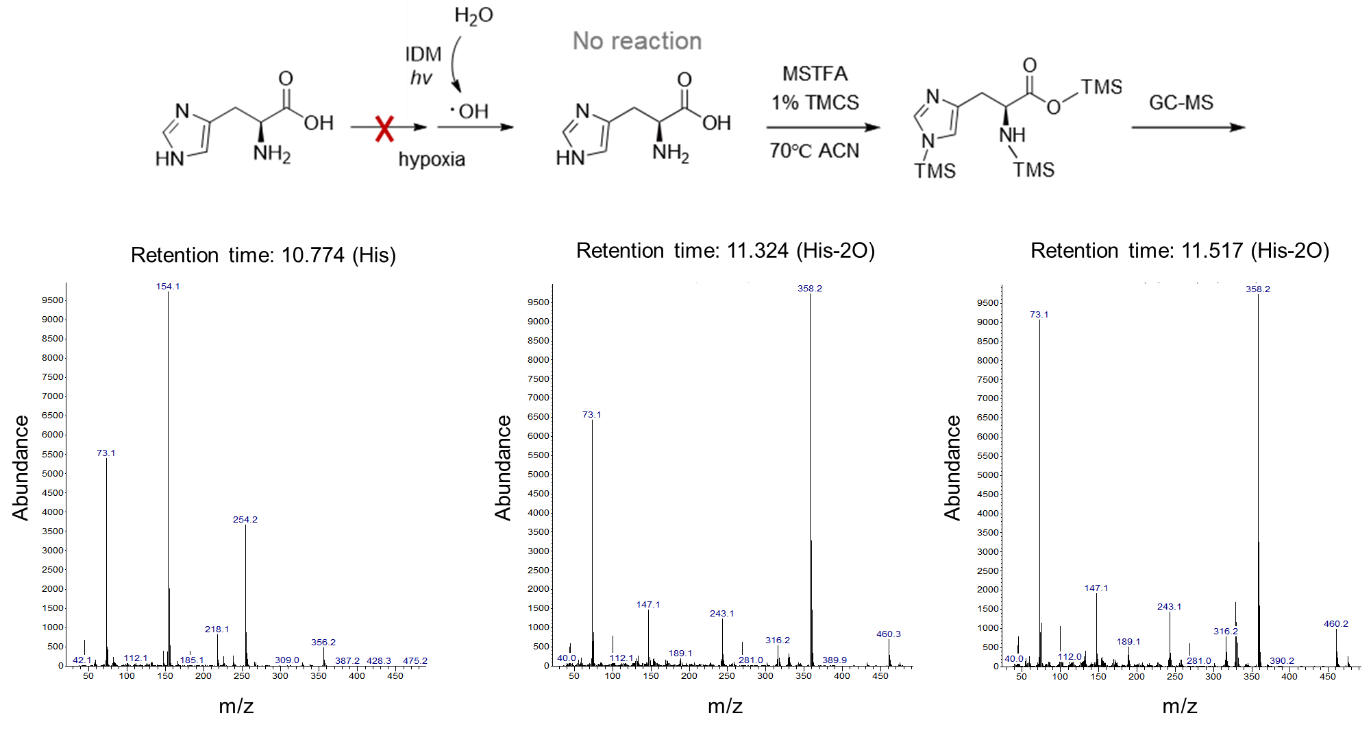

**Supplementary Fig. 11. GC-MS analysis for histidine oxidation in a hypoxic environment.** GC-MS spectra of silylated histidine in Fig. 3a. Reaction conditions: [IDM] = 0.5 mM, [His] = 5 mM in 500 µL aqueous solution (Argon bubbled for 30 min), green light irradiation: 120 J∙cm^−2^ (16.6 mW·cm^−2^ for 2 hours). After photocatalysis, dioxidized histidine was further silylated by MSTFA for gas chromatography.

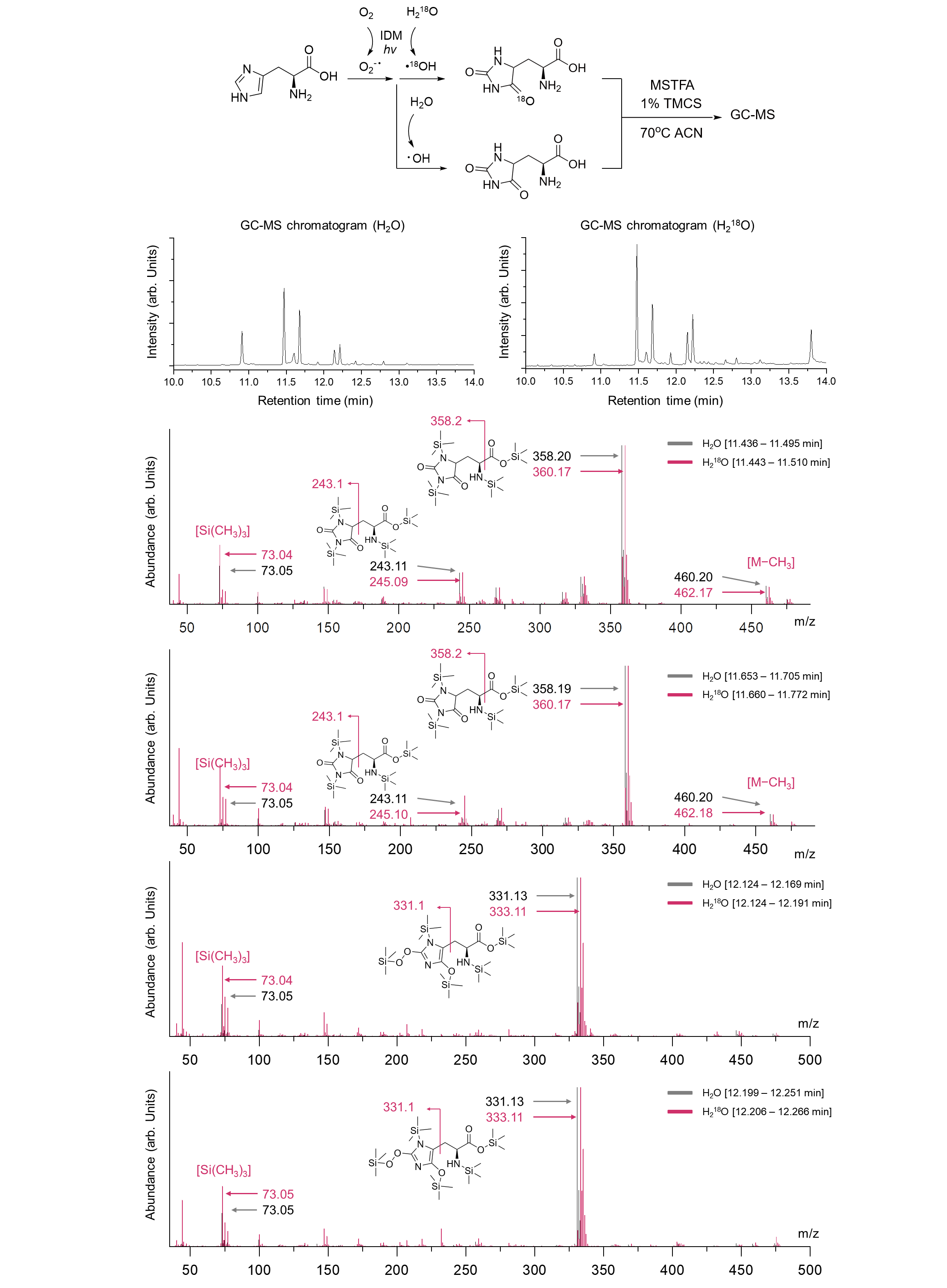

**Supplementary Fig. 12. GC-MS analysis for histidine oxidation in H_2_^18^O.** GC-MS chromatograms and spectra of silylated histidine in Fig. 3b. Reaction conditions: [IDM] = 0.5 mM, [His] = 5 mM in 500 µL aqueous solution (in H_2_O or H_2_^18^O), green light irradiation: 120 J∙cm^−2^ (16.6 mW·cm^−2^ for 2 hours). After photocatalysis, dioxidized histidine was further silylated by MSTFA for gas chromatography.

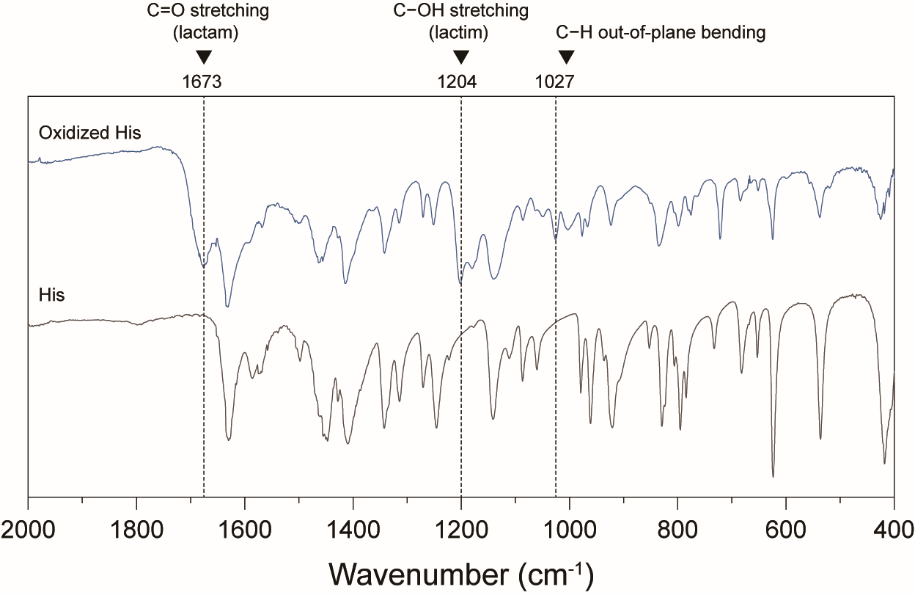

**Supplementary Fig. 13. FT-IR analysis of oxidized histidines.** FT-IR spectroscopic validation of histidine lactam-lactim tautomerization. FT-IR spectra of histidine before (His, bottom) and after (Oxidized His, top) IDM photocatalysis. Three new peaks emerged in oxidized histidine at 1673 cm⁻¹ (C=O stretching in lactam), 1204 cm⁻¹ (C–OH stretching in lactim), and 1027 cm⁻¹ (C–H out-of-plane bending due to loss of aromaticity), confirming lactam-lactim tautomerization and structural rearrangement. Oxidized histidine was HPLC-purified prior to FT-IR measurement.

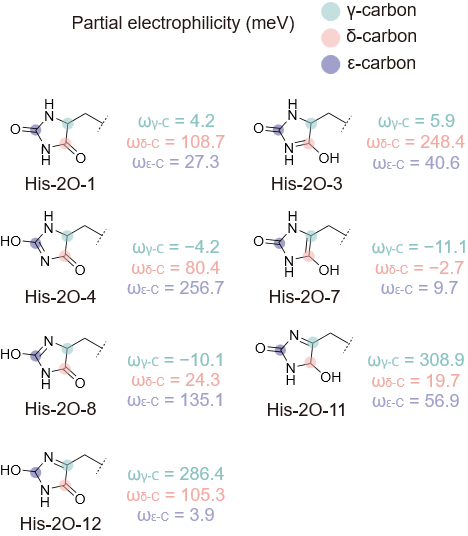

**Supplementary Fig. 14.** Partial electrophilicity of each carbon of His-2O tautomers. The highlighted atoms indicate γ-, δ-, and ε-carbon. The Fukui function was used for calculation.

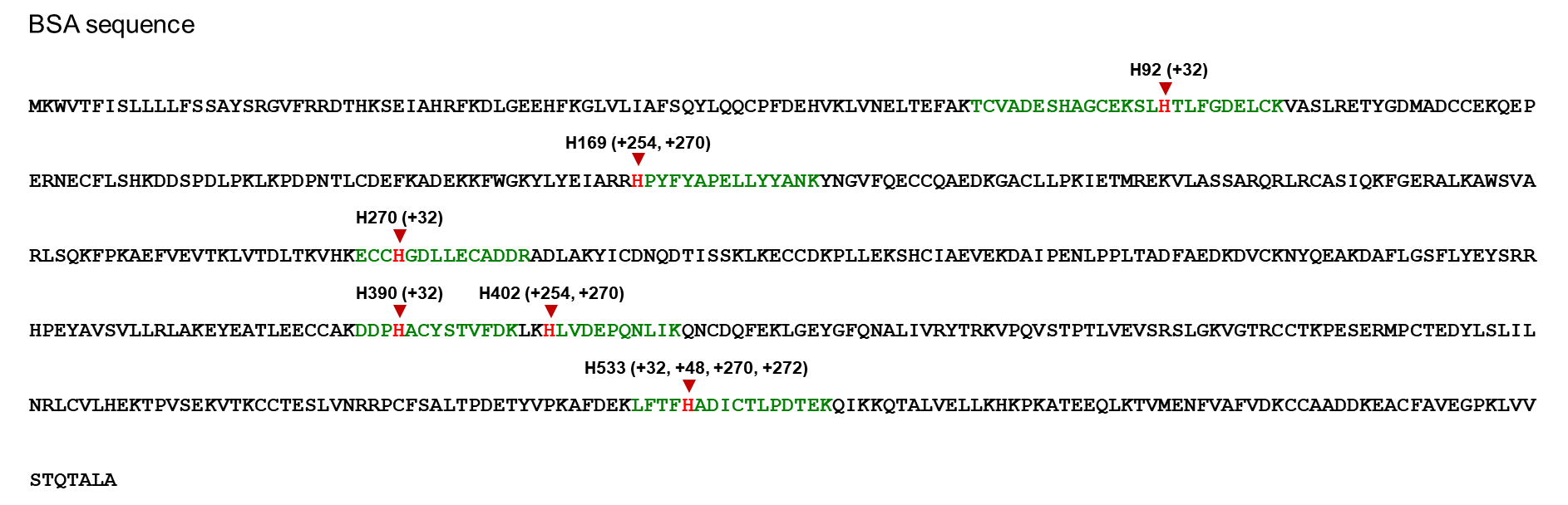
`

**Supplementary Fig. 15. Identified histidine biotinylation of BSA.** BSA amino acid sequence and identified histidine with di-/trioxidation (+32 and +48) and biotinylation (+254, +270, and +272). The identified histidine was marked red, and the corresponding peptides were marked green.

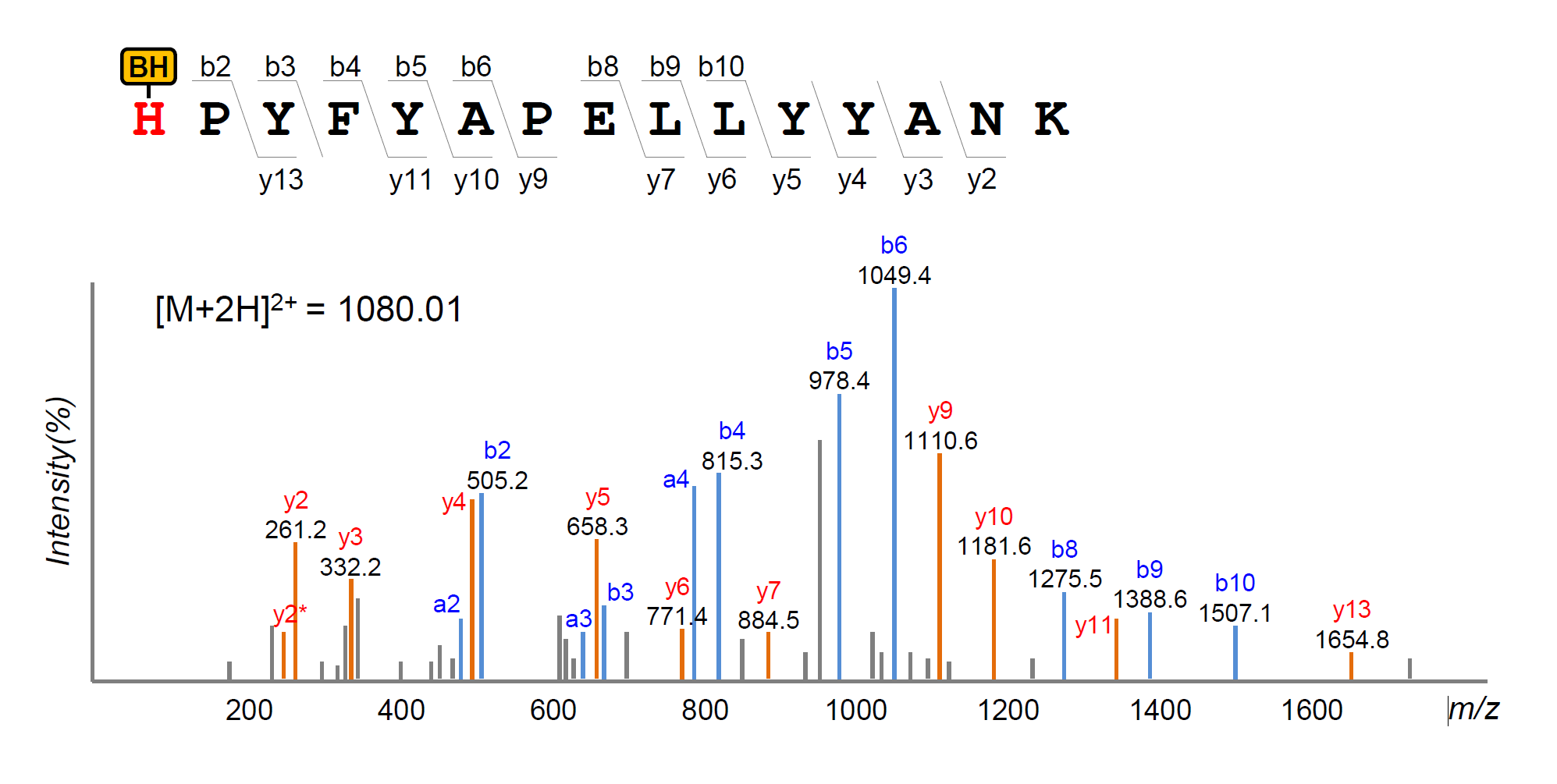

**Supplementary Fig. 16. MS/MS spectrum of biotinylated BSA.** Representative MS/MS spectrum of the BSA tryptic peptide containing a biotin-hydrazide modification (+270 Da) at His169. The observed b- and y-ion series are annotated, with b⁺ fragments highlighted in red.

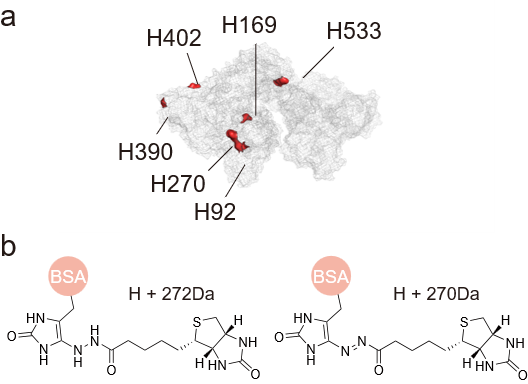

**Supplementary Fig. 17. Histidine labeling site of BSA and structures of biotinylated histidine. a** description of biotinylated (or dioxidized) histidine in BSA (PDB: 4F5S). **b** molecular structures for biotinylated histidine. The azo group's double bond can be reduced during the proteomics workflow.

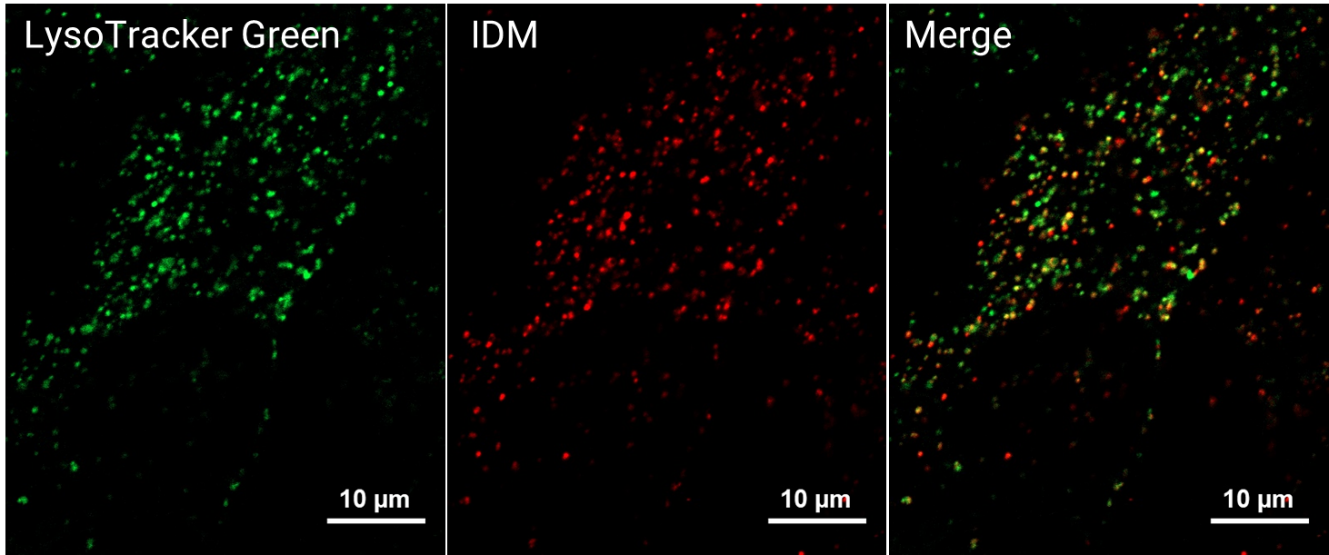

**Supplementary Fig. 18. Colocalization images with LysoTracker using confocal microscopy.** A549 cells were incubated with **IDM** (10 μM in media:DMF = 200:1, v/v) and LysoTracker Green (50 nM) for 24 hours. Then, the co-localization images were obtained in a CO2 incubator at 37 °C under a humidified atmosphere and 5% CO2 using a Carl Zeiss LSM980 with a 63X objective. (LysoG: λ_excitation_ = 488 nm, emission gain range: 500-530 nm; **IDM**: λ_excitation_ = 561 nm, emission gain range: 564-693 nm).

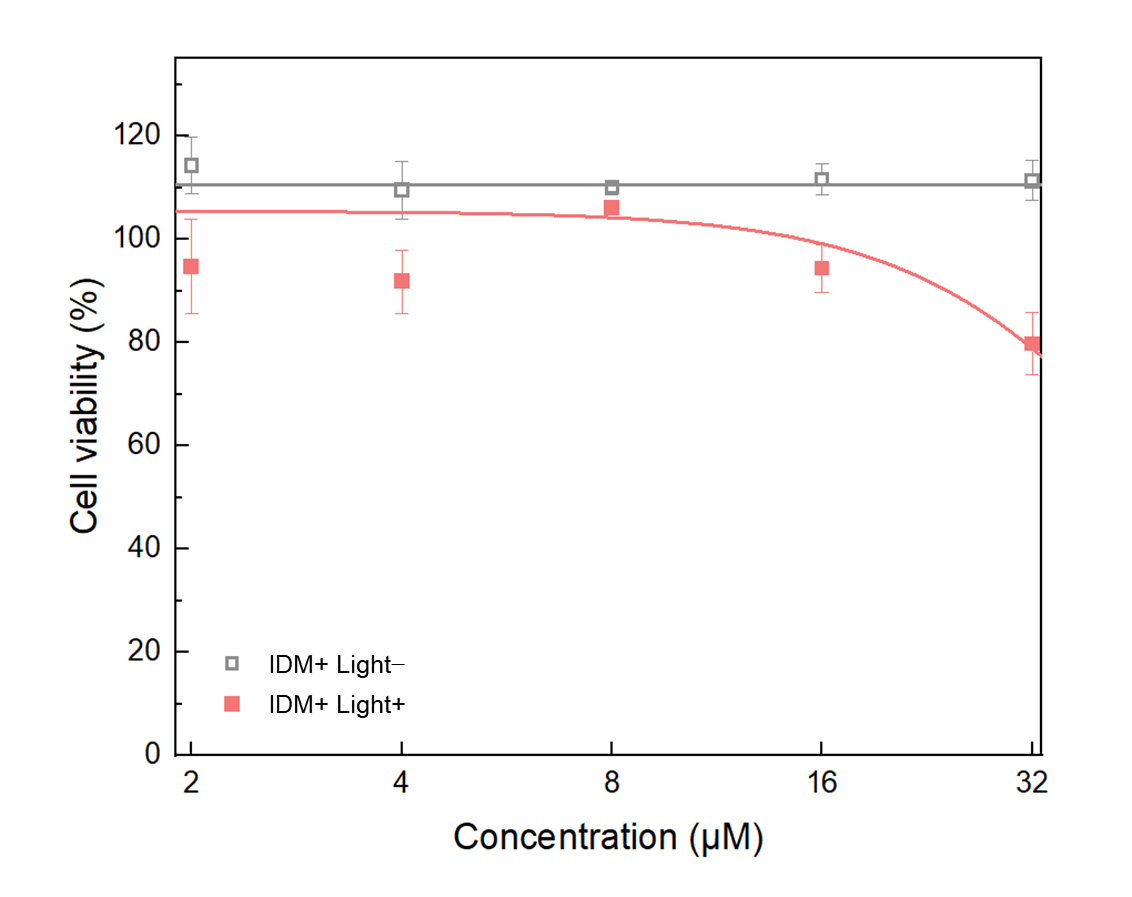

**Supplementary Fig. 19. Cell viability assessed by MTT assay.** HeLa cells were incubated with 2, 4, 8, 16, and 32 µM IDM for 2 hours. Green light was irradiated to IDM hv+ condition (λ_max_ = 525 nm; 30 J∙cm^−2^; 16.6 mW·cm^−2^ for 30 minutes), (*n* = 4).

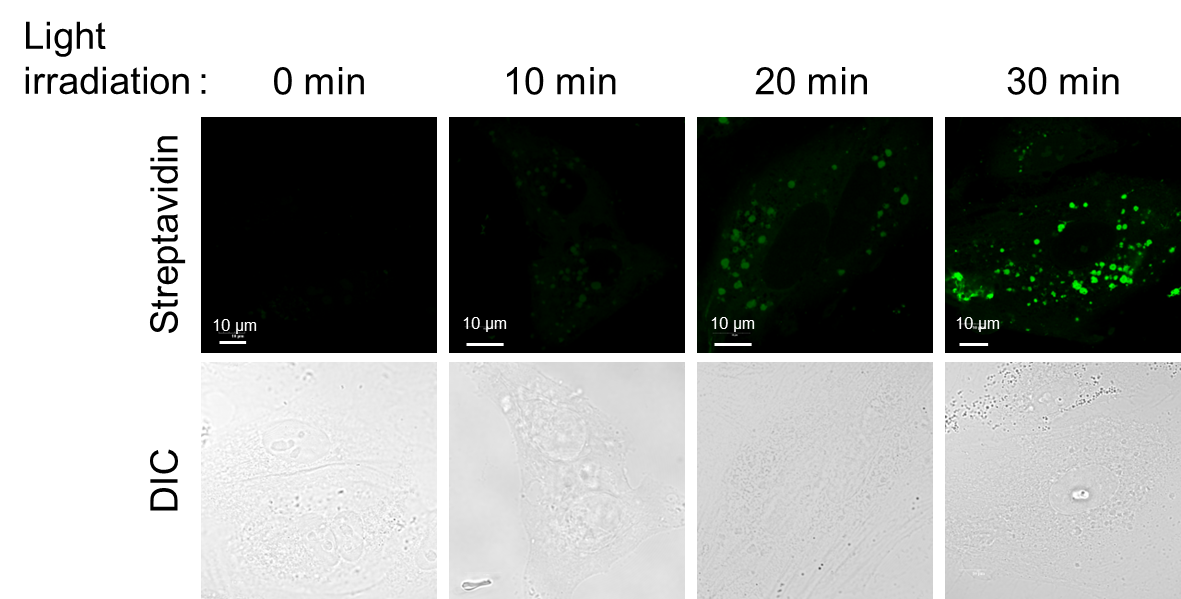

**Supplementary Fig. 20. Confocal images of intracellular biotinylated proteins.** A549 cells were pre-treated with 10 μM IDM in DMEM media overnight, followed by pre-incubation with biotin-hydrazide in DPBS (100 μM, 30 min). Cells were then exposed to green LED illumination (525 nm) for the indicated durations (0, 10, 20, or 30 min). After illumination, cells were washed, fixed, blocked, and stained with streptavidin–Alexa Fluor 488 to visualize biotinylated proteins. Confocal fluorescence images are shown for each illumination time point. Scale bar, 10 μm.

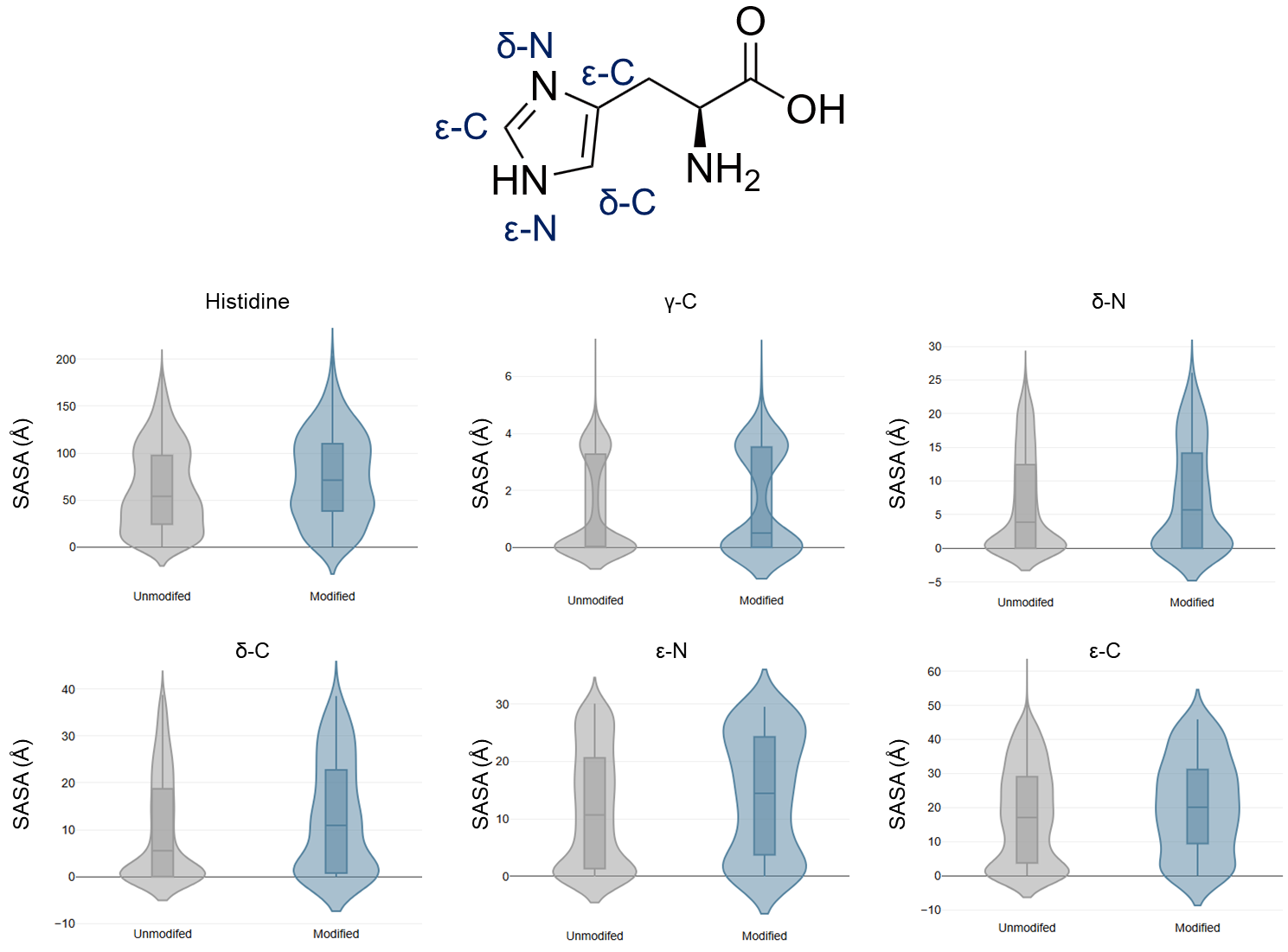

**Supplementary Fig. 21. Solvent-accessible surface area (SASA) analysis.** Solvent-accessible surface area (SASA) for 337 of PF-proteome’s AF2 structure model using a 1.4 Å water probe. SASA of every single atom in the imidazole ring of the histidine residue was analyzed to compare unmodified histidines and modified histidines. Only regions with high structural confidence (pLDDT ≥ 70) were included.
